# Haplotype-Resolved Genomics Reveals Conserved Chromatin Architecture and Epigenetic Constraints of Human Neocentromeres

**DOI:** 10.64898/2025.12.23.696241

**Authors:** Savannah J. Hoyt, Gabrielle A. Hartley, Mariah C. Antopia, Thomas W. Tullius, Dylan J. Taylor, Nicole M. Tillquist, Shane Neph, Nicole Pauloski, Katherine M. Munson, Kendra Hoekzema, Evan E. Eichler, Andrew B. Stergachis, Glennis A. Logsdon, Rachel J. O’Neill

## Abstract

Human neocentromeres are functional centromeres demarcated by CENP-A nucleosomes that form ectopically at alpha satellite-free loci. How neocentromeres reshape local chromatin and which features of native centromeric chromatin are preserved are unknown. We generated gapless, haplotype-resolved assemblies of native and neocentromeres from three patient-derived cell lines. Integrating CpG methylation, CENP-A profiling, and single-molecule chromatin fiber sequencing, we reveal chromatin features that define the essential centromeric architecture reconstituted during neocentromere establishment. We find that a deletion within the satellite array encompassing the hypo-CpG methylation centromere dip regions (CDRs) led to native centromere inactivation, that neocentromeres harbor CDRs and a dichromatin architecture, recapitulating features of alpha-satellite centromeres, and that LINEs demarcate neocentromere boundaries, implicating transposable elements in restricting CENP-A domain spreading. Moreover, neocentromeric chromatin is incompatible with promoter-like chromatin states, redefining the regulatory landscape within genic regions. Finally, using haplotype-specific chromatin footprinting, we resolve CENP-A nucleosome chromatin architecture of active centromeres.

## Introduction

Centromeres, the chromosomal sites of kinetochore formation and microtubule attachment, are essential for the proper segregation of chromosomes during both mitosis and meiosis^1^. Errors in centromere function can lead to chromosome loss, breakage, or structural rearrangements resulting in genomic instability^2,3^. In humans, a single functional centromere is located within alpha satellite (αSat) monomers organized into chromosome-specific higher-order repeat (HOR) arrays^4^. While some human chromosomes contain multiple αSat HOR arrays, only one array is active in most humans, demarcated by the centromere-specific histone H3 centromere protein A (CENP-A) which replaces the canonical histone H3 and epigenetically distinguishes centromeric from non-centromeric chromatin^5–7^. The first complete human genome sequence, T2T-CHM13^8^, provided an opportunity to study the sequence, structure, and epigenetic features that distinguish active endogenous centromeric chromatin domains across all human chromosomes^9–11^. These centromere assemblies revealed that within each active αSat HOR array lies a smaller region enriched with CENP-A nucleosomes coupled with a drop in CpG methylation frequency (centromere dip region, CDR) in dividing cells^9,11,12^. All native centromeres (NativeCens) in the T2T-CHM13 genome form on αSat HOR arrays ranging in size from ∼340kb (chr21) to 4.8Mb (chr18)^11^. Additionally, embedded transposable elements (TEs) mark shifts from regions of hypermethylation (αSat in the active HOR) to hypomethylation that correspond with transcriptional activity, establishing epigenetic boundaries surrounding active αSat HOR array domains that otherwise exhibit minimal transcriptional activity^10^.

Despite the ubiquity of αSat DNA in NativeCens, αSats are not required for centromere establishment^13^. Since the discovery of the first human neocentromere (NeoCen)^14^, a functional centromere at a non-canonical site, more than 100 NeoCen cases have been identified wherein CENP-A nucleosomes are assembled and the kinetochore forms in a region devoid of αSats^13–22^. Thus, NeoCens serve as satellite-independent models allowing for the decoupling of sequence-based epigenetic marks from the epigenetic framework that specifies centromeric chromatin. However, it remains unknown if other canonical epigenetic features of NativeCens are found in NeoCens or if NeoCen formation impacts local chromatin and gene transcription. Moreover, studies of patient-derived NeoCens and inducible NeoCens have not resolved each haplotype; rather, these studies have relied on a single consensus haplotype, preventing a clear delineation of the allelic differences between active and inactive NativeCens and the accompanying active NeoCen and its homologous, non-centromeric region.

Herein we focused on three commonly studied human patient-derived cell lines, each carrying a NeoCen on one homolog of a single chromosome pair: PDNC4 (on chr4), MS4221 (on chr8), and IMS13q (on chr13) (Fig. 1). In each cell line, the affected chromosome is epigenetically heterozygous at the centromere, with one homolog carrying an active centromere in a region on the chromosome arm lacking αSat (NeoCen) and the other homolog carrying an active centromere at the endogenous αSat-rich location (NativeCen). To assess the cause of NativeCen inactivation and genomic changes accompanying NeoCen formation elsewhere on the same homolog, we generated regional, haplotype-phased and gapless centromere assemblies spanning both alleles at the NativeCen and NeoCen locations for all three cell lines to define the genetic and epigenetic framework distinguishing non-centromeric regions (inactive NativeCen and the homologous region of the NeoCen locus without a centromere) from their centromeric counterparts (active NativeCen and NeoCen; Fig. 1). Further, we used CENP-A CUT&RUN, ChIP-seq, and DiMeLo-seq, single nucleotide-resolution CpG methylation status, long-read based chromatin accessibility (Fiber-seq^23^), RNA-seq (gene transcription), and Precision Run-On (PRO)-seq (sites of nascent transcription)^24^ to resolve allele-specific epigenetic and regulatory landscapes of NeoCens.

**Figure 1.**
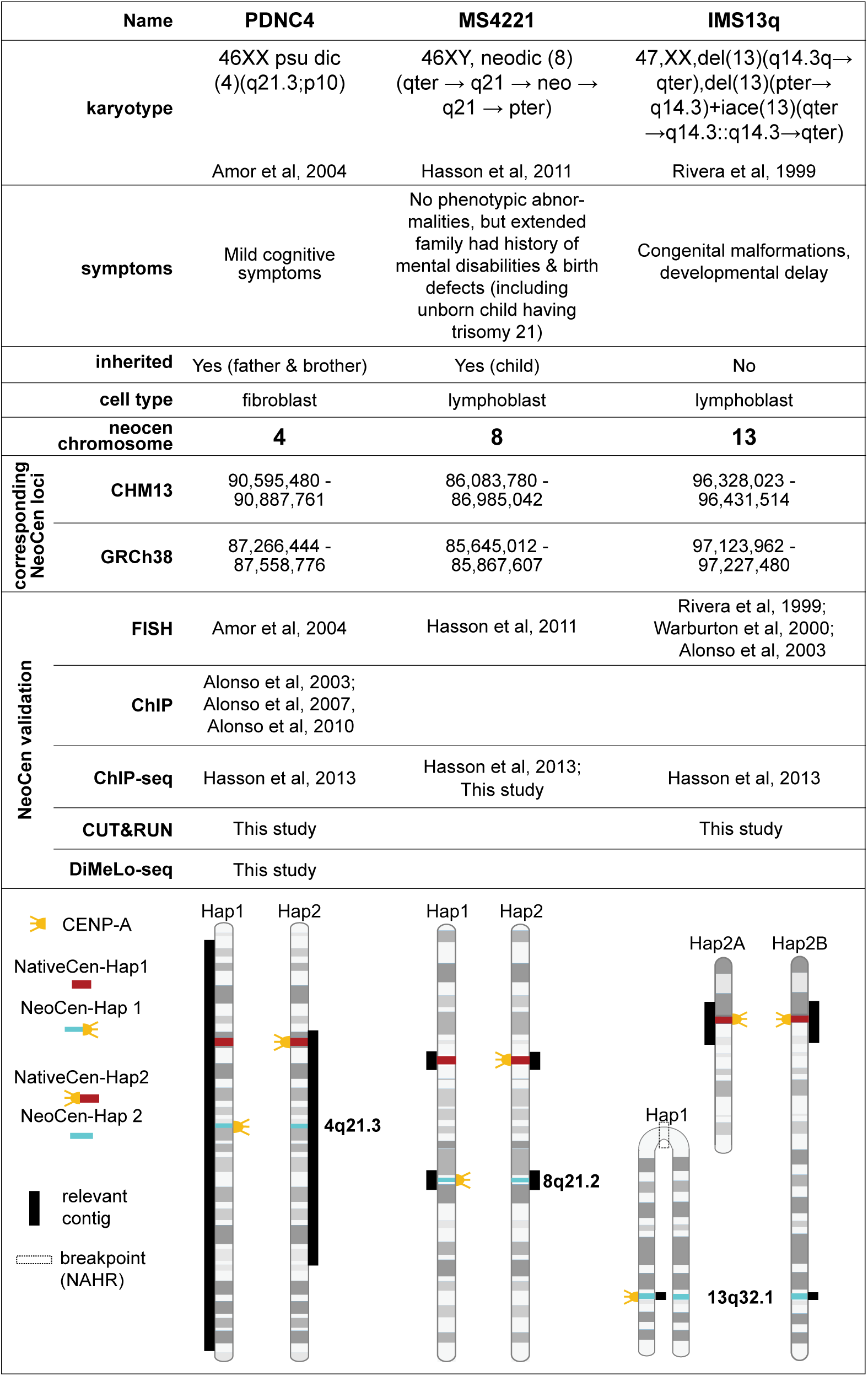
Three cell lines carrying alleles with neocentromeres linked to centromere silencing. Karyotype, phenotypic and genomic data associated with previous studies of PDNC4, MS4221, and IMS13q; ideogram of alleles of the affected chromosome for each line (bottom). Yellow: CENP-A assembly sites for native centromeres (red bars) and neocentromeres (blue bars). NAHR breakpoint is indicated (box) for the derivative chr13 carrying the active neocentromere in IMS13q. Cell-line specific phased, gapless contigs (black bars) are shown alongside respective chromosomes.

## Results

### Generation of contiguous and gapless assemblies for NativeCen and NeoCen loci

To generate haplotype-phased assemblies for NativeCens and NeoCens for PDNC4 (chr4), MS4221 (chr8), and IMS13q (chr13), we generated and assembled both PacBio HiFi (36-52X coverage) and Oxford Nanopore Technologies (ONT) sequencing data (55-107X coverage) (SuppTables 1,2). Each assembly was validated to confirm contiguity across the targeted regions (Methods, SuppTable 2, SuppFigs. 1-4). PDNC4 contains an inherited neodicentric chr4, with the active NeoCen in 4q21.3^25^ spanning a set of four protein-coding genes. MS4221 contains a heritable neodicentric chr8, with the active NeoCen in 8q21.2^22^ on a large VNTR (Variable Number of Tandem Repeats) array. IMS13q carries a NeoCen on chr13, formed via mitotic rescue of an inverted duplication (invdup) marker chromosome originating from a 13q14.3 break (SuppFig. 5) and lacking an active NativeCen^16,17^. Across all three lines, we designated the contigs carrying the active NeoCen and inactivated NativeCen as haplotype1 (Hap1) and the haplotype carrying the active NativeCen and canonical, non-centromeric NeoCen loci as haplotype 2 (Hap2) (Fig. 1). The NeoCen loci of Hap1were confirmed for each line (Methods). Using long-read sequencing, we refined the breakpoint in IMS13q from 13q21.1 (based on previous FISH studies^16,17^) to 13q14.3 and revealed mechanisms of post-breakage stabilization (Methods; SuppFig. 5). We found that NeoCen formation occurred at 13q32.1, and the NativeCen chromosome carrying the terminal-deletion was retained, containing 13pter-13q14.3 and resulting in two haplotypes for the active NativeCen (Hap2A and Hap2B). Like PDNC4, the IMS13q NeoCen-Hap1 lacks any predicted tandem repeat array and resides between two protein-coding genes^26^.

### IMS13q contains two highly similar, active chr13 NativeCens

Using both normalized ChIP-seq^27^ and CUT&RUN (Methods) to define CENP-A nucleosome occupancy across each haplotype of chr13 NativeCens (Hap2A and 2B) (SuppTable 3), we confirmed that both chr13 NativeCens are functionally active with near equal levels of CENP-A enrichment, one located on the 13pter-q14.3 (+13q34-qter) marker chromosome (Hap2A) and the second located on the unaffected chr13 (Hap2B). To determine CDR status of Hap2A and Hap2B, we derived ONT-based CpG methylation calls and observed a CDR on the proximal side of the αSat HOR of both haplotypes, yet each exhibits a unique CDR pattern (Fig. 2A-B, SuppFig. 6, SuppTable 4). Hap2B has a single CDR of uniform hypomethylation (104 kb), whereas Hap2A has two hypermethylation spikes within its CDR, creating three smaller CDRs (14kb, 34kb, 48kb, respectively), resembling CDRs in active chr13 NativeCens in T2T-CHM13^9^ and HG002^28^. Across both haplotypes, CENP-A enrichment is strongest within and around the CDRs (Fig. 2A-B, SuppFig 6). To determine whether these two haplotypes share a similar genetic structure, we annotated the NativeCens for all repeats and αSat, and performed pairwise dot plots, generating maximal exact matches (MEMs). We found minimal structural differences between Hap2A and Hap2B, with only a 125kb size difference in the αSat HOR array (Hap2A is larger) both haplotypes carry similar αSat suprachromosomal families, HOR monomers (reviewed in^4,29^), and HOR structural variants (SVs)^4,11,30^ (SuppFigs. 6-8, SuppTables 6,7). Thus, despite the similar genetic structure of these two active NativeCens, the two haplotypes carry different epigenetic signatures in CDRs and, to a lesser degree, in their corresponding CENP-A domains.

**Figure 2.**
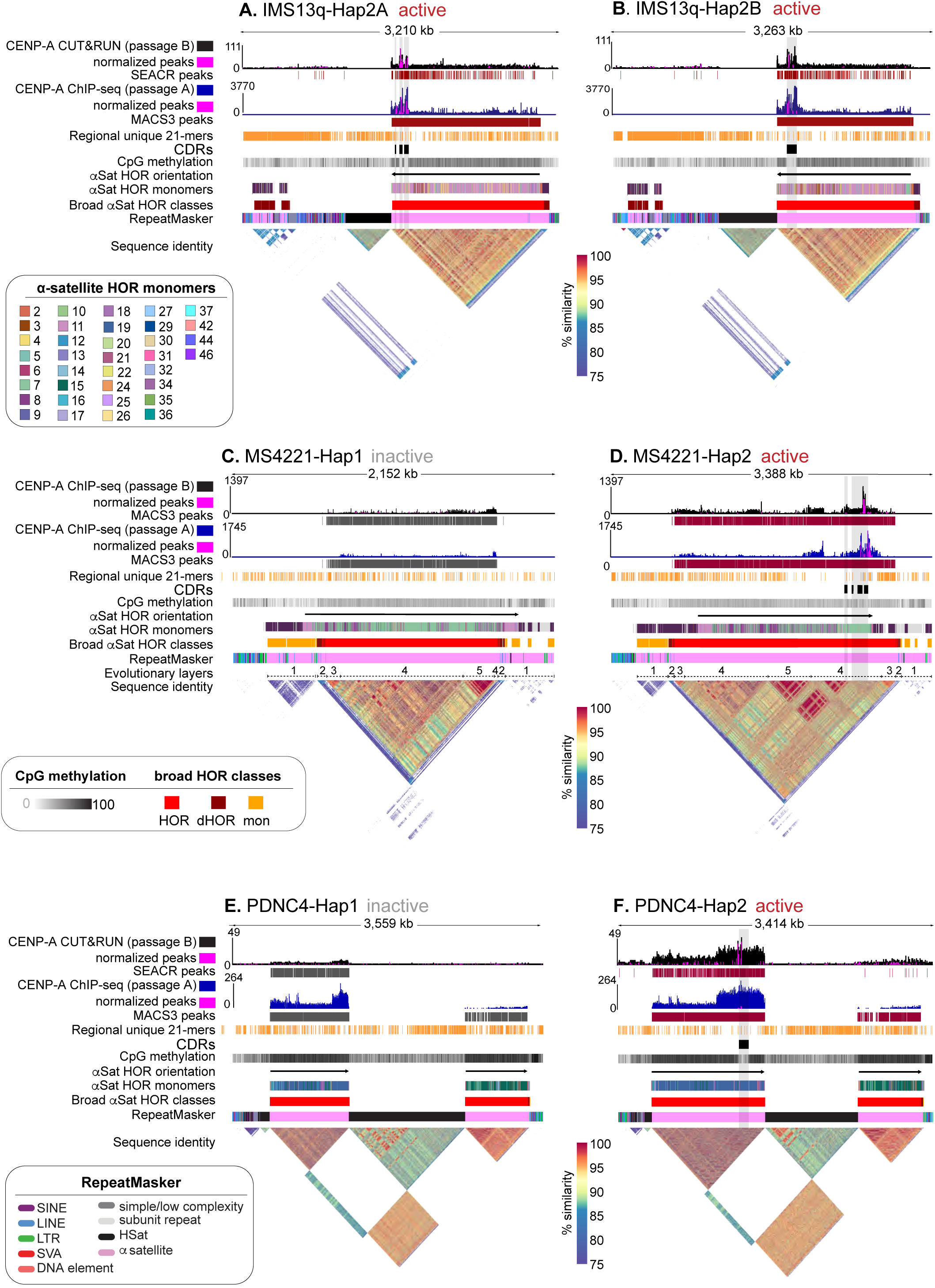
Epigenetic status of active and inactive NativeCen haplotypes. A comparison of sequence and epigenetic features of the NativeCen haplotypes across all three cell lines. **(A)** Hap2A(active) and **(B)** Hap2B(active) alleles of IMS13q, **(C)** Hap1(inactive) and **(D)** Hap2(active) alleles of MS4221, and **(E)** Hap1(inactive) and **(F)** Hap2(active) alleles of PDNC4. For each pair of haplotypes, tracks are indicated on the left. (Bottom) StainedGlass pairwise sequence identity heatmap across the NativeCen; gradient denotes color scale for % identity (shared between haplotypes) Broad αSat HOR classifications, k-mer number per HOR unit, and RepeatMasker annotations are color coded as per inset Keys. **(C-D)** include the evolutionary layer^12^ annotations.

### PDNC4 and MS4221 contain epigenetically heterozygous NativeCens caused by large SVs

In contrast to IMS13q, both PDNC4 and MS4221 are heterozygous for centromere activity at the NativeCens on chr4 and chr8, respectively, with only one haplotype (Hap2) of each chromosome carrying an active NativeCen, allowing for haplotype-specific analyses to test whether both epigenetic and sequence and/or structural changes accompany NativeCen inactivation. We compared NativeCen-Hap1 and -Hap2 within PDNC4(chr4) and MS4221(chr8) and found a CDR in the αSat HOR array of NativeCen-Hap2 yet absent in NativeCen-Hap1 (Fig. 2C-F, SuppFigs. 9-10, SuppTable 4). Using normalized CENP-A ChIP-seq/CUT&RUN data (Methods) to phase reads across haplotypes in both PDNC4(chr4) and MS4221(chr8), we find that CENP-A enrichment at the active NativeCen-Hap2 colocalizes with methylation-based CDRs, as expected (Fig. 2D-F, SuppFigs. 9B,10B). In addition, we performed long-read CENP-A DiMeLo-seq on PDNC4 to enhance haplotype phasing of CENP-A across the CDR, further supporting the observation that the active NativeCen on Hap2 carries a CDR that overlaps with CENP-A within the p-arm-proximal αSat HOR array (SuppFig. 10).

Given the previously observed reduction of αSat FISH signal in inactive Hap1 compared to active Hap2 for both PDNC4 and MS4221 (53%^25^ and 60%^22^, respectively), we reasoned that epigenetic differences between the NativeCen haplotypes may be caused by structural differences. To determine if there are any structural and/or sequence differences among active and inactive NativeCens, we generated both pairwise dot plots and αSat specific annotations for MS4221(chr8) and PDNC4(chr4)(Fig. 2C-F, SuppFigs. 9-14, SuppTables 5-8). We resolved the size difference within PDNC4 to 16% in the overall span of αSats and a 30% difference in the p-arm-proximal HOR (Hap2; SuppTable 5), and in MS4221 to a 48% difference in the overall span of αSat and a 64% difference in the HOR (Hap2 is larger; SuppTable 5). In PDNC4, both NativeCen haplotypes share a region of high sequence identity within the p-arm-proximal D4Z1 HOR, denoted by a dense concentration of MEMs, the boundaries of which correspond to the boundaries of CENP-A nucleosome occupancy found in the active NativeCen-Hap2. Despite sharing a region of αSat homogeneity within the D4Z1 HOR, the inactive NativeCen-Hap1 has a ∼30kb deletion within the D4Z1 HOR. This region corresponds to the region in the active NativeCen-Hap2 that carries the CDR, indicating a deletion in NativeCen-Hap1 led to the loss of this epigenetic mark (SuppFigs. 12B, 13A). In the active NativeCen-Hap2 of MS4221(chr8), there are two CENP-A ChIP-seq peaks, a major peak that coincides with a CDR and a minor peak with no underlying CDR (Fig. 2D, SuppFig. 9B). Both peaks share αSat sequence identity with CENP-A signal on older αSat expansions flanked by more recent ones, as has been observed on chr8 in CHM13^12^ and HG0073^12^, both of which also share similar HOR sizes, 2.08Mb-2.36Mb (SuppFigs. 11B, 14). In contrast, the inactive NativeCen-Hap1 carries only a tiny, low-density CENP-A peak (∼70kb) with no detectable CDR (Fig. 2C-D, SuppFigs. 11B). As observed in PDNC4, the inactive NativeCen-Hap1 carries a deletion (∼1.2Mb), encompassing the αSat layers that define the active centromere on Hap2 (Fig. 2C-D, SuppFigs. 9, 11, 14). In both cell lines, the deletion within the HOR of the CDR region is likely the primary cause of loss of function of the NativeCen and chromosome rescue by NeoCen formation. The detection of CENP-A by immunofluorescence at inactive NativeCens for both PDNC4 and MS4221, albeit at reduced levels^22,25^, coupled with our detection of CENP-A nucleosomes in our filtered ChIP-seq/CUT&RUN mapping data, suggests CENP-A continues to be incorporated at these inactive sites, but at a density too low to establish a CDR and a functional centromere.

### Failure of the Chr4 epiallele to rescue NativeCen inactivation

Each centromere in the human genome is classified by chromosome-specific HORs, each defined by the number of monomers per HOR unit. In CHM13, chr4 NativeCens carry a block of HSat1A that separates a single HOR of D4Z1 (predominantly 19 monomers (19-mer) per HOR unit) into two blocks of the same monomer-HOR structure^11^. While the second D4Z1 array is the site of the CDR/CENP-A domain in CHM13 and MS4221 (Fig. 3A, SuppFig. 15, SuppTable 5), other individuals in the human population carry a different structure, with a single D4Z1 array bounded by blocks of HSat and a second αSat array carrying a different monomer-HOR structure (15-mer) (Fig. 3B,C). Upon closer inspection, this 15-mer HOR array is found in both CHM13 and MS4221, but only as a small, distal array (Fig. 3A). However, we find that this 15-mer HOR is expanded in several human chr4s, including those in HG002^28^ and several individuals in the HPRC/HGSVC data (HG00733, HG0114, and HG02492)^31,32^ (Fig. 3C) and that the 15-mer HOR array carries a CDR in HG02492 and HG00733 and thus acts as a centromeric epiallele, which we named D4Z1B (Fig. 3B-C, SuppTable 5).

**Figure 3.**
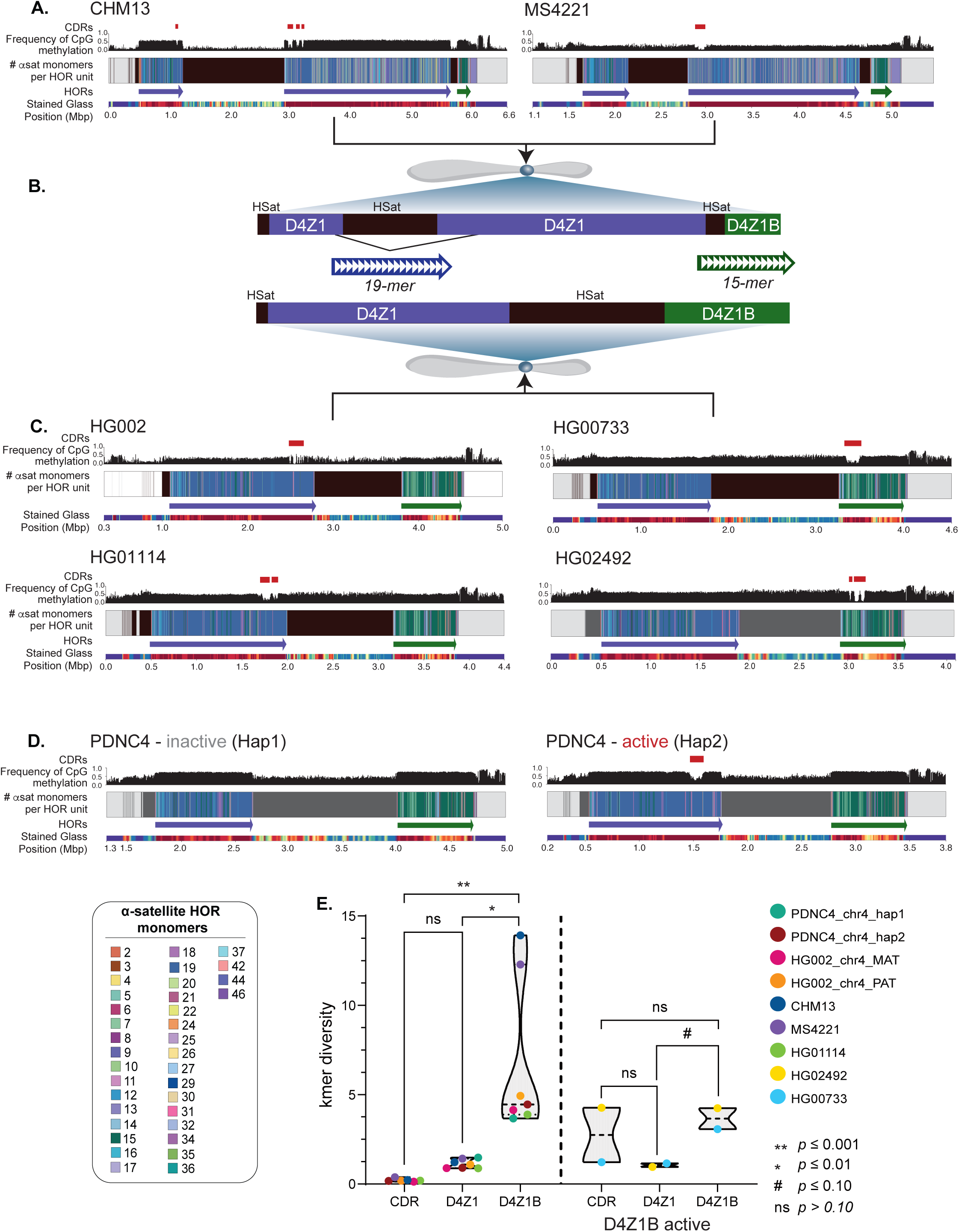
NativeCens on chr4 carry functional epialleles. **(A)** Chr4 NativeCens for a single haplotype of CHM13 and MS4221. **(B)** The relationship of **(A)** centromere structures to **(C)** centromere structures. Epialleles are indicated by name and color, with the dominant monomer number shown. **(C)** Chr4 NativeCens for a single haplotype of HG002, HG0114, HG00733, and HG02492. **(D)** Chr4 NativeCens for both haplotypes of PDNC4. Tracks for (A, C-D) are shown on the left (blue=D4Z1, green=D4Z1B). **(E)** Percent coverage of unique k-mers per allele across CDRs and D4Z1 and D4Z1B HORs. D4Z1-active alleles are shown on the left and D4Z1B alleles on the right. Statistical significance from Type I ANOVA are indicated.

Given the presence of differential epiallele activity for chr4 in the human population, we sought to understand why the loss of D4Z1 activity in PDNC4 NativeCen-Hap1 was not rescued by activation of the D4Z1B epiallele, rather than the formation of an active NeoCen at a distal locus ∼36Mb away (Fig. 2E, Fig. 3D, SuppFig. 10A). Previous studies have implicated αSat sequence identity as a driver of epiallele strength, wherein greater variation is associated with inactive epialleles^33,34^. D4Z1 in both PDNC4 NativeCen-Hap1 and -Hap2 are similar in their αSat organization and carry high MEMs, whereas D4Z1B in both NativeCens carry a much lower concentration of MEMs (SuppFig. 13), indicative of high sequence variation and high k-mer diversity. Across chr4s where D4Z1 is the active array, PDNC4-Hap2, CHM13, HG002, MS4221, and HG0114, the CDR contains markedly fewer unique 21-mers compared to D4Z1B and the inactive portion of D4Z1, reflecting minimal sequence variation and a high degree of αSat monomer similarity when D4Z1 is active (Fig. 3E). However, when D4Z1B is the active array––HG00733 and HG02492––k-mer diversity is higher in D4Z1B than D4Z1 (P=0.054; Fig. 3E), indicating sequence diversity is not a driving factor in epiallele activity. The failure of D4Z1B to rescue loss of the D4Z1-CDR in PDNC4-Hap2 indicates that the CENP-A domain and its accompanying CDR cannot spontaneously form on a different array, but rather, active D4Z1B arrays are inherited.

### CDRs are found at active NeoCens irrespective of underlying repeat content

While over 100 cases of NeoCen carriers have been documented in the human population, MS4221(chr8) NeoCen-Hap1 is the only neocentromere described to date that assembles CENP-A nucleosomes in a VNTR array. This VNTR, at 8q21.2, is a tandem array of a 12.2kb composite repeat unit that is human-specific, with each unit including LINEs, SINEs, DNA transposons, two composite subunits, and a REXO1L1 pseudogene^35^ (Fig. 4A-B inset). The copy number of composite units and their domain orientation^12^ within this VNTR varies in the human population from 26.5 (326kb) to 163 (1.985Mb) per haplotype. In MS4221, the chr8-VNTRs differ in size (805.9kb vs. 1.01MB) and number of composite units (66 across 5 domains vs. 83 across 11) between Hap1 and Hap2, respectively (Fig. 4A-B, SuppFig. 16, SuppTable 9).

**Figure 4.**
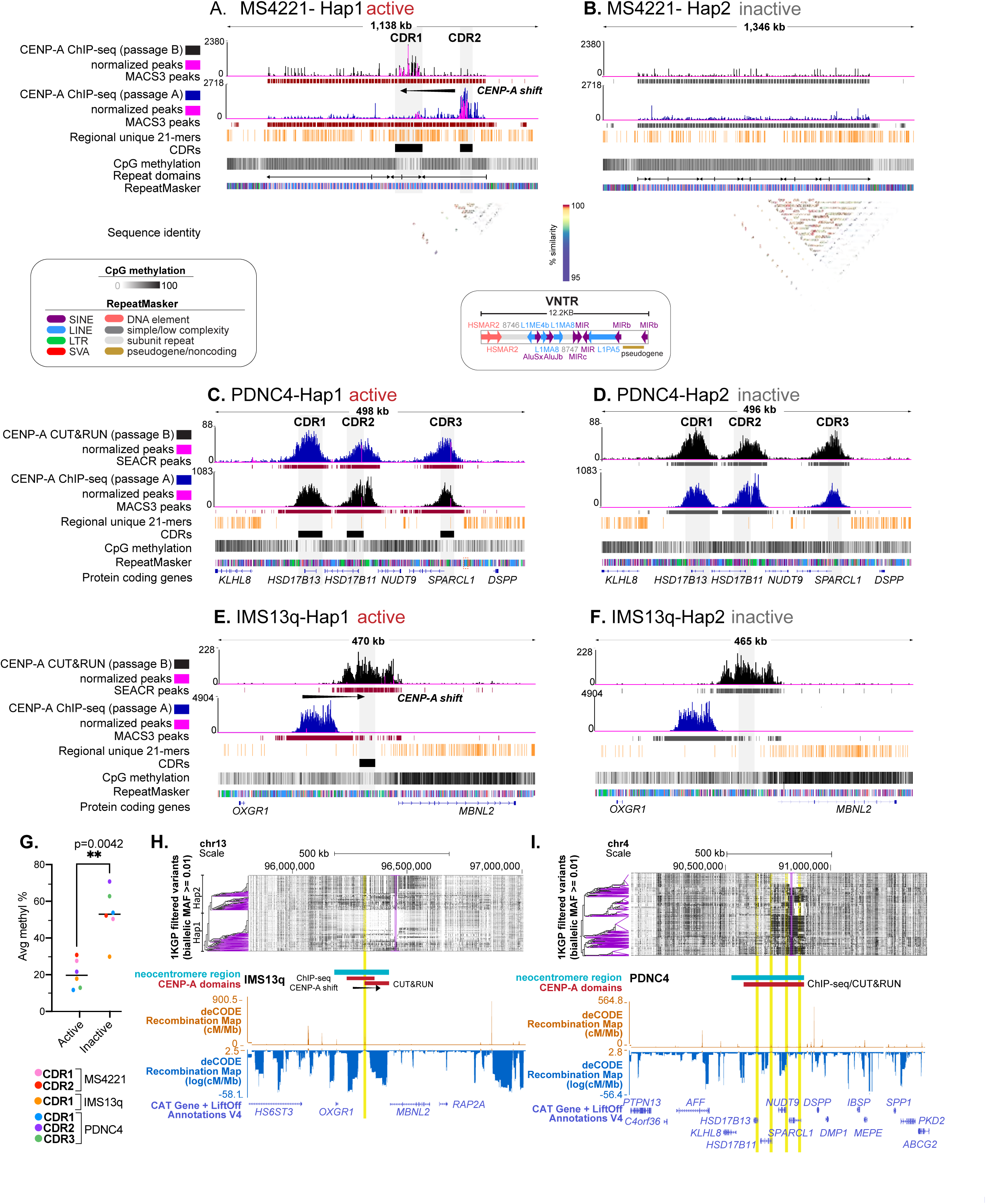
CDRs and low heterozygosity demarcate active NeoCens. **(A)** A comparison of sequence and epigenetic features of the active NeoCen haplotype and the non-NeoCen haplotype across all three cell lines. **(A)** Hap1(active) and **(B)** Hap2(inactive) alleles of MS4221, **(C)** Hap1(active) and **(D)** Hap2(inactive) alleles of PDNC4, and **(E)** Hap1(active) and Hap2(inactive) alleles of IMS13q. For each pair of haplotypes, tracks are indicated on the left. RepeatMasker annotations are color coded per repeat class in key. In **(A-B)**, StainedGlass pairwise sequence identity heatmap across each region is shown, with a gradient denotes color scale for % identity of alignements. Repeat domains for the VNTR (composite structure of the VNTR is shown) as per^12^. **(C-F)** Bottom track shows protein coding genes. **(G)** Average % CpG methylation among all active CDRs in Hap1 NeoCens and their inactive homologous regions on Hap2. **(H-I) (**Top) phased haplotypes from 1KGP for **(H)** IMS13q and **(I)** PDNC4 NeoCen loci (teal bars). Each row is a single haplotype where black represents the ALT allele and white represents the REF allele relative to T2T-CHM13. Vertical purple lines denote the variant used for clustering haplotypes. CENP-A domains are shown in red. (Bottom) Recombination rate tracks from deCODE are included as raw values (orange) (cM/Mb) and log values (blue) (log(cM/Mb), with recombination hotspots highlighted vertically across all tracks in yellow. Protein-coding genes are shown at the bottom.

Previously reported MS4221 CENP-A ChIP-seq data^27^ (Passage A) mapped to our assembly shows an increased signal intensity across a major and minor peak on the smaller VNTR array resolving the haplotype carrying the smaller VNTR as the active NeoCen-Hap1 (Fig. 4A-B). Both the major and minor CENP-A peaks correspond with CDRs; however, the minor CENP-A peak corresponds to a CDR that is larger (73.97%) and is slightly less (3%) methylated (Fig. 4A, SuppTable 4). In contrast, the same region of the unaffected Hap2 remains methylated across the VNTR with no detectable CDRs nor CENP-A peaks (Fig. 4B). Given the 10+ year age discrepancy between available CENP-A ChIP-seq data^27^ (Passage A) and the CpG methylation calls derived in our study, we generated CENP-A ChIP-seq data to match cell passage numbers used for our assembly (Passage B). The paired CENP-A ChIP-seq (Passage B) and CpG methylation data revealed that CENP-A enrichment in NeoCen-Hap1 has shifted 235kb upstream within the VNTR, aligning with the larger, more hypomethylated CDR coincident with diminishment of the previous CENP-A peak seen in Passage A (Fig. 4A). Thus, tandem arrays can support neocentromere activity in the absence of the canonical satellite size (∼171bp) found in human centromeres and shifts in CDR/CENP-A domains can occur in NeoCen.

In contrast to MS4221 NeoCen-Hap1, neither NeoCen location in PDNC4(chr4) nor IMS13q(chr13) contains tandemly arrayed sequences (Fig. 4C-F, SuppFig. 17A-B). In the PDNC4 NeoCen, previous ChIP-seq^27^ (Passage A), our CUT&RUN data (Passage B), and DiMeLo-seq (passage B) revealed three major peaks of CENP-A nucleosomes spread across a 292kb region carrying TEs, four protein-coding genes, *HSD17B13, HSD17B11, NUDT9, SPARCL1*, and an additional protein coding gene in each ∼100kb flanking region, *KLHL8, DSPP* (Fig. 4C-D, SuppFig. 17C, SuppTables 8, 10). We predicted these collective genomic features would form a more heterogeneous methylation pattern across the region rendering identification of a CDR challenging. However, we find that each of the three major CENP-A peaks corresponds with a CDR in only one haplotype, the active NeoCen-Hap1 (Fig. 4C, SuppTable 4). Thus, NeoCen CDRs are established independent of tandemly arrayed repeats.

As observed in MS4221, our CENP-A CUT&RUN data (Passage B) and CpG methylation calls for IMS13q revealed that CENP-A position has shifted ∼70Kb downstream from the peaks defined by older CENP-A ChIP-seq data^27^ (Passage A) mapped to our assembly, yet the domain size remains similar at ∼100kb (Fig. 4E-F, SuppTable 8). While the Passage A CENP-A domain resided between two protein coding genes (*OXGR1, MBNL2*), the Passage B CENP-A domain encompasses the promoter of *MBNL2,* which carries a hypomethylation dip in both haplotypes (Fig. 4E-F, SuppTable 10). IMS13q carries three copies of the 13q14.3-qter region (based on breakpoint refinement, Methods) and the NeoCen locus (13q32.1) contained within (Fig. 1, SuppFig. 5), with one haplotype harboring the active NeoCen (Hap1) and two haplotypes without any NeoCen activity (Hap1(invdup) and Hap2B). However, due to technological limitations in assembly of >2N ploidy (see Methods), our Verkko assembly only includes two phased haplotypes, meaning two of the three haplotypes are merged into one. To determine which two haplotypes were merged, we relied on the ONT CpG methylation data and the prediction that the NeoCen-Hap1 would harbor a CDR, as found in active NeoCens of both MS4221 and PDNC4. We assessed the methylation patterns from individual reads mapped to each haplotype (Methods), revealing that the two non-NeoCen homologous regions (Hap1(invdup) and 2B) were merged, resulting in a single, fully resolved and un-collapsed active NeoCen-Hap1 (SuppFig. 17D-E).

A comparison of average methylation frequency (ONT) across the CDRs of NeoCen-Hap1 and their corresponding homologous loci (Hap2) for IMS13q, PDNC4 and MS4221 revealed a reduction in methylation in NeoCen-Hap1 in each line (SuppFig. 18A, SuppTable 4). When methylation frequencies are clustered by NeoCen activity (Hap1 vs Hap2), we see a significant difference in methylation frequencies in an unpaired t-test (p-value = 0.0042) with a drop in overall methylation within the CDR of the active NeoCen compared to the unaffected homologous regions across all lines (Fig. 4G, SuppTable 4). To increase the resolution of the analysis, each CDR was partitioned into repetitive and non-repetitive sequences, and the average methylation was calculated. In all cases, significant differences in methylation levels were observed between CDRs in Hap1 and their inactive Hap2 counterparts (p≤0.01) (SuppFig. 18B). While methylation differences between the CENP-A domains of Hap1 and their homologous regions on Hap2 remained significant, no consistent differences were observed in the ± 100kb flanking regions that distinguished NeoCen-Hap1 and Hap2 in any cell line (Methods, SuppTable 8, SuppFig. 18C), suggesting that the observed methylation differences are specific to the establishment of an active NeoCen on a single allele.

### Low heterozygosity demarcates NeoCens of PDNC4 and IMS13q

Active NativeCens show high sequence homogenization, and thus low heterozygosity, across αSat/HORs, particularly within the CDR/CENP-A domain^11^. In pairwise-sequence comparisons, we uncovered a surprisingly high level of sequence identity matches and a low number of unique 21-mers and SNPs between each NeoCen-Hap1 and Hap2 pair in both PDNC4(chr4) (0.36 per kb) and IMS13q(chr13) (0.33 per kb), particularly within the CENP-A domain, mimicking the high sequence identity across αSats found in the active domains of NativeCen haplotypes in humans (SuppFig. 19A-B, SuppTable 11). Given that these two NeoCen-Hap1/Hap2 regions do not contain any repeat arrays, we wondered if low heterozygosity is linked to NeoCen formation in these cell lines. We reasoned that either low heterozygosity in a specific region may provide a favorable substrate for neocentromere formation or that low heterozygosity may be the result of gene conversion between alleles, as has been observed at active NativeCens. The former scenario would predict the low heterozygosity at NeoCen-Hap1/2 loci are found in the human population, while the latter would predict the low heterozygosity is specific to cell lines carrying active NeoCen loci. To test for either scenario, we generated phased variant calls against T2T-CHM13 using the 1000 Genomes Project (1KGP) data (*n*=2504) for the IMS13q(chr13) and PDNC4(chr4) NeoCen-Hap1/Hap2 loci (Fig. 4H-I). At the IMS13q(chr13) NeoCen-Hap1/Hap2 locus, we detected two common haplotypes spanning ∼250kb, suggesting a low recombination rate and thus low heterozygosity in this region, further supported by deCODE recombination maps (Fig. 4H). A single exception is a recombination hotspot in the middle of this NeoCen-Hap1/Hap2 locus, which falls within the Passage A CENP-A ChIP-seq^27^ peak (Fig. 4H). The CDR/CENP-A domain in Passage B has shifted such that the recombination hotspot borders the CENP-A peak, with *MBNL2* on the opposing side. In contrast, we did not observe any dominating haplotypes across the PDNC4 (chr4) NeoCen-Hap1/Hap2 loci, nor did we observe low recombination as several hotspots fall within the CENP-A domain (Fig. 4I).

Next, we calculated the number of heterozygous (het) variants per 1KGP individual in 10kb windows across each chromosome. We found significantly lower mean het counts in the IMS13q(chr13) NeoCen-Hap1/Hap2 locus relative to the rest of the chromosome (p=0.0018, one-tailed Mann-Whitney U test), but not in the PDNC4(chr4) NeoCen-Hap1/Hap2 locus, where the mean het count was higher (SuppFig. 19C-D). Lastly, we performed a Hardy-Weinberg Equilibrium (HWE) analysis using the same 1KGP data to determine if the number of observed heterozygous sites within each 1KGP population was lower than expected. This analysis was only possible at the chr13 locus due to the presence of two clear common haplotypes and SNPs that distinguish them – attributes not found at the chr4 locus (Fig. 4H-I). The expected number of heterozygous samples under HWE was calculated for two haplotype-distinguishing SNPs based on their allelic frequencies per 1KGP population. In comparison to the observed number of heterozygous samples, we found no significant deviation from the expected in any population (p>0.05), indicating that the IMS13q(chr13) NeoCen-Hap1/Hap2 locus falls within normal levels of heterozygosity with no apparent selective pressure (SuppFig. 19E, SuppTable 12).

Collectively, these data demonstrate that while the IMS13q(chr13) NeoCen-Hap1/Hap2 locus may be a site of low heterozygosity in the population, it is characterized by low recombination, a feature also associated with active NativeCens. In contrast, the low heterozygosity found in the PDNC4(chr4) NeoCen-Hap1/Hap2 locus is not reflected in any of our population-based analyses, suggesting the variant pattern seen in PDNC4 is likely cell line and/or patient specific. We propose that the observed low heterozygosity of both NeoCen-Hap1/Hap2 loci may be a consequence of gene conversion after NeoCen formation, a phenomenon known to occur at high frequency in active centromeres (reviewed in). However, without sequence-resolution of the progenitor chromosomal loci, we cannot determine whether low heterozygosity precedes or is a consequence of NeoCen formation; regardless, these data implicate a preference for CENP-A assembly in low heterozygosity regions.

### L1s serve as centromeric boundaries to CENP-A spreading in NeoCens

A commonality observed across NeoCens is the vast difference in repeat content compared to their endogenous, native centromeric counterparts. The CENP-A domains of all three NeoCen-Hap1/Hap2 loci vary in their overall repeat content (MS4221: 75%, PDNC4: 60%, IMS13q: 30%), with each lower than the near 100% repeat content of the αSat HOR arrays found in endogenous human centromeres (SuppFig. 20A). Similarly, CENP-B boxes, 17-bp motifs that denote the site of CENP-B binding, are highly enriched in the HOR arrays (regardless of centromeric activity; the exception being chrY) yet are largely absent in the NeoCen-Hap1/Hap2 loci, consistent with previous studies (SuppFig. 20B). However, we find that all centromeres in this study (Hap1 and Hap2 of both NativeCens and NeoCens) are AT-rich (59-69% and 52-61%, respectively; SuppFig. 20C, SuppTable 8), consistent with that observed for αSat.

We then asked if there was a difference in the repeat class distribution of the NeoCen-Hap1/Hap2 loci compared to flanking regions (+100kb on each side of the CENP-A domain). We find that in both PDNC4 and IMS13q NeoCen-Hap1/Hap2 loci, LINEs are more common in the flanks than in the CENP-A domains, mirroring that observed in endogenous centromeres^10,11^ (SuppFig 20D-F). Moreover, when comparing the IMS13q(chr13) active NeoCen-Hap1 CENP-A domains between Passage A and Passage B of this cell line, we find that the CENP-A domain not only shifted to a region of lower overall repeat content (42% to 33%; SuppFig. 20D), but to a region with lower LINE density (SuppTable 13).

LINEs, specifically L1s, have been previously linked to NeoCen activity^15,36^. Moreover, in the context of NativeCens, our earlier work showed that LINEs were found to be more enriched in centromeric and pericentromeric regions compared to other TEs, with L1s being the predominant TE insertion in αSat in CHM13^10,11^. L1s were found to form a gradient of sequence divergence traversing along the different layers of αSat divergence, with young L1s found in young αSat (HORs), and older L1s found in older αSat (dHORs, monomeric periphery)^11^. We find that there is a mix of LINE subfamilies and relative age groups (Methods) across NeoCen-Hap1 CENP-A domains, NeoCen-Hap2, and NeoCen-Hap1/Hap2 flanking regions (Fig. 5A-C, SuppFig. 20G), indicating that spontaneous neocentromere formation does not attract L1Hs transposition events.

**Figure 5.**
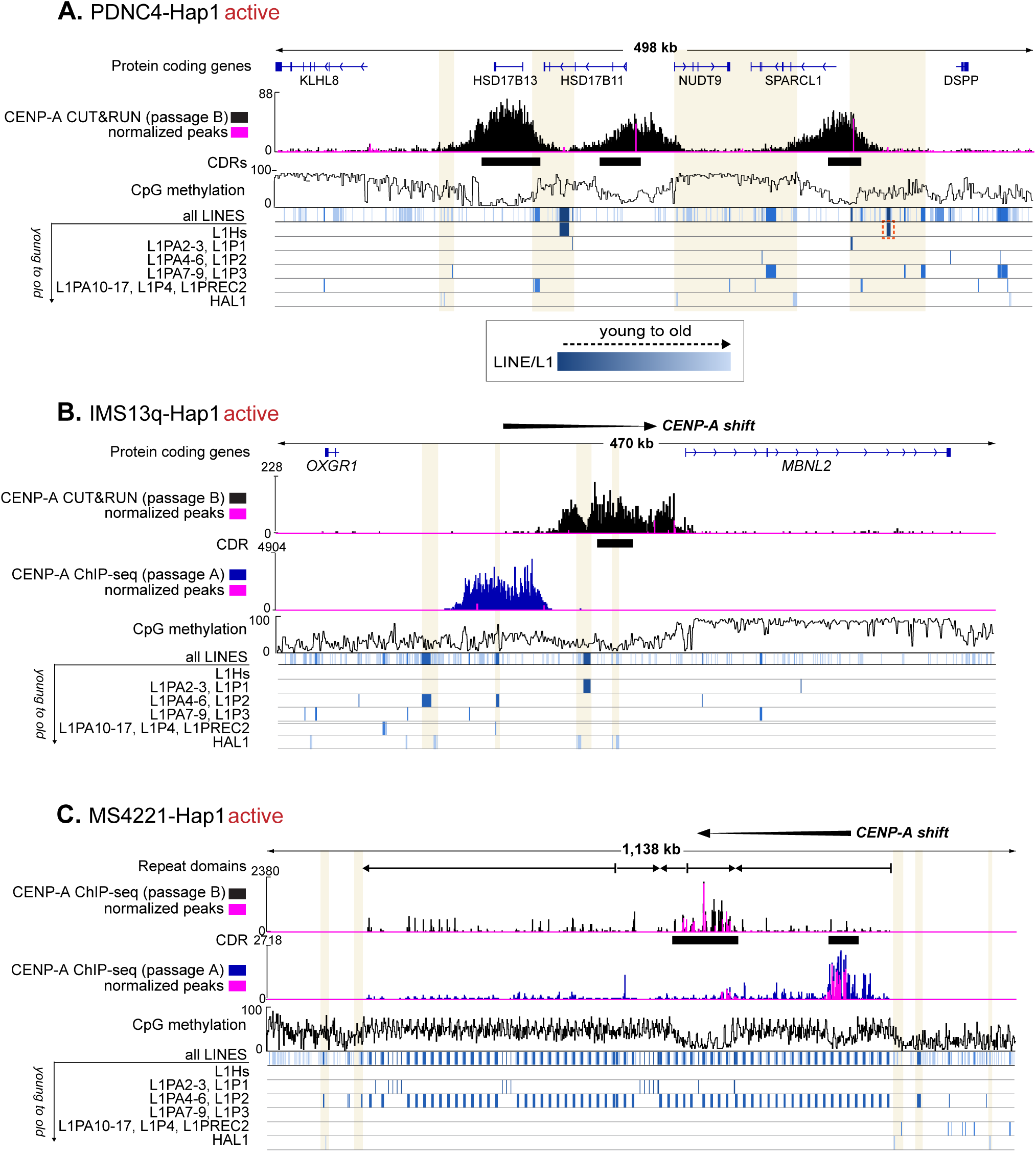
LINEs serve as boundaries for CENP-A spreading in NeoCens. Annotations for the active neocentromere in **(A)** PDNC4-Hap1, **(B)** IMS13q-Hap1, and **(C)** MS4221-Hap1. From top, tracks for each NeoCen are indicated on the left. All LINEs (L1s) are shown in one track denoted by phylogenetic age (colored based on blue gradient, left), followed by tracks for each of the individual LINE subclasses. A red box in **(A)** indicates an L1Hs that is specific to Hap1 and missing in Hap2. Additional tracks in **(B,C)** represent the Passage A CENP-A data with the shift in the CENP-A domain shown at the top.

However, we observed a pattern across all NeoCen loci wherein only young L1s (L1Hs, L1PA) and an older L1 subfamily, HAL1, reside at the boundaries of the CENP-A domains and/or the major CENP-A peaks in each active NeoCen-Hap1. This pattern is most striking in the PDNC4(chr4) NeoCen-Hap1 (Fig. 5A) where L1s flank each of the three CENP-A peaks and a single L1Hs insertion is specific to Hap1. In IMS13q(chr13) NeoCen-Hap1 (Fig. 5B) L1s form boundaries between the old (ChIP-seq, Passage A) and new (CUT&RUN, Passage B) CENP-A peaks. In MS4221(chr8) NeoCen-Hap1 (Fig. 5C), the composite units of the VNTR are L1PA-rich, yet we observe HAL1s at the CENP-A ChIP-seq domain boundaries. Together, these data suggest that young L1s and older HAL1s serve as boundaries that limit CENP-A spreading, even in the context of a NeoCen located on a tandem repeat array (MS4221).

### PDNC4 neocentromere induces transcriptional silencing by abolishing focal chromatin accessibility

Transcriptional activity across the NeoCen-Hap1/Hap2 loci was initially assessed using total RNA-seq and RNA polymerase occupancy measured by PRO-seq^10,24^ revealing that only two of the four genes located within the PDNC4 NeoCen are transcriptionally active (*HSD17B11, NUDT9*), and only one of the genes near the IMS13q NeoCen is active (*MBNL2*; Fig. 6A-B). Due to the low heterozygosity, we observed in both the PDNC4(chr4) and IMS13q(chr13) NeoCen loci, we were unable to phase short read PRO-seq data, preventing the determination of whether the transcriptional activity was equally present on each NeoCen haplotype. Evaluation of haplotype-phased ONT CpG methylation data demonstrated that the promoters of these three genes were hypo-CpG methylated on both haplotypes, with hypo-CpG methylation often associated with transcriptionally active promoters (Fig. 6A-B, SuppFig. 21, SuppTable 10). However, only one of these active promoters, *HSD17B11,* coincides with a CDR, making it impossible to discern transcriptional activity in Hap1 (active PDNC4 NeoCen) based on their methylation status.

**Figure 6.**
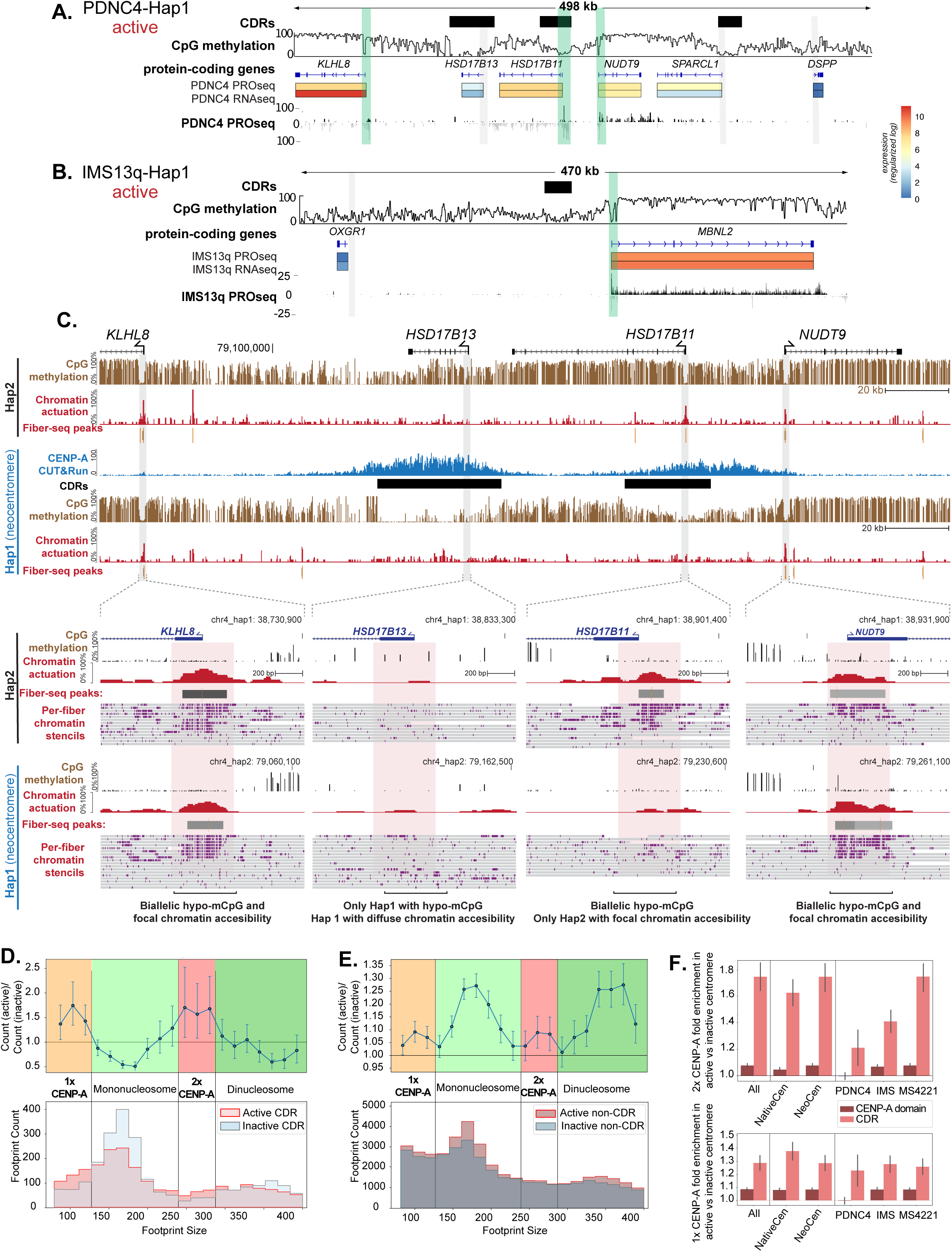
Neocentromeres remodel local chromatin and inhibit gene transcription. Gene expression based on PRO-seq and RNA-seq (log expression values are indicated as per key) for **(A)** PDNC4 and **(B)** IMS13q Hap1(active) NeoCens. Tracks are as indicated on the left, including stranded PRO-seq signal. Vertical green bars indicate expressed gene TSSs while grey indicate silenced TSSs. **(C)** (Top) PDNC4 NeoCen showing the Fiber-seq derived chromatin actuation and CpG methylation tracks from Hap1 and Hap2. CENP-A CUT&RUN is shown (blue) above Hap1 tracks, including the CDRs (black bars). (Bottom) Insets showing the Fiber-seq chromatin patterns at four TSSs within the above locus, including the single-molecule Fiber-seq chromatin patterns wherein individual fibers are marked by grey bars and single-molecule m6A events in purple ticks. Vertical pink bars indicate regions of monoallelic or biallelic hypo-methylation and focal chromatin accessibility. **(D)** (Top) Enrichment of binned footprint sizes in active NativeCen and NeoCen CDRs and their inactive counterparts (inactive CDR). Error bars indicate 99th percentile from 10,000x bootstrapped re-samplings of each population of footprints. (Bottom) Corresponding histogram of footprint sizes in active and inactive CDRs. **(E)** (Top) Enrichment of binned footprint sizes in active CENP-A domains (active CENP-A domain minus CDR(s)) vs their inactive counterpart (inactive CENP-A domain minus inactive CDR region). Error bars are as in **(D)**. (Bottom) Corresponding histogram of footprint sizes in active and inactive CENP-A domains. **(F)** Enrichment of 2X CENP-A footprints (Top) and 1X CENP-A footprints (Bottom) in centromeres and subpopulations of centromeres, split by CENP-A domain (minus CDR(s)) and CDR regions. Error bars are as in **(D)**.

Having demonstrated that we can accurately assemble and resolve the genomic and CpG methylation patterns of neocentromeres, we next sought to evaluate the chromatin patterns that are associated with neocentromeres and CDRs. Centromeres are known to be populated by a unique pattern of histone modifications (i.e., centro-chromatin) and CpG methylation^9^. Furthermore, we have recently demonstrated that chromatin within the CDR in active NativeCens adopts a dichotomous architecture characterized by both regions of chromatin accessibility as well as compacted dinucleosomes. Within the euchromatic genome, hypo-CpG methylation and chromatin accessibility are often a key feature of gene regulatory elements such as promoters. As the fibroblast PDNC4(chr4) active NeoCen-Hap1/Hap2 locus directly overlaps a gene promoter (*HSD17B11*) that typically demonstrates focal chromatin accessibility within fibroblasts, we sought to leverage our Fiber-seq data (SuppFig. 22, SuppTable 1) to directly assess the chromatin architecture along the neocentromere and at this gene, thereby determining if one or both haplotypes express *HSD17B11*. As Fiber-seq provides near-single-nucleotide resolution maps of chromatin architectures along individual ∼20kb long fibers, we were able to use this technology to accurately distinguish the chromatin architecture of the active NeoCen-Hap1 from the haplotype containing native euchromatin.

We found that focal chromatin accessibility at the *HSD17B11* promoter was completely lost in the active NeoCen-Hap1 (Fig. 6C). Rather, the region surrounding the *HSD17B11* promoter in NeoCen-Hap1 was marked by chromatin accessibility consistent with that of dichromatin. Furthermore, a focal site of chromatin accessibility within the gene body of *HSD17B11* also showed a loss of focal chromatin accessibility selectively along NeoCen-Hap1 (Fig. 6C). Importantly, this additional accessible element is also contained within the NeoCen-Hap1 CDR. In contrast, the neighboring *NUDT9* gene promoter, which is not within any CDR in NeoCen-Hap1, did not show a marked change in chromatin accessibility along NeoCen-Hap1 (Fig 6C). Overall, this demonstrates that although centromere chromatin is typified by hypo-CpG methylation and chromatin accessibility, this chromatin environment does not appear to permit the formation of focal chromatin accessibility in a form typically seen within the euchromatic genome. Moreover, these data resolved the NeoCen-Hap1 as the transcriptionally silenced allele for *HSD17B11* and implicates transcriptional silencing in neocentromere formation.

### Fiber-seq reveals haplotype-specific chromatin footprints and an enrichment of CENP-A dinucleosomes within active CDRs

Having demonstrated that haplotype-specific neocentromeres can disentangle the impact of CDR/CENP-A domains on focal chromatin accessibility, we next sought to explore whether haplotype-specific NeoCens and NativeCens can be leveraged to precisely measure the size of kinetochore-associated footprints. Unlike standard nucleosomes, CENP-A nucleosomes wrap 121 bp of DNA in humans^27,37^, with *in vivo* ChIP-MNase-seq studies showing that CENP-A protects 110-150 bp of αSat DNA^27,38^. Cryo-EM studies of the human constitutive centromere associated network (CCAN) have consequently only included a single CENP-A nucleosome, wherein the CCAN forms contacts with a CENP-A nucleosome and an adjacent linker DNA segment. However, evidence for such a structure *in vivo* has remained elusive. We hypothesized that directly comparing single-molecule footprints between the active and inactive haplotypes of the NativeCen and NeoCen loci would enable us to control for any sequence biases in footprint detection, thereby permitting the precise delineation of protein complex footprints formed within CDR/CENP-A domains and across the entire CENP-A region (minus the CDR) *in vivo*. Consistent with prior reports, we observed a clear enrichment of mononucleosome footprints within the CDR/CENP-A domains that were similar in size to CENP-A nucleosomes (*i.e.,* <130 bp in length [1x CENP-A]) (Fig. 6D). However, footprint sizes corresponding to a protein complex occupying a CENP-A mononucleosome and adjacent linker region (*i.e.,* ∼160-180 bp in length) were significantly depleted within CDR/CENP-A domains (Fig. 6D). In contrast, we observed a significant enrichment of footprint sizes corresponding to a protein complex occupying two CENP-A nucleosomes and the intervening linker region (*i.e.,* ∼240-280 bp in length [2x CENP-A]) (Fig. 6D). Notably, the enrichment of these 1x and 2x CENP-A footprints was similar along both the active NativeCen-Hap2 and active NeoCen-Hap1(Fig. 6D-F), consistent with these active NeoCens having similar function to that of active NativeCens. In contrast, regions adjacent to the CDRs, but within the overall CENP-A region, were significantly enriched for H3-containing mononucleosomes (*i.e.,* ∼150 bp in length) and dinucleosomes (*i.e.,* ∼350 bp in length), and a significant increase in >350 bp footprints, indicative of a more heterochromatin state (Fig. 6E, SuppFig. 23), indicating that the chromatin structural impacts of CDR/CENP-A domains extend to adjacent chromatin, even along neocentromeres.

## Conclusions

The completion of the first gapless human genome, T2T-CHM13^8^, provided a framework to understand transcription^10^, CpG methylation^9^ and CENP-A chromatin^10,11^ in the context of αSat monomers and HORs. However, αSats are absent in NeoCens, challenging our understanding of the driving genetic and/or epigenetic changes that accompany the silencing of an αSat-rich NativeCen and establishment and maintenance of CENP-A assembly at a new location lacking αSat sequences elsewhere on the same chromosome. Through our work herein, we identified the initial cause and epigenetic outcome of NativeCen inactivation and the epigenetic changes that define NeoCen establishment on one haplotype, distinguishing it from the non-NeoCen homologous region on the other haplotype. While IMS13q NativeCens are both active, in PDNC4 and MS4221 we find that a deletion within the αSat HOR region encompassing the CDR led to inactivation of one NativeCen and epigenetic rescue through a NeoCen on the same chromosome. Confirming earlier FISH studies^22,25^, the inactive NativeCen still recruits CENP-A nucleosomes in the αSat HORs near the deletion site but at levels likely too low to form a functional kinetochore, indicating there is a lower limit to the size of CENP-A domains and accompanying block(s) of hypomethylation that is required to recruit the 16-subunit constitutive centromere-associated network (CCAN) complex (reviewed in) and form a functional kinetochore in humans.

The appearance of a NeoCen on the chromosome arm to rescue the chromosome from loss in mitosis or meiosis indicates that CDR/CENP-A domains cannot re-form on another site within the same αSat HOR following a deletion of the entire CDR/CENP-A domain. Given the recent finding that younger αSats expand in copy number within NativeCen, termed layered expansions, it is possible that the CDR shifts with these expansions during the normal process of HOR evolution^11^, with the CpG hypermethylation boundary sliding along with it, but only within an array (Fig. 7A). If those regions of expansions and their accompanying CDR are deleted, as observed herein, then CENP-A assembly elsewhere in the αSat array may be blocked by the underlying CpG hypermethylation and chromatin compaction (Fig. 7A). Recent work has shown that CpG hypermethylation within NativeCens stops the spread of CENP-A nucleosomes, and the converse, hypomethylation within NativeCens allows spreading of CENP-A nucleosomes resulting in aneuploidy and cell lethality, indicating there is an as yet unknown upper limit to the size of the CDR/CENP-A domain that is compatible with life. Thus, the requisite for a small hypomethylated CDR within a much larger hypermethylated αSat array may indicate that the overall size of the CDR in humans is constrained by selection against instability and inviability. Within active NeoCens, we find CDR sizes like those observed in active NativeCens, indicating this size constraint is independent of underlying sequence.

**Figure 7.**
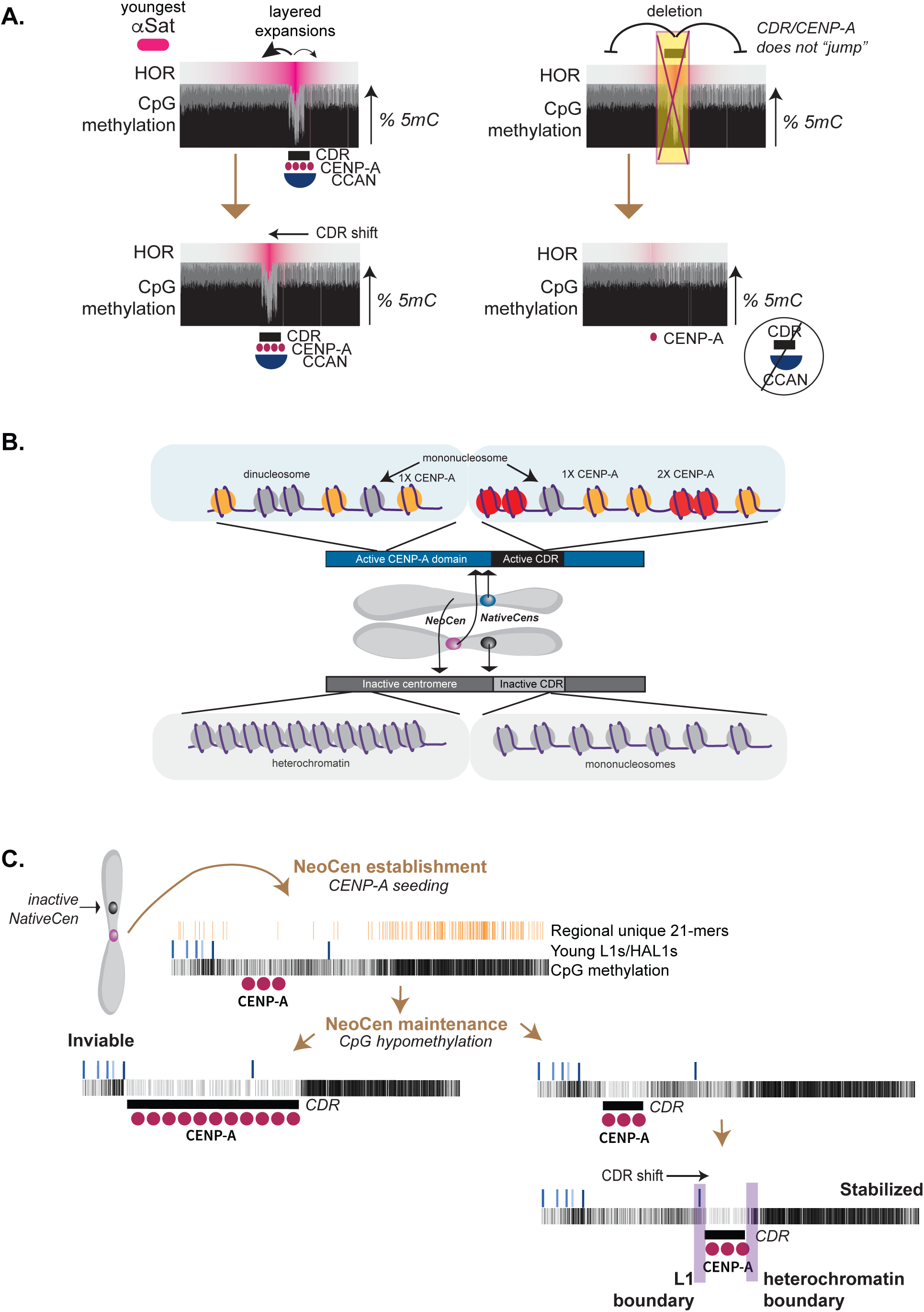
Deletion in αSAT within the CDR causes NeoCen activation and subsequent stabilization. **(A)** (Left) Within NativeCens, a shift in the CDR (bottom), and consequently CENP-A and the CCAN, accompanies layered expansions of the youngest αSats (pink). (Right) Following deletion of the CDR/CENP-A domain, hypermethylation blocks the assembly of CENP-A at a new site (top), leading to the loss of a CDR, with CENP-A levels too low to form a CDR and support CCAN assembly (bottom). **(B)** Active CDRs show patches of 1X and 2X CENP-A and mono- and di-nucleosome footprints. The alternate haplotype from the active CDRs show only patches of mono- and di-nucleosome footprints. Outside of the CDR within the CENP-A domain on active centromeres there is an enrichment of mono-and di-nucleosome footprints, with their inactive counterparts showing denser patterns consistent with heterochromatin. **(C)** Following inactivation of the NativeCen, a NeoCen is established by CENP-A seeding. NeoCen maintenance involves the establishment of a CDR, which if too large leads to instability/inviability. Once established, the CDR/CENP-A domain may shift to a more stable location with boundary elements that prevent spreading of CENP-A.

Using Fiber-seq data across haplotype-phased inactive and active NativeCens and NeoCens, we found enrichment of footprints corresponding to mono- (<130 bp) and di-nucleosomes (∼240–280 bp) of CENP-A, but depletion of intermediate-sized footprints of a single CENP-A and linker DNA (∼160–180 bp) in active NativeCen and NeoCen (Fig. 7B). This observation suggests that the structural constraints that define nucleosome patterning *in vivo* are not strictly defined by αSat monomer sizes and are independent of underlying sequence. Surprisingly, regions adjacent to the active CDRs were also enriched for H3-containing mono- and di-nucleosomes, indicating CENP-A impacts nearby chromatin structure of both NativeCens and NeoCens. In contrast, the Hap2 region homologous to the active CDR in the NeoCen-hap1 resembles that of the inactive NativeCen, with a more euchromatin-like state enriched for mononucleosomes that likely reflects the chromatin state prior to neocentromere formation surrounded by tightly packed chromatin (Fig. 7B). Finally, we find that NeoCen formation suppresses focal chromatin accessibility within the CDR implicating transcriptional silencing as result of neocentromere formation and indicating that NeoCens that form on genes or enhancers/promoters whose dosage is critical for viability would be incompatible with life. Given this finding, we predict that the transcripts for αSat that have been detected, albeit at low levels in normal cells^10,39^ are likely from αSats outside of the CDR/CENP-A domain; however this does not necessarily preclude these transcripts from playing a critical role in centromere identity and chromosome stability^40–42^.

Herein, we also observed CDR/CENP-A domain shifts within NeoCens. This CDR/CENP-A “sliding” at NeoCen occurs independent of the layered expansions of satellites observed in NativeCens, but may be part of the process of centromere stabilization after initial establishment. We observe that NeoCens that have shifted over time (MS4221 and IMS13q) are bounded by either LINEs or TSS of local genes whose gene body is hypermethylated. It is possible that NeoCens that support viability are located where LINEs, often bound by repressive marks such as KAP1/HP1, or regions of hypermethylation, can act to block the spread of CENP-A chromatin that would otherwise form a large domain that may be inherently unstable. Over time, the CDR/CENP-A domain may shift to a more stable location with strong boundaries.

A central question remains: *why do NeoCens form at these specific loci?* Comparative sequence analysis across the three cell lines revealed substantial differences in local genomic context at neocentromere loci, including the presence of tandem repeats (present in MS4221 but absent in PDNC4 and IMS13q), variation in overall repeat content (ranging from 30% to 75%), and the presence of protein-coding genes (present in PDNC4 and IMS13q, absent in MS4221). One potential commonality is the finding of low heterozygosity in non-repetitive NeoCen loci. While we cannot determine whether CENP-A assembly preferentially occurs at sites of low heterozygosity, or if low heterozygosity is the result of centromere activation on one allele, on an evolutionary timescale the latter could be a foundational step towards the fixation of Evolutionary New Centromeres (ENCs) by providing a substrate for the seeding of a NeoCen on the unaffected haplotype, necessitating the inactivation of the NativeCen on the same chromosome. Ultimately, rather than a common genetic feature defining NeoCen competency in the human genome, it is likely that observable active NeoCen are simply those that do not affect viability following the chromatin remodeling that accompanies CDR/CENP-A domain formation. One commonality among two of the cell lines is that the NeoCen forms in an area of lower overall methylation; however, the canonical CDR is only specific to the active NeoCen indicating that: 1) CENP-A mono- and di-nucleosome occupancy precedes CpG methylation in initial establishment of a new centromere (Fig. 7C), and that 2) once established, coincident CDRs and CENP-A domains are required for centromere maintenance and chromosome stability, as are boundaries that maintain CDRs within the size threshold needed to both recruit the CCAN and avoid aneuploidy (Fig. 7C).

A limitation of this study is that we do not know when the initial deletion at the NativeCen occurred (i.e. in early development or gametogenesis of one of the parents), nor the genetic/epigenomic landscape of either parent’s NativeCen on the affected chromosome. While this study is limited to only three NeoCen, future comparative studies of other NeoCen may further inform the minimum requirements for establishment and maintenance of active NeoCen. Through our work herein and more expansive studies, we may gain a better understanding of both clinical NeoCen cases and the initiation and fixation of ENCs that define species-specific karyotypes.

## Acknowledgements

The Center for Genome Innovation (in the Institute for Systems Genomics (ISG), University of Connecticut) and Bo Reese for sequencing support, Christine McCann for DNA extraction, Leighton Core for PRO-seq protocols, the ISG Computational Biology Core for high performance computing resources, the John and Donna Krenicki Professor in Genomics and Personalized Healthcare at UConn (RJO), and Ben Black for cell lines. Rajiv McCoy for assistance and funding to support low heterozygosity analyses. Research reported in this publication was supported, in part, by the NIH: R01GM123312-02 (RJO), R00GM147352 (GAL), R01HG010169 (EEE), F31HG012900 (DJT), R35GM133747 (RCM), U01HG013744 and 1DP5OD029630 (ABS). The content is solely the responsibility of the authors and does not necessarily represent the official views of the NIH. ABS holds a Career Award for Medical Scientists (Burroughs Wellcome Fund) and is a Pew Biomedical Scholar. EEE is an investigator of the Howard Hughes Medical Institute, RJO is an SAB for Colossal Biosciences, EEE is a scientific advisory board (SAB) member of Variant Bio, Inc.

## Author Contributions

**SJH:** data generation, assembly generation and validation, annotations analyses, figure generation and manuscript writing

**GAH:** CUT&RUN data generation, Fiber-seq data generation and pre-processing, figure generation

**MCA:** generation of αSat monomer annotations/tracks for all assemblies

**TWT:** Fiber-seq analyses and associated figures, manuscript editing

**DJT:** 1KGP data analyses and associated figures

**NMT:** k-mer statistics, transcription analyses and associated figures

**SN:** hosted UCSC browser tracks, generated hubs

**NP:** DNA extraction for ONT library preparation and sequencing

**KMM, KH**: generation of standard HiFi libraries and sequencing

**EEE:** funding for HiFi standard sequencing and UW personnel, manuscript editing

**GAL:** generation of assemblies, MS4221 ChIP-seq, manuscript editing

**ABS:** Fiber-seq analyses and interpretations, associated figures, manuscript editing

**RJO:** project conceptualization, oversight, funding, figure generation and manuscript writing

