## Supplemental Figures and Methods for "Haplotype-Resolved Genomics Reveals Conserved Chromatin Architecture and Epigenetic Constraints of Human Neocentromeres"

**SUPPLEMENTAL INFORMATION**

#### Table of Contents

*Supplemental Figures* ... 3

*Methods* ... 31

**Culturing of Human cell lines: PDNC4, MS4221, IMS13q** ... 31

**Oxford Nanopore Technologies (ONT) sequencing of PDNC4, MS4221, IMS13q** ... 31

**PDNC4** ... 31

MS4221 and IMS13q ... 31

ONT methods applied across all three lines ... 32

**PacBio HiFi sequencing of PDNC4, MS4221, IMS13q** ... 32

**Assembly, validation, and annotation of NativeCen and NeoCen regions in MS4221 (chr8), IMS13q (chr13), and PDNC4 (chr4)** ... 33

Assembly generation ... 33

Assembly validation and repeat annotation ... 34

Assembly and validation of chr4 NativeCen in MS4221 ... 35

HG002, T2T-CHM13, HG01114, HG00733 and HG02492 chr4 NativeCens ... 35

IMS13q breakpoint resolution ... 35

Derivation of ONT CpG DNA methylation ... 36

Identification of merged NeoCen haplotypes in IMS13q using individual methylated ONT reads ... 37

Assembly and regional based unique K-mer generation with Meryl ... 37

**Chromatin Immunoprecipitation sequencing (ChIP-seq) data processing – Passage A (IMS13q, MS4221 and PDNC4)** ... 38

Pre-processing and mapping ... 38

Post-processing ... 38

**Chromatin Immunoprecipitation sequencing (ChIP-seq) for Passage B MS4221** ... 38

ChIP-seq procedure ... 38

Illumina Library Preparation ... 39

Pre-processing, mapping, and post-processing ... 40

**CUT&RUN sequencing for Passage B PDNC4 and Passage B IMS13q** ... 40

CUT&RUN procedure ... 40

Illumina Library Preparation ... 40

Pre-processing, mapping, and post-processing ... 41

**Directed Methylation and Long-read sequencing (DiMeLo-seq)** ... 41

Experimentation, library prep, & sequencing ... 41

Data processing ... 42

**Chromatin accessibility assessment using Fiber-seq via Pacbio HiFi reads** ... 43

Fiber-seq experimentation ... 43

#### **Supplemental Figures**

#### Supplemental Figure 1

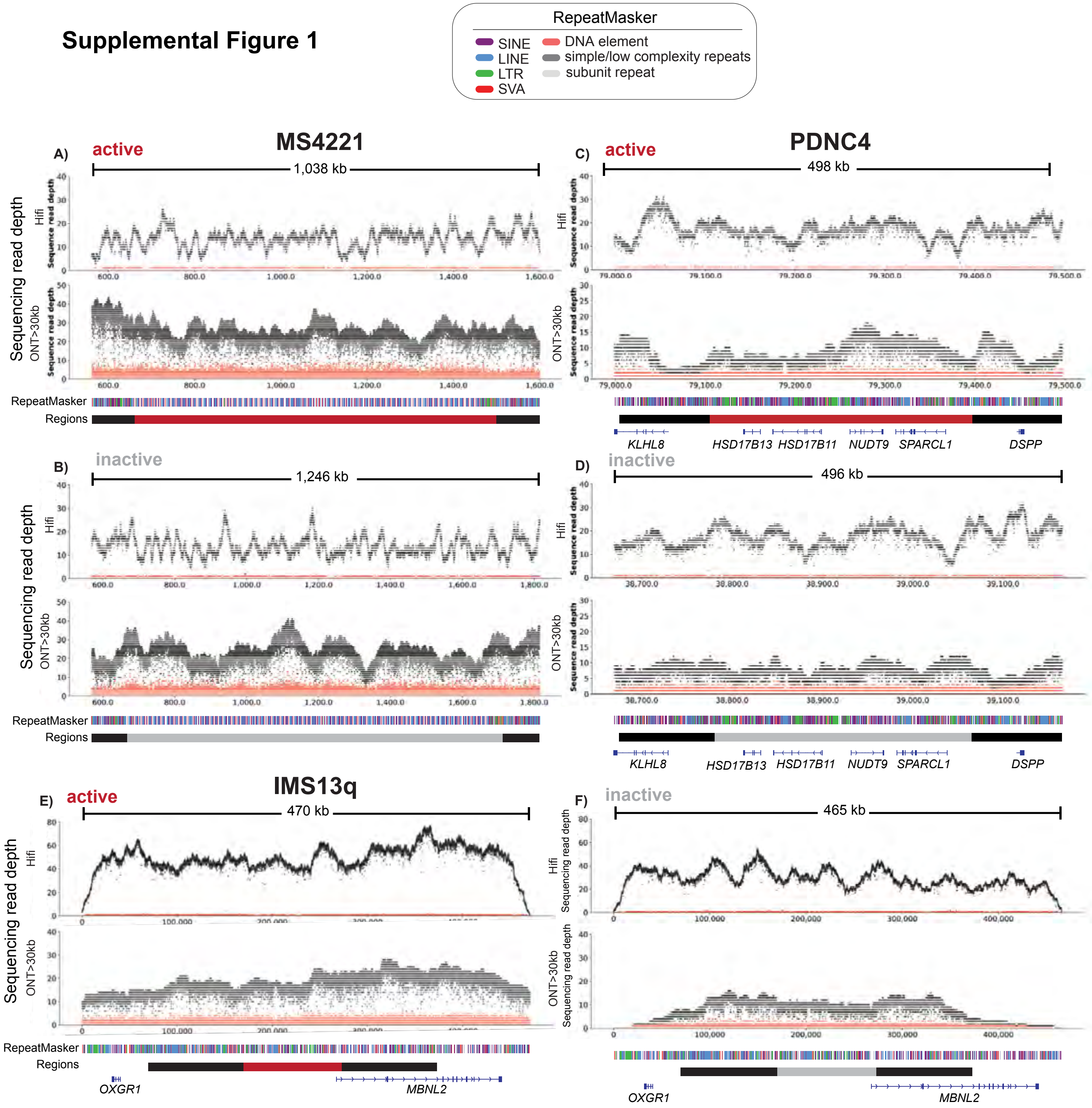

**Supplemental Figure 1. NeoCen assembly validation with NucFreq**. Long-read coverage plots via NucFreq across NeoCen haplotypes per assembly: MS4221 **(A, B)**, PDNC4 **(C, D)**, IMS13q **(E, F)**. PacBio HiFi reads (top) and ONT reads > 30kb (bottom) are shown per haplotype. Black dots represent the first most frequent base aligned, while red dots are the second. The "Regions" track denotes the CENP-A domain (red: active based on CDR presence, grey: inactive) and ~100kb flanking regions (black). RepeatMasker annotations (color coded per repeat class as per Key) and protein-coding genes (lifted NCBI RefSeq curated gene set) are shown in **(C-F)**.

**Supplemental Figures 2-4. NativeCen assembly validation with NucFreq and TandemQUAST.** Long-read coverage plots via NucFreq across NativeCen haplotypes per assembly: SuppFig. 2 IMS13q, SuppFig. 3 MS4221, and SuppFig. 4 PDNC4. HiFi reads (top) and ONT reads > 30kb (bottom) are shown per haplotype. Black dots represent the first most frequent base aligned, while red dots are the second. Tracks include (from top to bottom): RepeatMasker annotations (color coded per repeat class as per key in SuppFig. 2), the CENP-A domain (red: active based on CDR presence, grey: inactive), and the TandemQuast assessed region (black) corresponding to plots below. TandemQUAST plots are shown for Hifi reads (top) and ONT reads > 30kb (bottom) in sets of three, including: coverage, breakpoint ratio, and unique solid k-mer distribution. The latter two plots include light grey vertical bars highlighting regions of low coverage (<10 reads). A lack of red peaks (potential assembly errors) in the breakpoint ratio plot, and a lack of orange (multiple-clumps) and green (no-clumps) bars in the k-mer distribution plot indicate a high-quality assembly at the structural and sequence levels, respectively. See "Assembly validation and repeat annotation" in methods for further details on these plots.

### Supplemental Figure 2

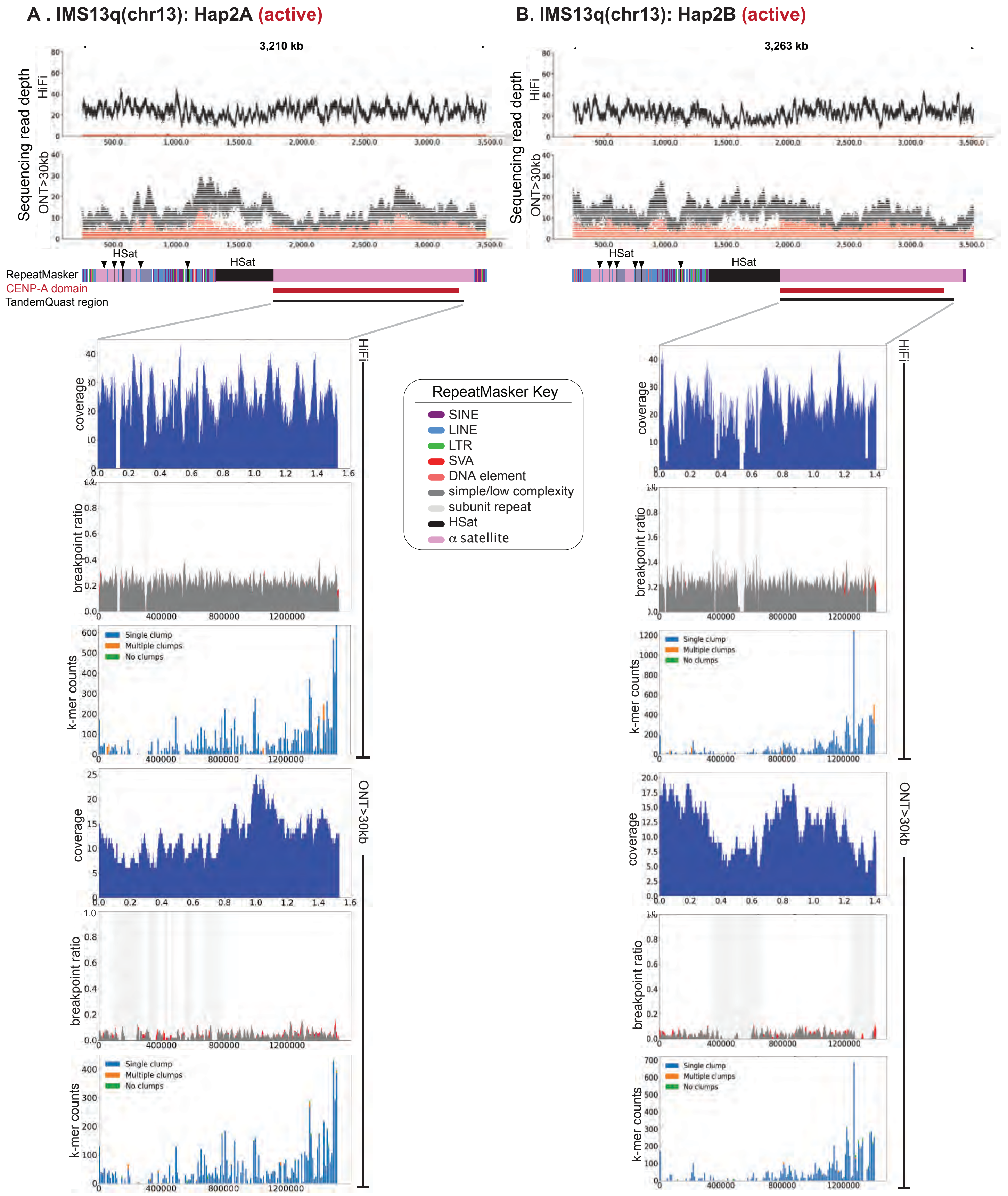

#### Supplemental Figure 3

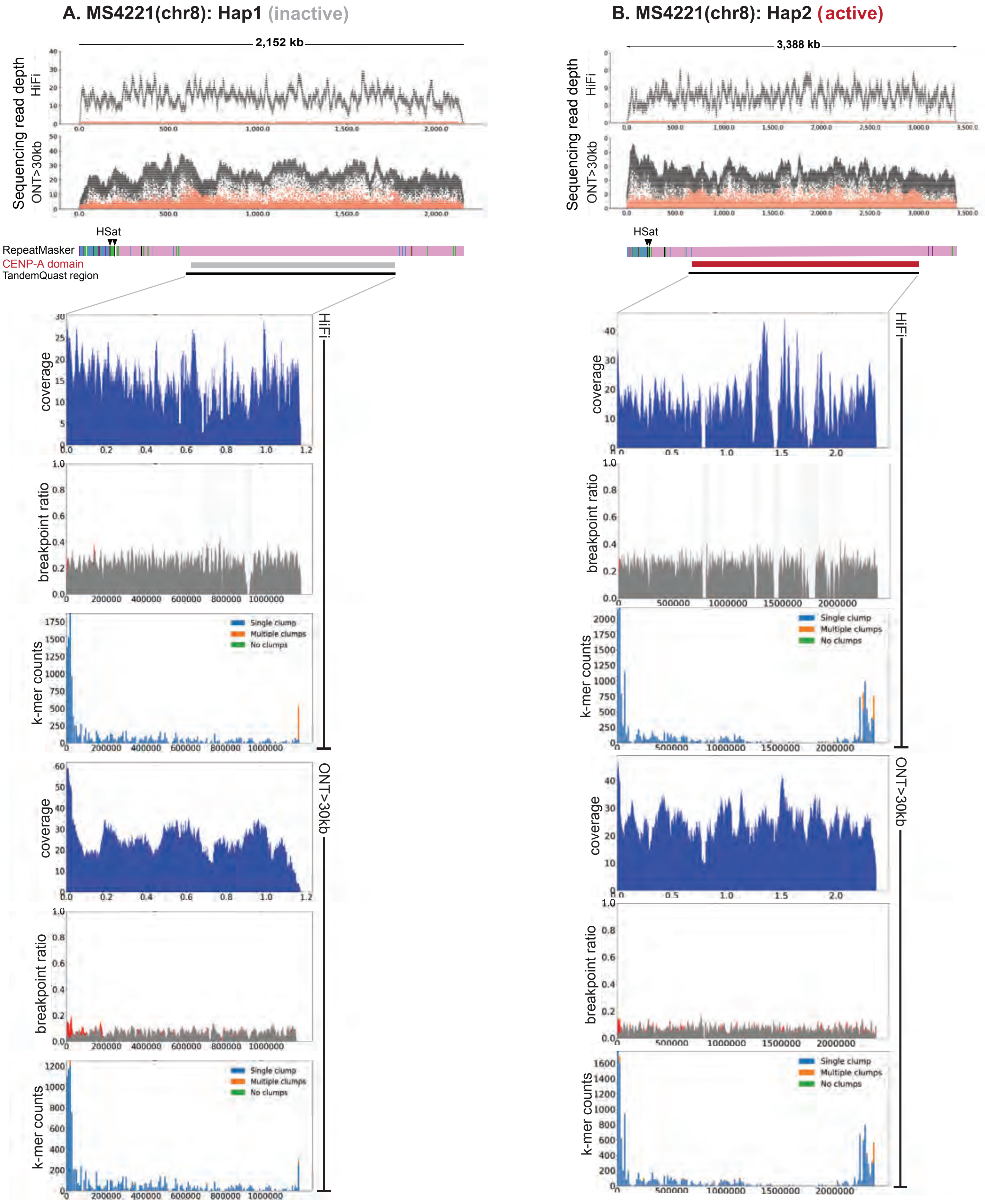

#### Supplemental Figure 4

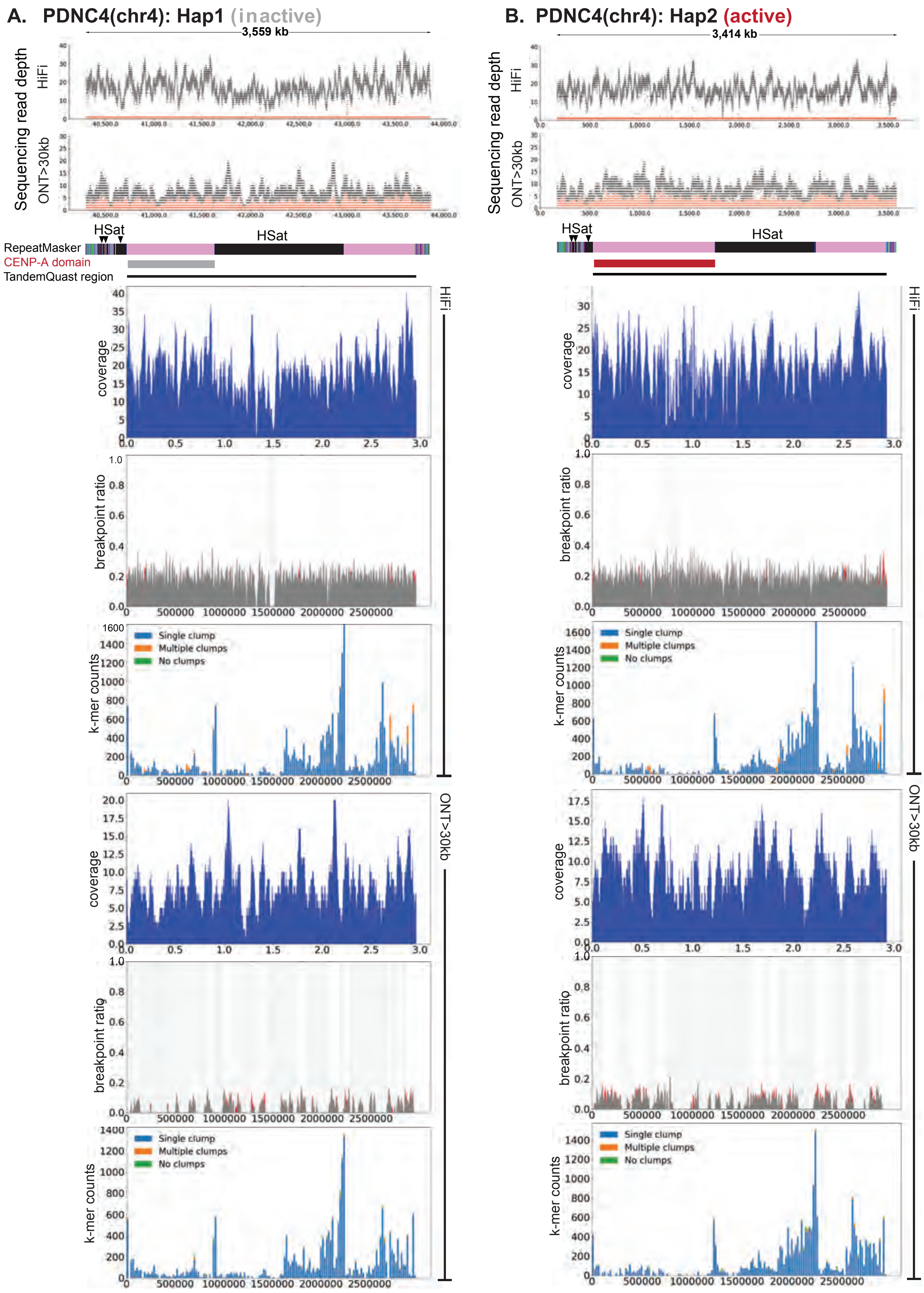

#### Supplemental Figure 5

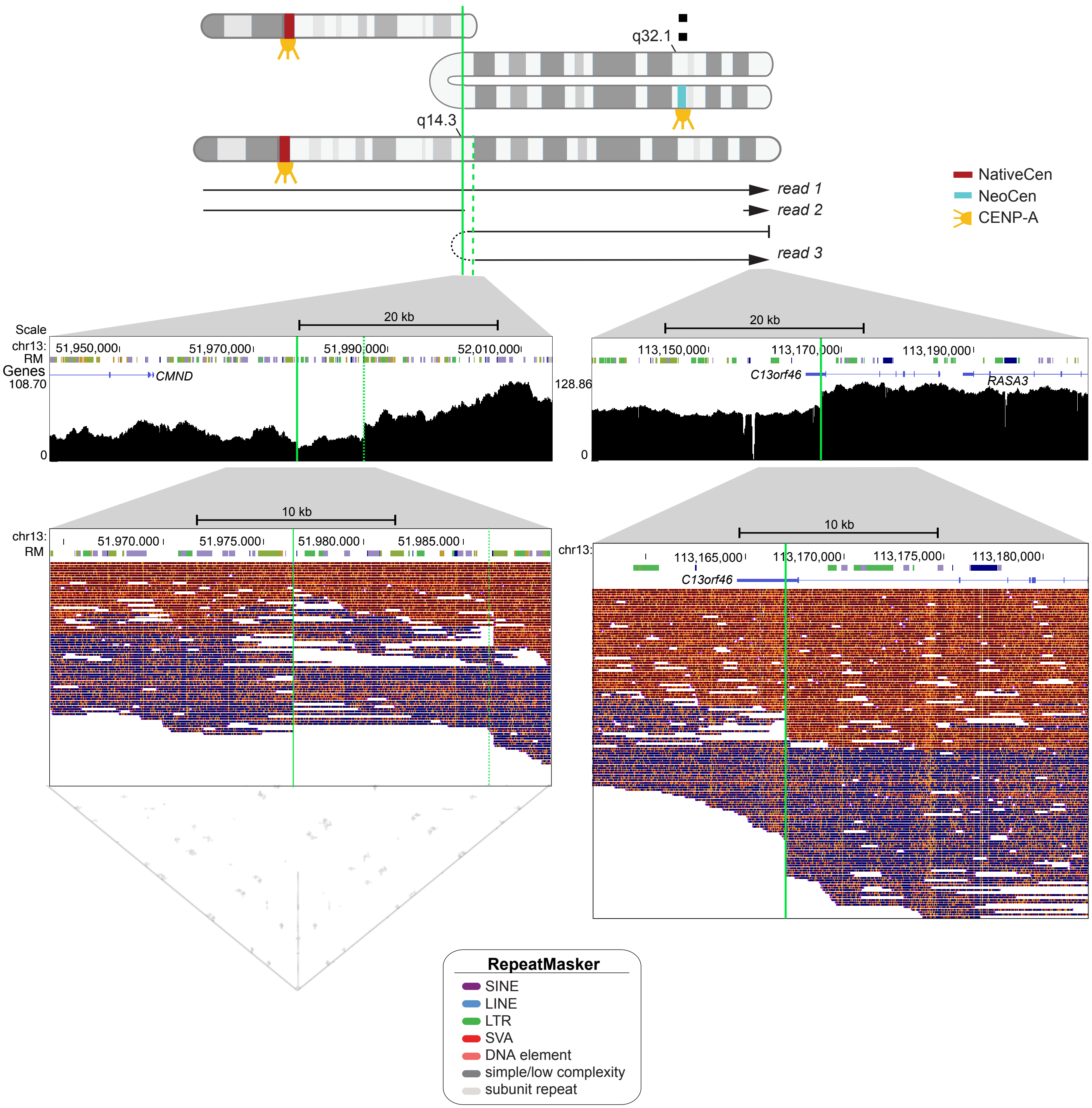

**Supplemental Figure 5. Resolution of the IMS13q breakpoints.** (Top) Ideogram as shown in Fig. 1 with homologous regions in alignment with one another. Three ONT read types are denoted as black arrows labeled as read 1-3. (Middle) T2T-CHM13 browser shots (coordinates on top) of >1kb ONT read alignments (Winnowmap) shown as density graphs (black) and zoomed into individually mapped reads (bottom)(red reads: mapped to reverse strand, blue reads: mapped to forward strand). dsDNA break sites denoted by solid green lines, and boundary of 10kb region involved in NAHR event denoted by dashed green line in the 13q34 breakpoint. (Bottom) Self-alignment of one 25kb ONT read spanning the corresponding region shown above indicating an inversion (perpendicular line). RepeatMasker (RM) annotations and protein-coding genes (CAT/Liftoff annotations V4) are shown.

### Supplemental Figure 6

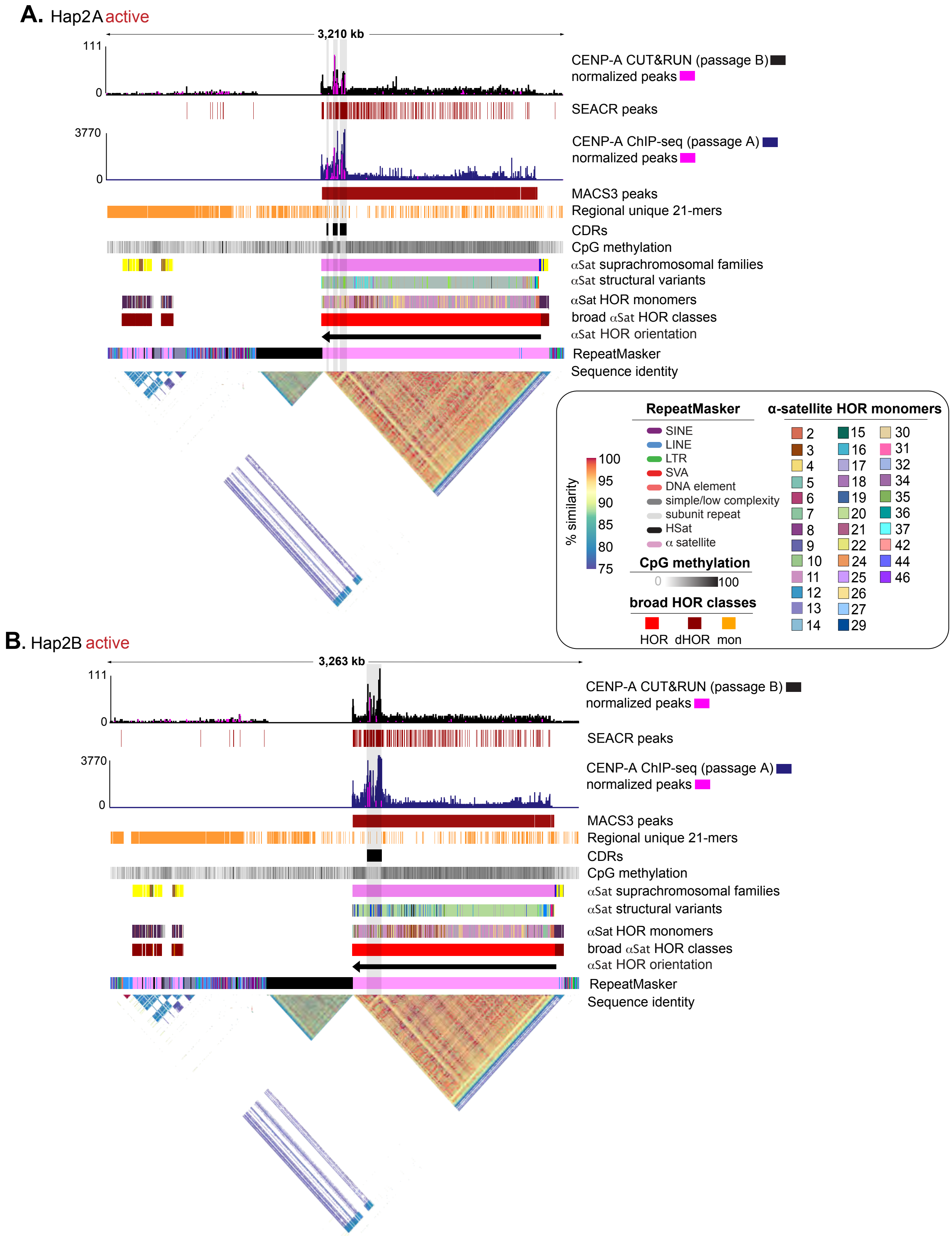

**Supplemental Figure 6. αSat variants, classes and families for the epigenetically homozygous NativeCen haplotypes of IMS13q (chr13).** A comparison of the chr13 NativeCen haplotypes, **(A)** Hap2A(active) and **(B)** Hap2B(active). Tracks are indicated on the right, color coded as per the Key. Tracks 8-12, αSat classifications shown from top to bottom as: suprachromosomal families (color key included in SuppTable 6), structural variants (SV), HOR monomers, broad HOR classes (color key included in SuppTable 6), and HOR orientation. (Bottom) StainedGlass pairwise sequence identity heatmap across the NativeCens, with a gradient denoting color scale for % identity (shared between haplotypes) and percent similarity of alignments.

#### Supplemental Figure 7

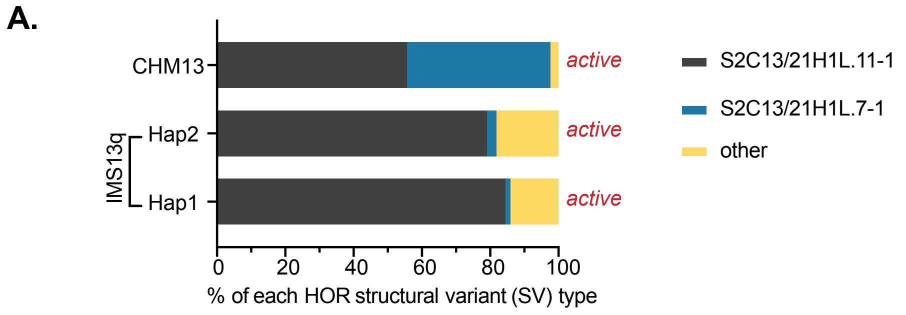

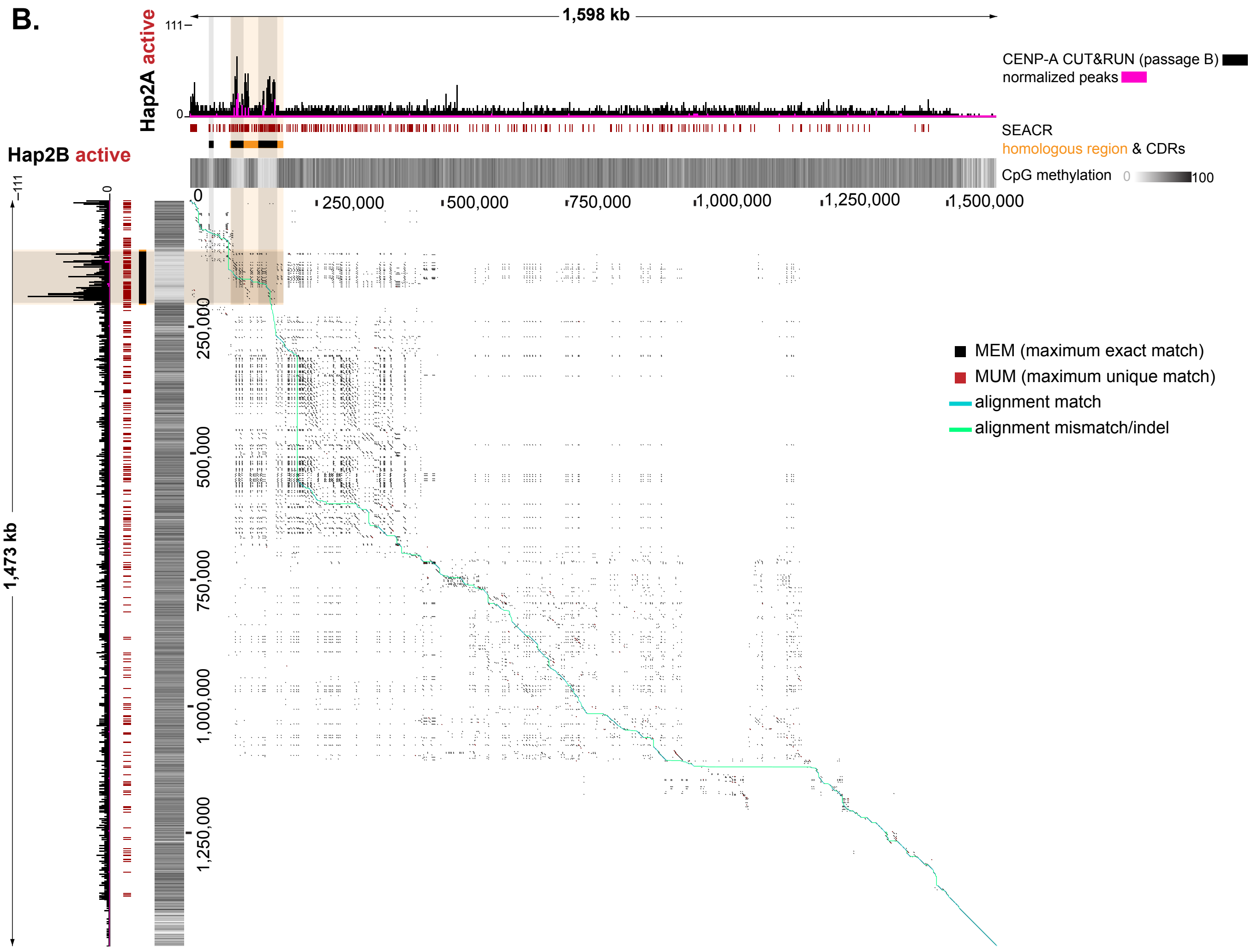

**Supplemental Figure 7. Comparative analysis of IMS13q (chr13) NativeCen HORs reveals high sequence and structural similarities.** **(A)** Composition of HOR structural variant (SV) types across the chr13 HOR haplotypes of IMS13q, and CHM13 for reference, based on base pair coverage. SV nomenclature is described extensively in (Altemose et al. 2022). **(B)** TandemAlignment pairwise dot plot between IMS13q HOR haplotypes (Hap2A: top, Hap2B: left) with coordinates shown along length of HOR and color key included (right). Tracks are denoted on the right. Shared region between haplotypes and CDR positions (orange and black, respectively) are highlighted across dot plot.

#### Supplemental Figure 8

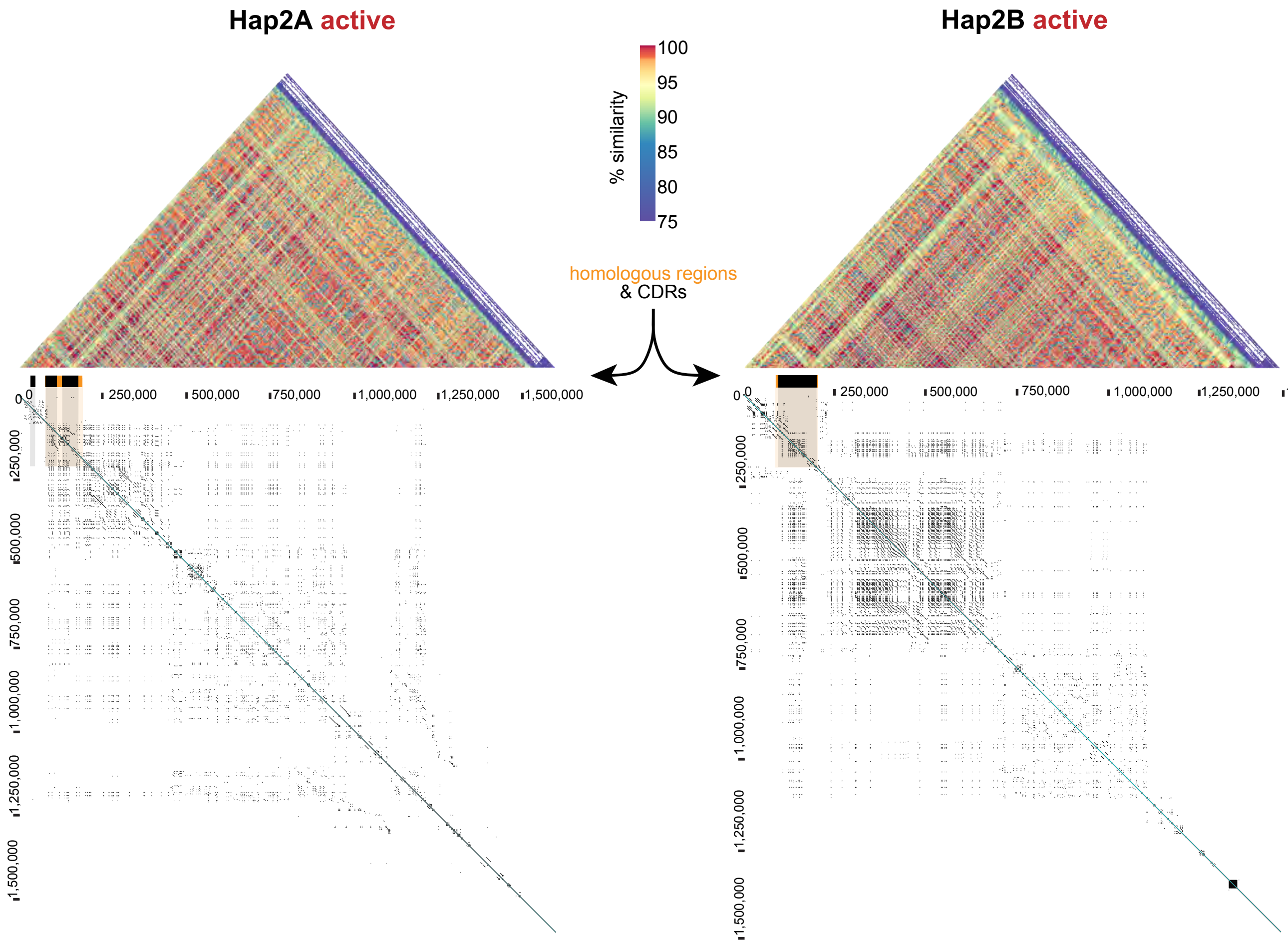

**Supplemental Figure 8. Individual StainedGlass and TandemAligner plots for IMS13q (chr13) NativeCen HORs reveals a high density of MEMs and recent αSat expansions.** StainedGlass pairwise sequence identity heatmaps (top) of each IMS13q HOR haplotype, with a gradient denoting color scale for % identity (shared between haplotypes). (middle) Homologous regions (orange) and CDRs (black) are highlighted across the self-alignment dot plot from TandemAligner (bottom) and correspond with a high density of MEMs.

### Supplemental Figure 9

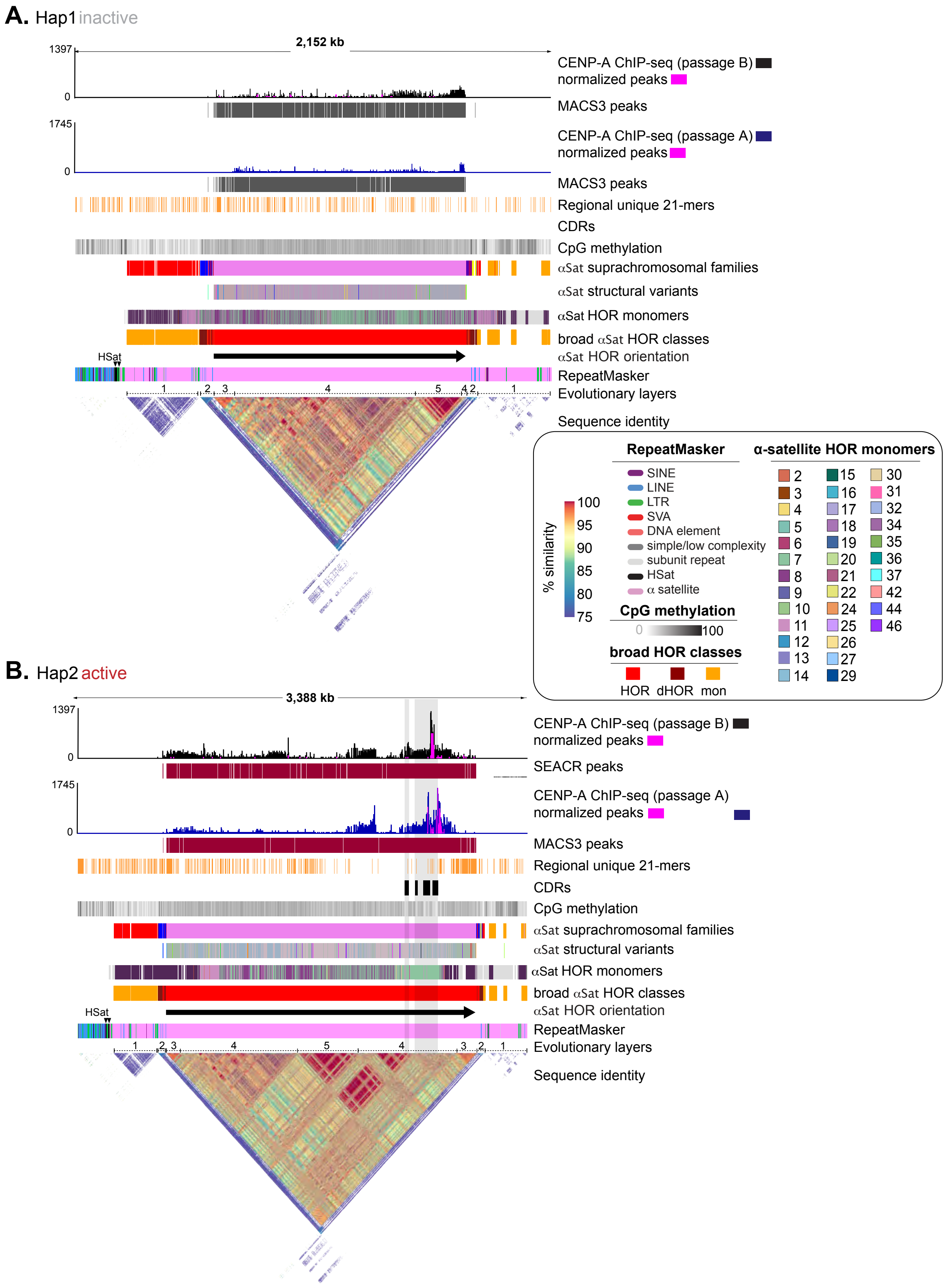

**Supplemental Figure 9. αSat variants, classes and families for the epigenetically heterozygous NativeCen haplotypes of MS4221 (chr8).** A comparison of the chr8 NativeCen haplotypes, **(A)** Hap1(inactive) and **(B)** Hap2(active), of MS4221. Tracks are indicated on the right, color coded as per the Key. Tracks 8-11, αSat classifications shown from most specific (suprachromosomal families; color keys included in Supp Table 6) to least specific (broad HOR; color coded as per Key). (Bottom) StainedGlass pairwise sequence identity heatmap across the NativeCens, with a histogram denoting the color scale (shared between haplotypes) and percent similarity of alignments.

### Supplemental Figure 10

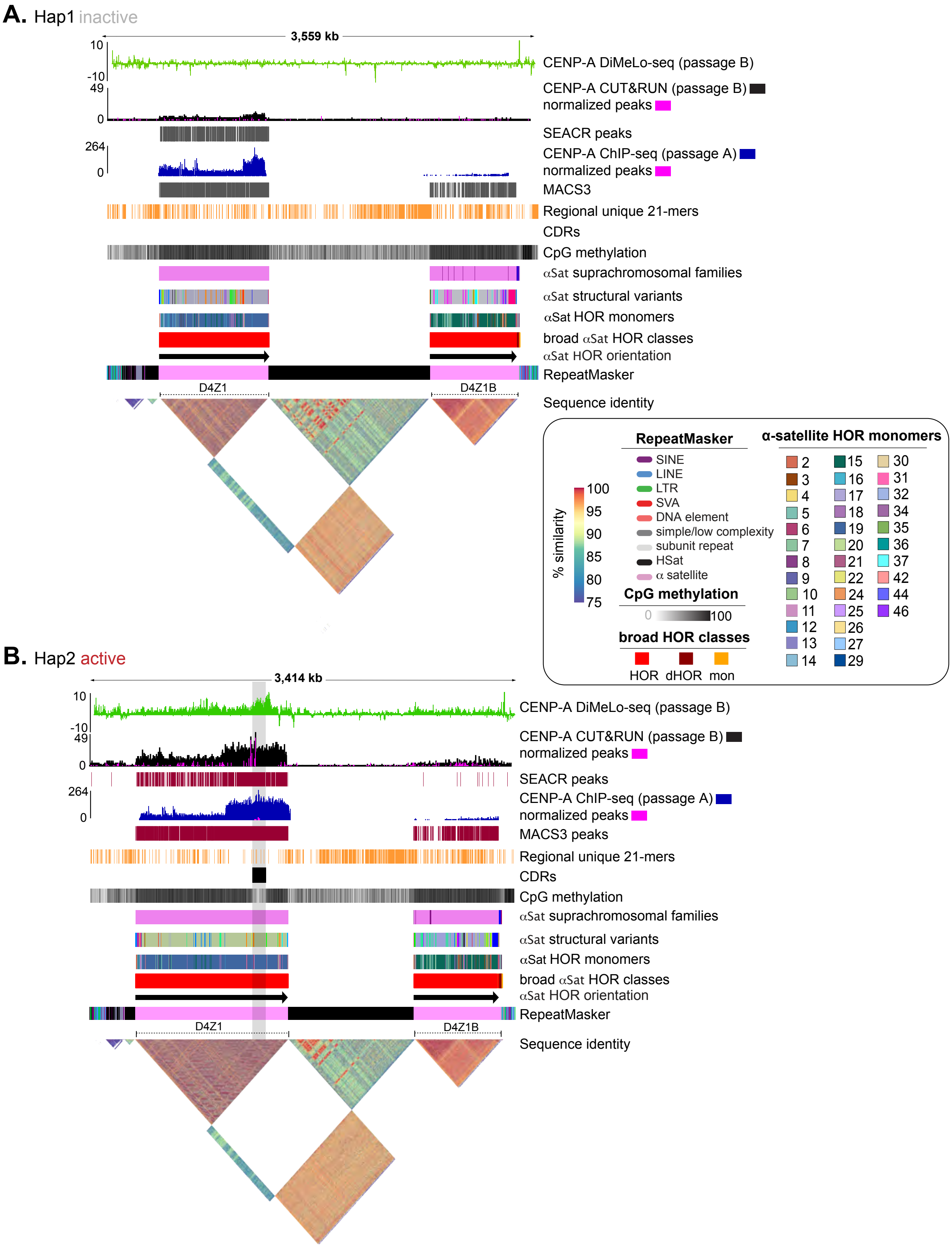

**Supplemental Figure 10. αSat variants, classes and families for the epigenetically heterozygous NativeCen haplotypes of PDNC4 (chr4).** A comparison of the chr4 NativeCen haplotypes, **(A)** Hap1(inactive) and **(B)** Hap2(active), of PDNC4. Tracks are indicated on the right, color coded as per the Key. Tracks 9-14, αSat classifications shown from most specific (suprachromosomal families; color keys included in Supp Table 6) to least specific (broad HOR; color coded as per Key). (Bottom) StainedGlass pairwise sequence identity heatmap across the NativeCens, with a gradient denoting color scale for % identity (shared between haplotypes) of alignments.

#### Supplemental Figure 11

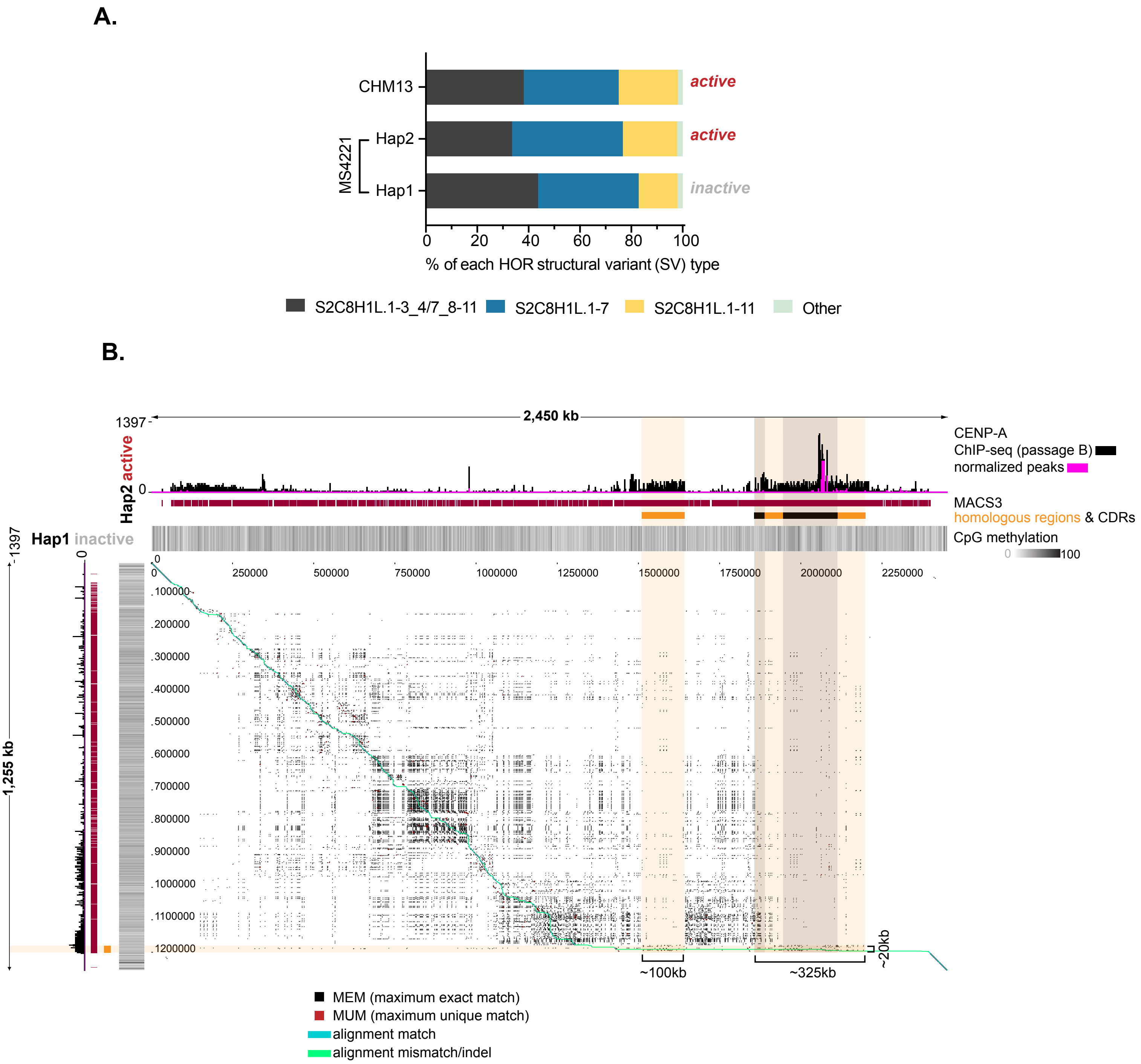

**Supplemental Figure 11. Comparative analysis of MS4221 (chr8) HORs reveals structural differences potentially linked to NativeCen instability. (A)** Composition of HOR SV types (as per SuppFig. 7) across the chr8 HOR haplotypes of MS4221, and CHM13 for reference, based on base pair coverage. **(B)** TandemAlignment pairwise dotplot between MS4221 HOR haplotypes (Hap2: top, Hap1: left) with coordinates shown along the length of the HOR and color key included (bottom). Tracks are denoted on the right. Homologous region between haplotypes and CDR positions (orange and black, respectively) are highlighted across the dot plot and fall in between the most recent αSat expansions (high density of MEMs). Brackets along the bottom indicate the size of the shared regions per haplotype.

#### Supplemental Figure 12

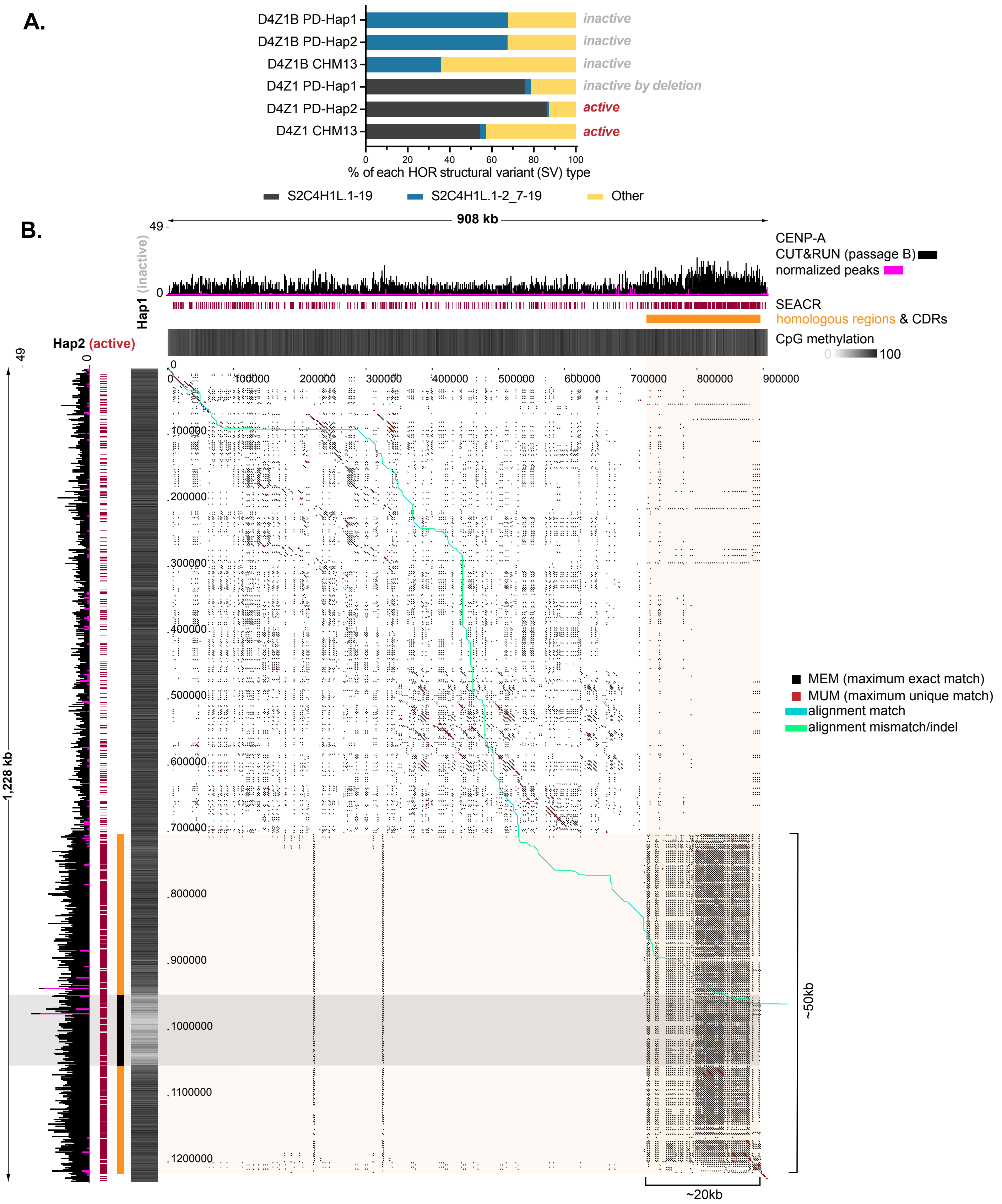

**Supplemental Figure 12. Sequence and structural variation of PDNC4 (chr4) HORs potentially linked to NativeCen instability. (A)** Composition of HOR structural variant (SV) types across the chr4 HOR haplotypes (D4Z1 and D4Z1B) of PDNC4, and CHM13, for reference, based on base pair coverage. **(B)** TandemAlignment pairwise dot plot between PDNC4 D4Z1 haplotypes (Hap1: top, Hap2: left) with coordinates shown along the length of D4Z1 and color key included (bottom). Tracks are indicated on the right. Homologous region and CDR positions (orange and black, respectively; both highlighted across dot plot) are shown. Brackets along the bottom indicate the size of the shared region per haplotype.

#### Supplemental Figure 13

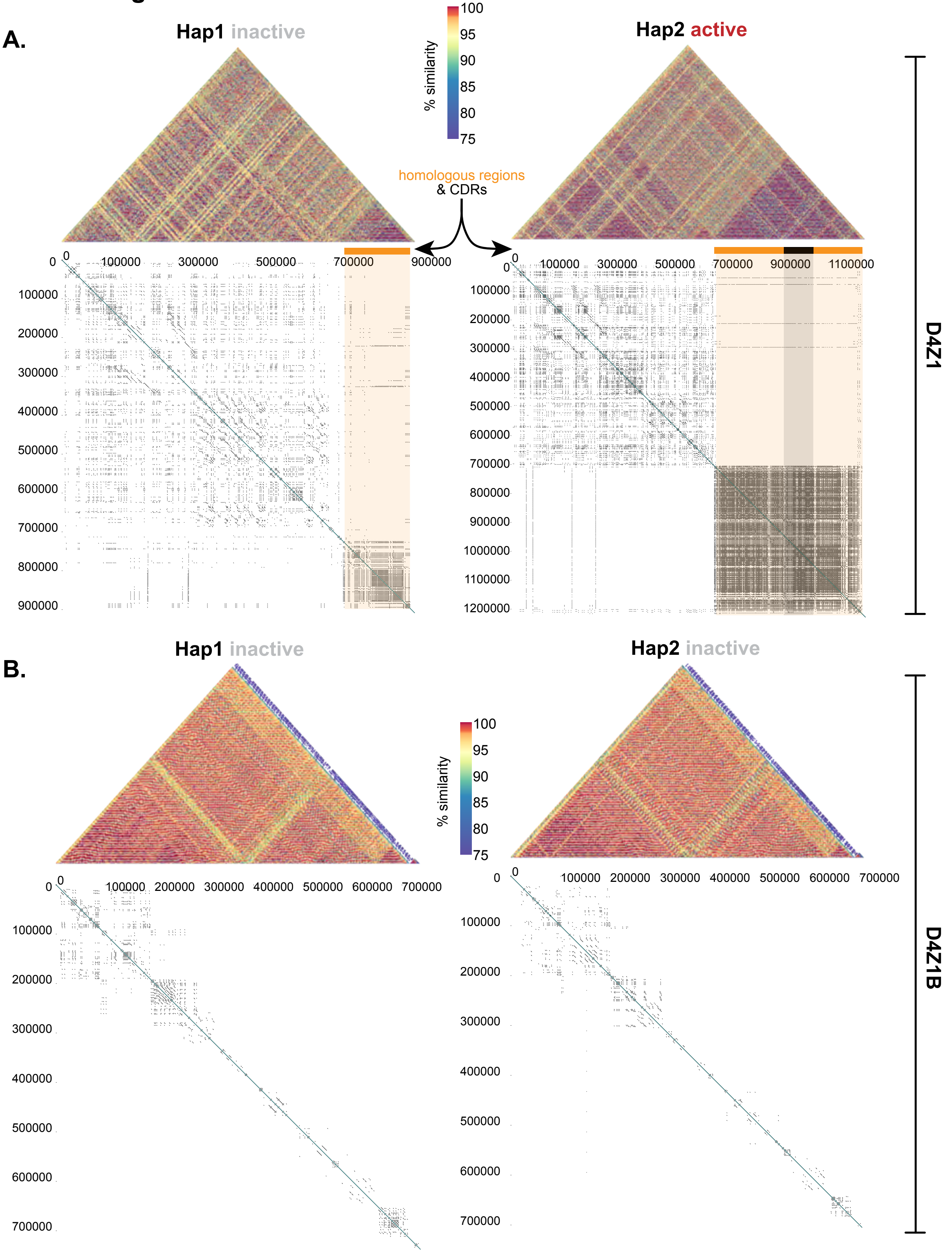

**Supplemental Figure 13. Individual StainedGlass TandemAligner plots for D4Z1 and D4Z1B of PDNC4 (chr4) NativeCen reveal a large region with a high density of MEMs and recent αSat expansions specific to D4Z1.** StainedGlass pairwise sequence identity heatmaps (top) of each PDNC4 **(A)** D4Z1 and **(B)** D4Z1B per haplotype, with a gradient denoting the color scale (shared between haplotypes). (middle) Homologous regions (orange) and CDRs (black) are highlighted across the self-alignment dot plot from TandemAligner (bottom) and correspond with a high density and large region of MEMs in D4Z1 of Hap2(active).

### Supplemental Figure 14

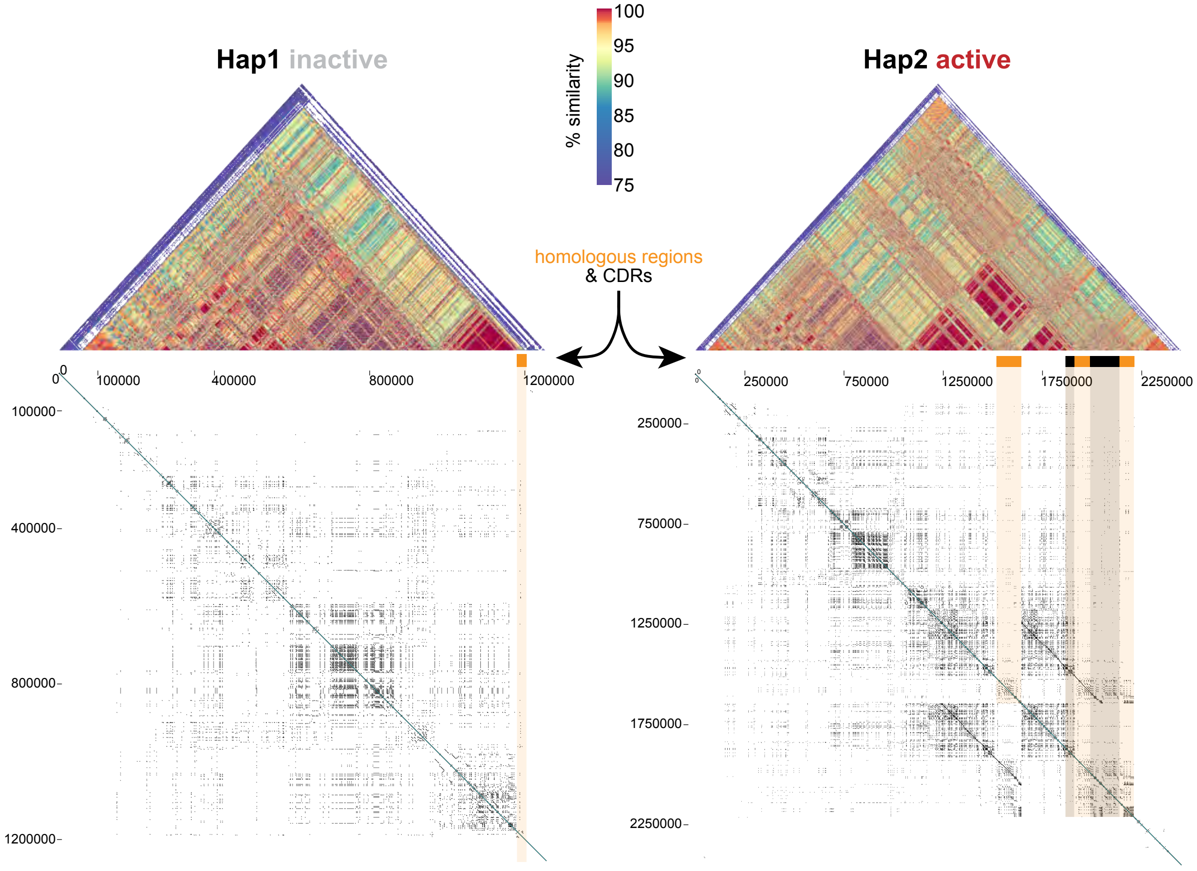

**Supplemental Figure 14. Individual StainedGlass TandemAligner plots for NativeCen HORs of MS4221 (chr8) NativeCens reveal a large deletion in the inactive haplotype.** StainedGlass pairwise sequence identity heatmaps (top) of each MS4221 haplotype, with a gradient denoting color scale for % identity (shared between haplotypes). (middle) Homologous regions (orange) and CDRs (black) are highlighted across the self-alignment dot plot from TandemAligner (bottom).

#### Supplemental Figure 15

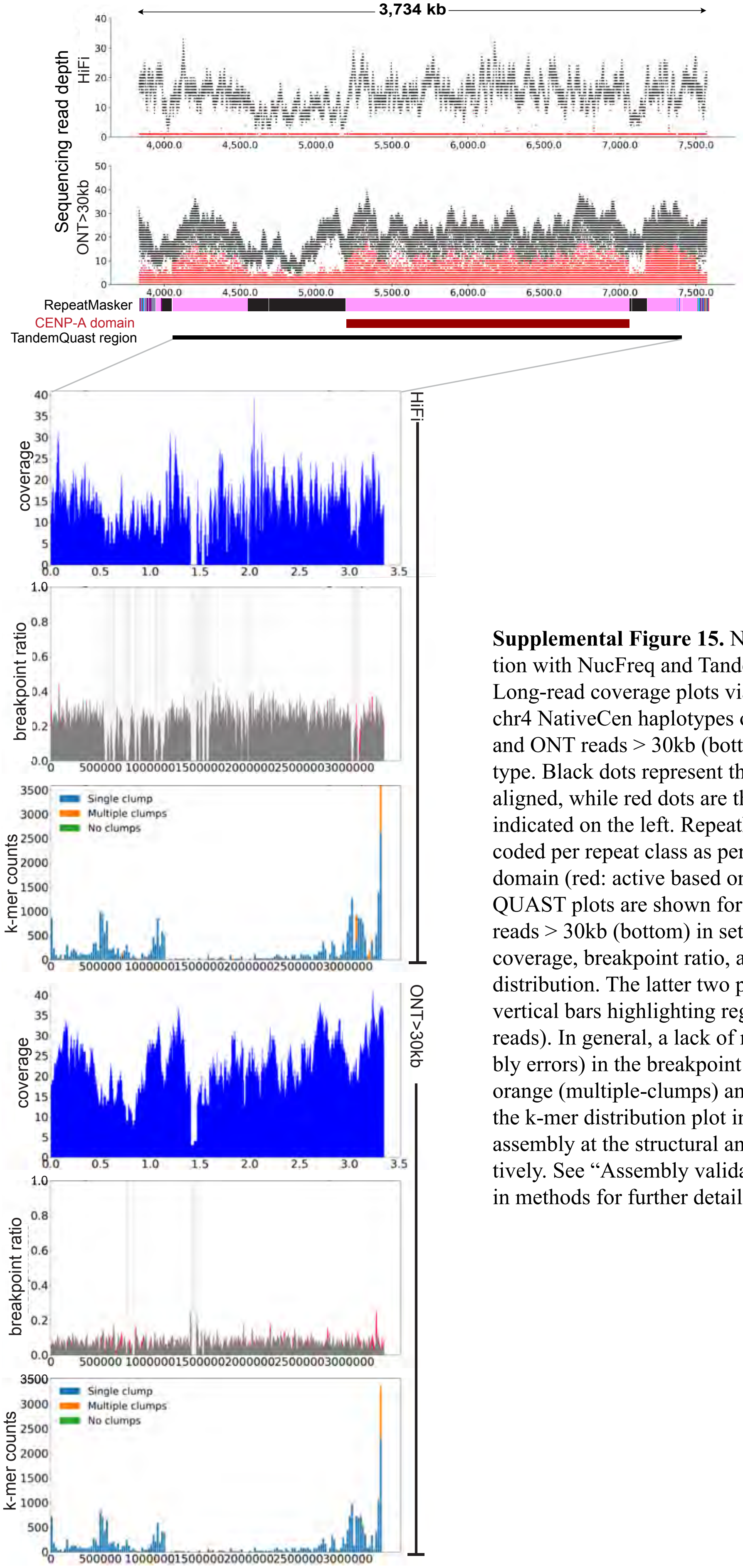

**Supplemental Figure 15.** NativeCen assembly validation with NucFreq and TandemQUAST of MS4221 chr4. Long-read coverage plots via NucFreq across one of the chr4 NativeCen haplotypes of MS4221. Hifi reads (top) and ONT reads > 30kb (bottom) are shown per haplotype. Black dots represent the first most frequent base aligned, while red dots are the second. Tracks are indicated on the left. RepeatMasker annotations are color coded per repeat class as per SuppFig. 2, the CENP-A domain (red: active based on CDR presence), TandemQUAST plots are shown for Hifi reads (top) and ONT reads > 30kb (bottom) in sets of three, including: coverage, breakpoint ratio, and unique solid k-mer distribution. The latter two plots include light grey vertical bars highlighting regions of low coverage (<10 reads). In general, a lack of red peaks (potential assembly errors) in the breakpoint ratio plot, and a lack of orange (multiple-clumps) and green (no-clumps) bars in the k-mer distribution plot indicate a high-quality assembly at the structural and sequence levels, respectively. See “Assembly validation and repeat annotation” in methods for further details.

### Supplemental Figure 16

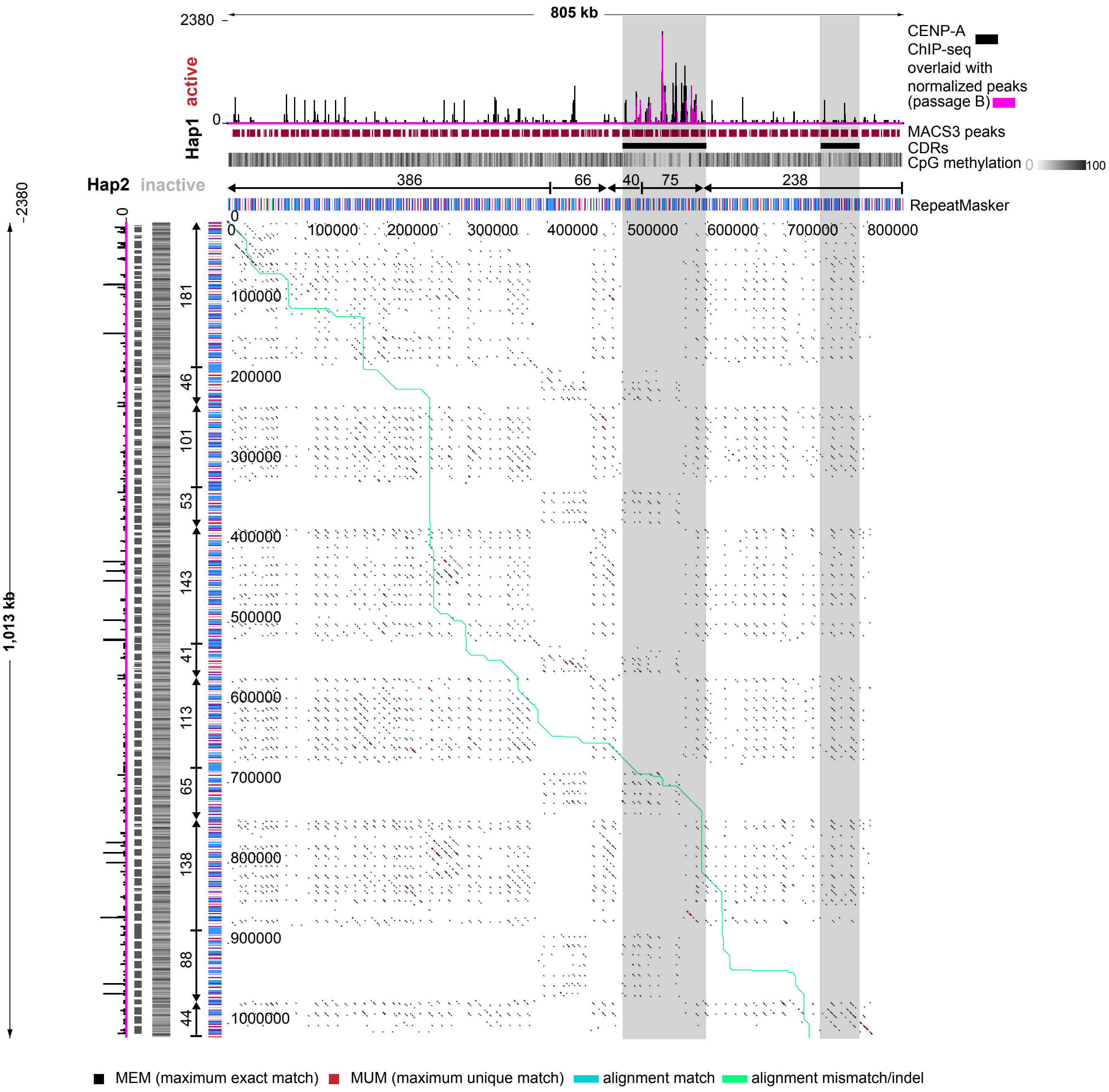

**Supplemental Figure 16. VNTR arrays of each haplotype of MS4221 differ size and number of composite units and domains.** TandemAlignment pairwise dot plot comparing the NeoCen active VNTR array on Hap1(active) and the inactive VNTR array on Hap2(inactive). Tracks are indicated on the right. CDR positions (black, highlighted in grey across dot plot) are shown. Above the RepeatMasker track (colored as per Key), arrows denote directionality of VNTR domains and numbers denote number of VNTR units per domain (included in SuppTable 9).

**Supplemental Figure 17. NeoCen genomic regions are distinct in PDNC4 and IMS13q.**
StainedGlass plots of active NeoCens on Hap1 of **(A)** PDNC4 and **(B)** IMS13q. **(C)** PDNC4 NeoCen haplotype comparison highlighting the ability to phase long-read sequencing data for CENP-A DiMeLo-seq compared to short-read CENP-A CUT&RUN (both from Passage B cells) to distinguish the active NeoCen versus inactive NeoCen, from both a zoomed-out view (top) and zoomed-in view (bottom). **(D)** In IMS13q, there are three homologous regions, one of which carries an active NeoCen.

#### Supplemental Figure 17

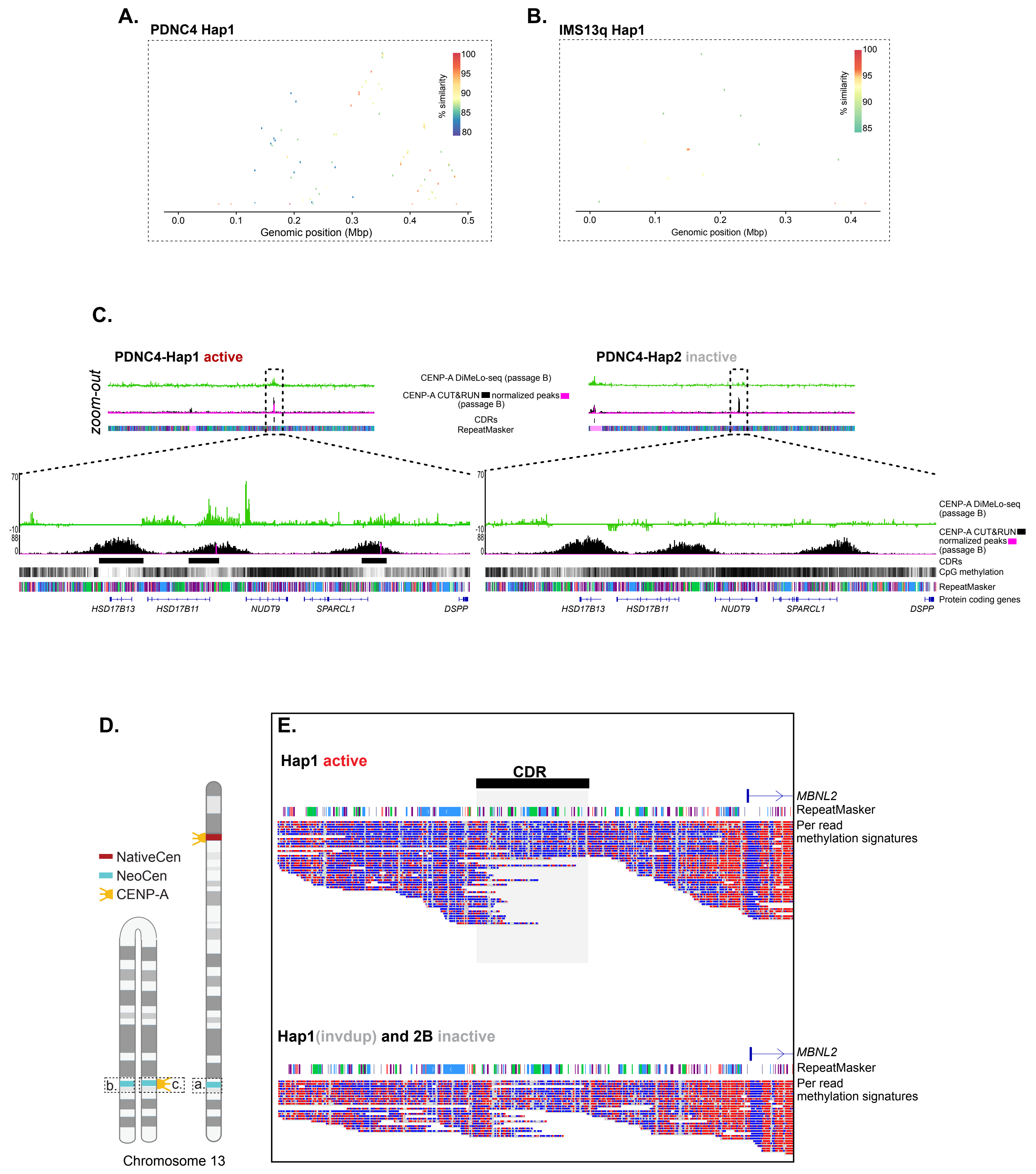

**Supplemental Figure 18. CpG methylation frequency comparisons across NeoCen CDRs, CENP-A domains, and flanking regions.** Methylation frequency was calculated as average methylation per Hap1(active) NeoCen CDR or corresponding CDR locus on the inactive Hap2 homologous region. **(A)** Histogram of average percent CpG methylation across all three lines **(B-C)** Boxplots are colored by activity (red: active, grey: inactive) for CDRs **(B)** and the region including ~100kb on each side (left flank, Lflank, and right flank, Rflank) and the CENP-A domain **(C)** for (from top) PDNC4, IMS13q, MS4221. Statistically significant differences were assessed with unpaired t-tests and marked with symbols between comparisons to denote the level of significance or non-significance (ns; p-value indicated in symbol Key).

#### Supplemental Figure 18

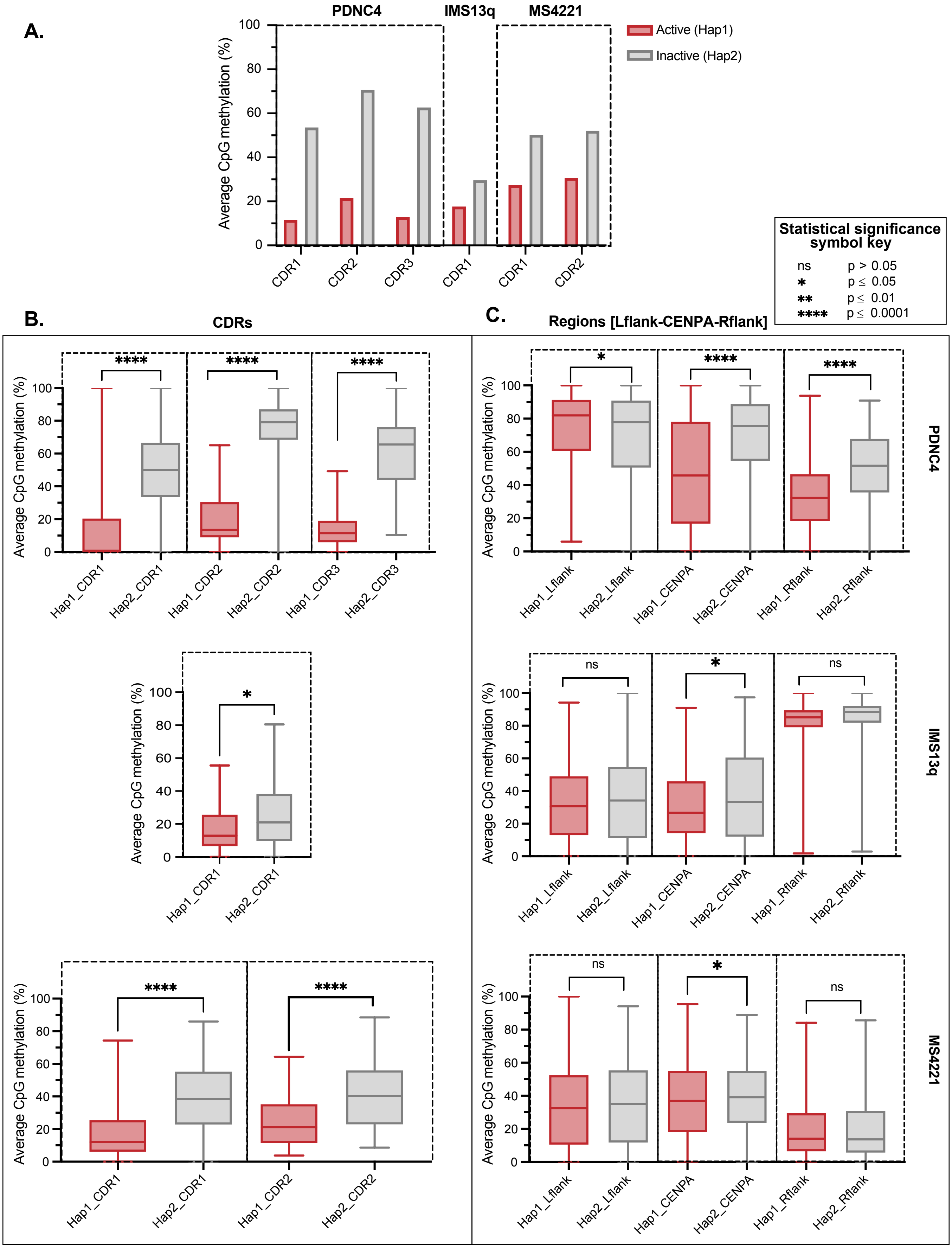

**Statistical significance symbol key**

| | |
|---|---|
| ns | $p > 0.05$ |
| * | $p \leq 0.05$ |
| ** | $p \leq 0.01$ |
| **** | $p \leq 0.0001$ |

### Supplemental Figure 19

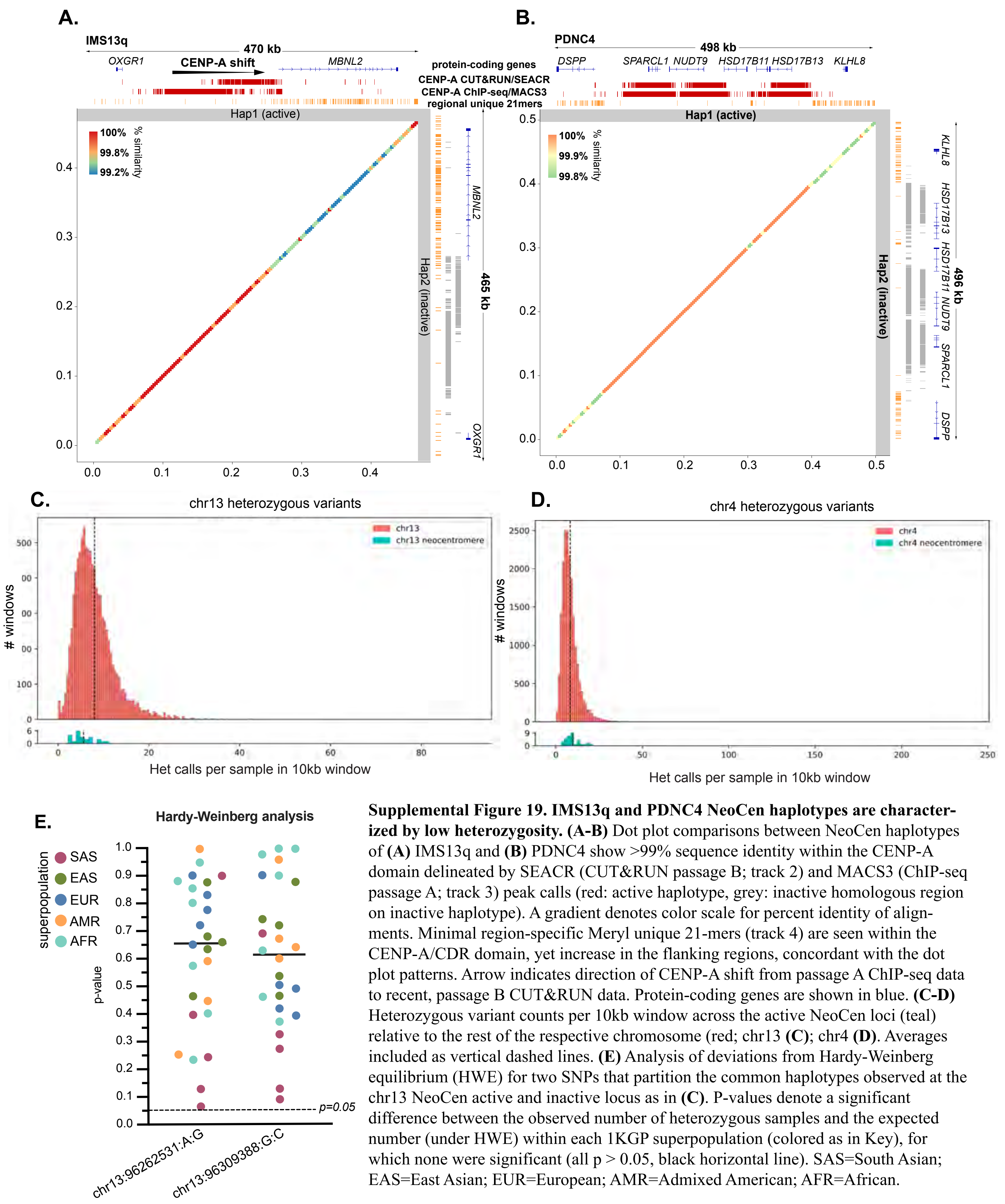

**Supplemental Figure 19. IMS13q and PDNC4 NeoCen haplotypes are characterized by low heterozygosity. (A-B)** Dot plot comparisons between NeoCen haplotypes of **(A)** IMS13q and **(B)** PDNC4 show >99% sequence identity within the CENP-A domain delineated by SEACR (CUT&RUN passage B; track 2) and MACS3 (ChIP-seq passage A; track 3) peak calls (red: active haplotype, grey: inactive homologous region on inactive haplotype). A gradient denotes color scale for percent identity of alignments. Minimal region-specific Meryl unique 21-mers (track 4) are seen within the CENP-A/CDR domain, yet increase in the flanking regions, concordant with the dot plot patterns. Arrow indicates direction of CENP-A shift from passage A ChIP-seq data to recent, passage B CUT&RUN data. Protein-coding genes are shown in blue. **(C-D)** Heterozygous variant counts per 10kb window across the active NeoCen loci (teal) relative to the rest of the respective chromosome (red; chr13 **(C)**; chr4 **(D)**. Averages included as vertical dashed lines. **(E)** Analysis of deviations from Hardy-Weinberg equilibrium (HWE) for two SNPs that partition the common haplotypes observed at the chr13 NeoCen active and inactive locus as in **(C)**. P-values denote a significant difference between the observed number of heterozygous samples and the expected number (under HWE) within each 1KGP superpopulation (colored as in Key), for which none were significant (all $p > 0.05$, black horizontal line). SAS=South Asian; EAS=East Asian; EUR=European; AMR=Admixed American; AFR=African.

**Supplemental Figure 20. Repeat composition varies between NativeCens and NeoCens, and amongst NeoCen loci. (All)** Only shown for active haplotypes (and, only Hap2B for IMS13q NativeCen) as other haplotype is near identical in the context of these analyses and therefore, redundant. **(A-C)** Dashed boxes highlight NeoCens when compared to NativeCens. **(A)** Repeat content (RepeatMasker) as fraction of total base pairs per CENP-A domain. Repetitive bases broken down into interspersed repeats (TEs and CHM13-derived composite subunits; teal), tandem repeats (satellites, simple/low complexity repeats, and RNAs; coral), or neither (non-repeats; grey). **(B)** CENP-B motif density was calculated as number of CENP-B motifs (based on FIMO assessment) normalized to total CENP-A domain length. The n values show the number of CENP-B motifs within each CENP-A domain. **(C)** AT content (based on QUAST) as fraction of total base pairs per CENP-A domain shown on limited y-axis (50-65% range) for enhanced resolution. **(D)** as in **(A)**, but NeoCens only and encompassing the ~100kb flanking regions (Lflank and Rflank). On the far right, the IMS13q ChIP-seq CENP-A domain for passage A is included for comparison. **(E)** Repeat class composition (RepeatMasker; color coded as per Key) as fraction of total base pairs per NeoCen region defined as [Lflank-CENPA domain-Rflank]. **(F)** Repeat types as in **(E)** but broken down into the three regional components per NeoCen, shown as a line graph to highlight the shifts between regions. **(G)** LINE subfamilies grouped by relative age and shown as fraction of total LINE base pairs per CENP-A domain (left, including IMS13q ChIP-seq passage A and CUT&RUN passage B) and NeoCen region (right) defined as [Lflank-CENPA domain-Rflank].

### Supplemental Figure 20

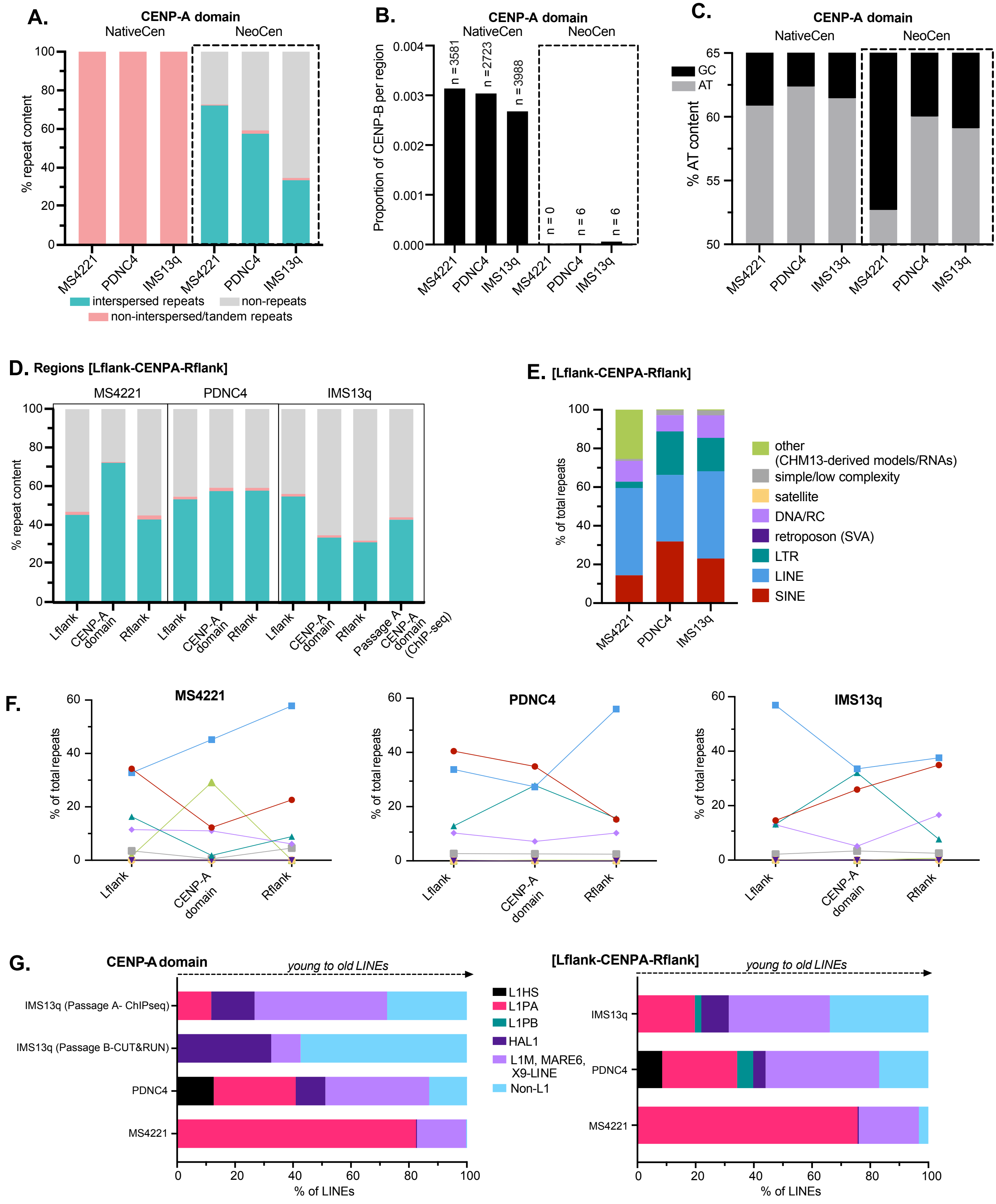

### Supplemental Figure 21

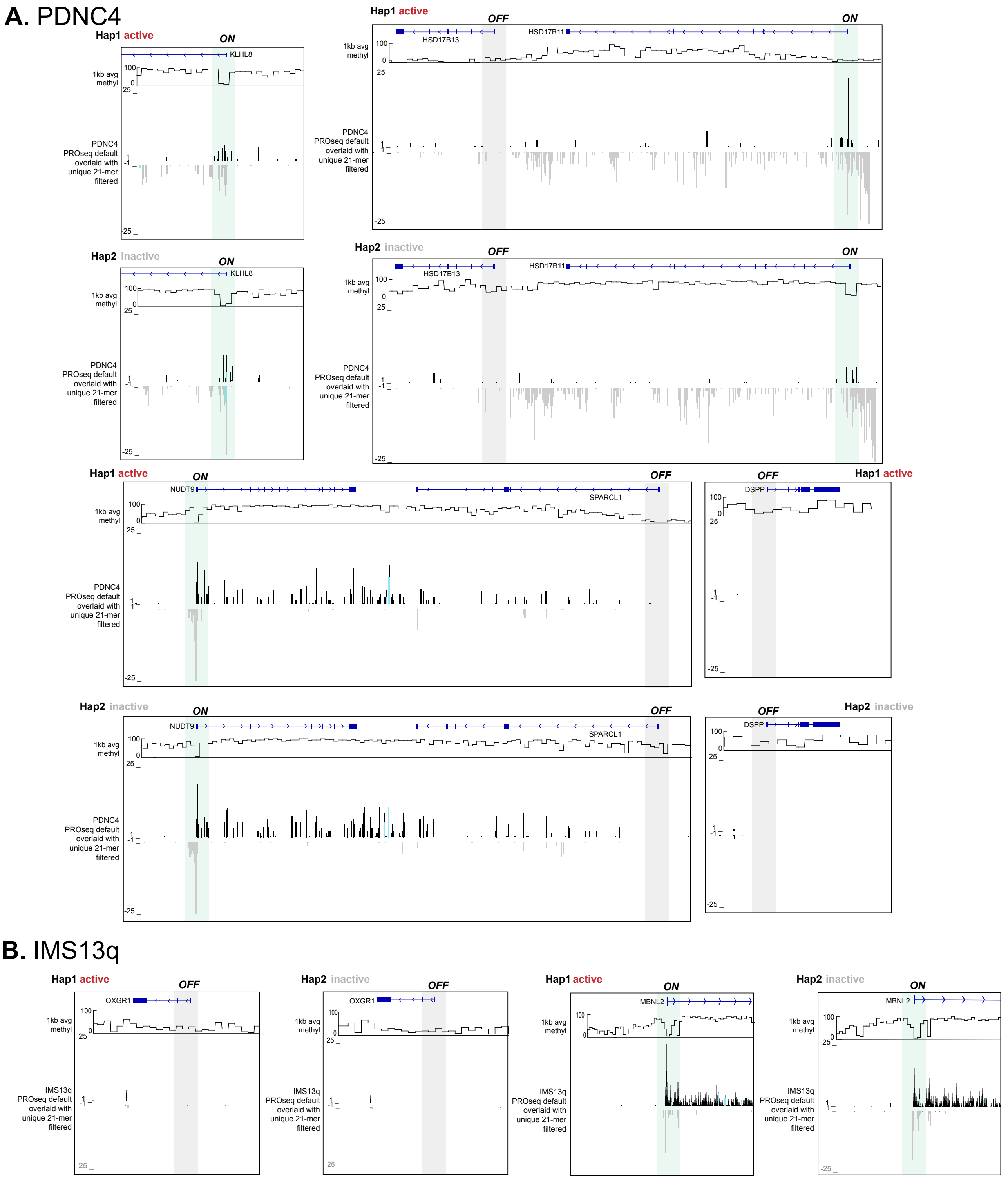

**Supplemental Figure 21. Protein-coding gene expression at the NeoCens of PDNC4 and IMS13q.** Zoomed-in IGV browser shots of each protein-coding gene promoters in **(A)** PDNC4 and **(B)** IMS13q NeoCens. Each browser shot includes protein-coding gene annotations, CpG methylation smoothed into 1kb windows, and stranded PRO-seq profiles, from their respective cell line, mapped with Bowtie2 overlaid with assembly-based Meryl unique 21-mer filtered coverage (bright blue). Hap1(active) and Hap2(inactive) are stacked for comparison between haplotypes. Gene promoters are highlighted as either “ON” (green) or “OFF” (grey) based on the association between PRO-seq signal and CpG methylation.

### Supplemental Figure 22

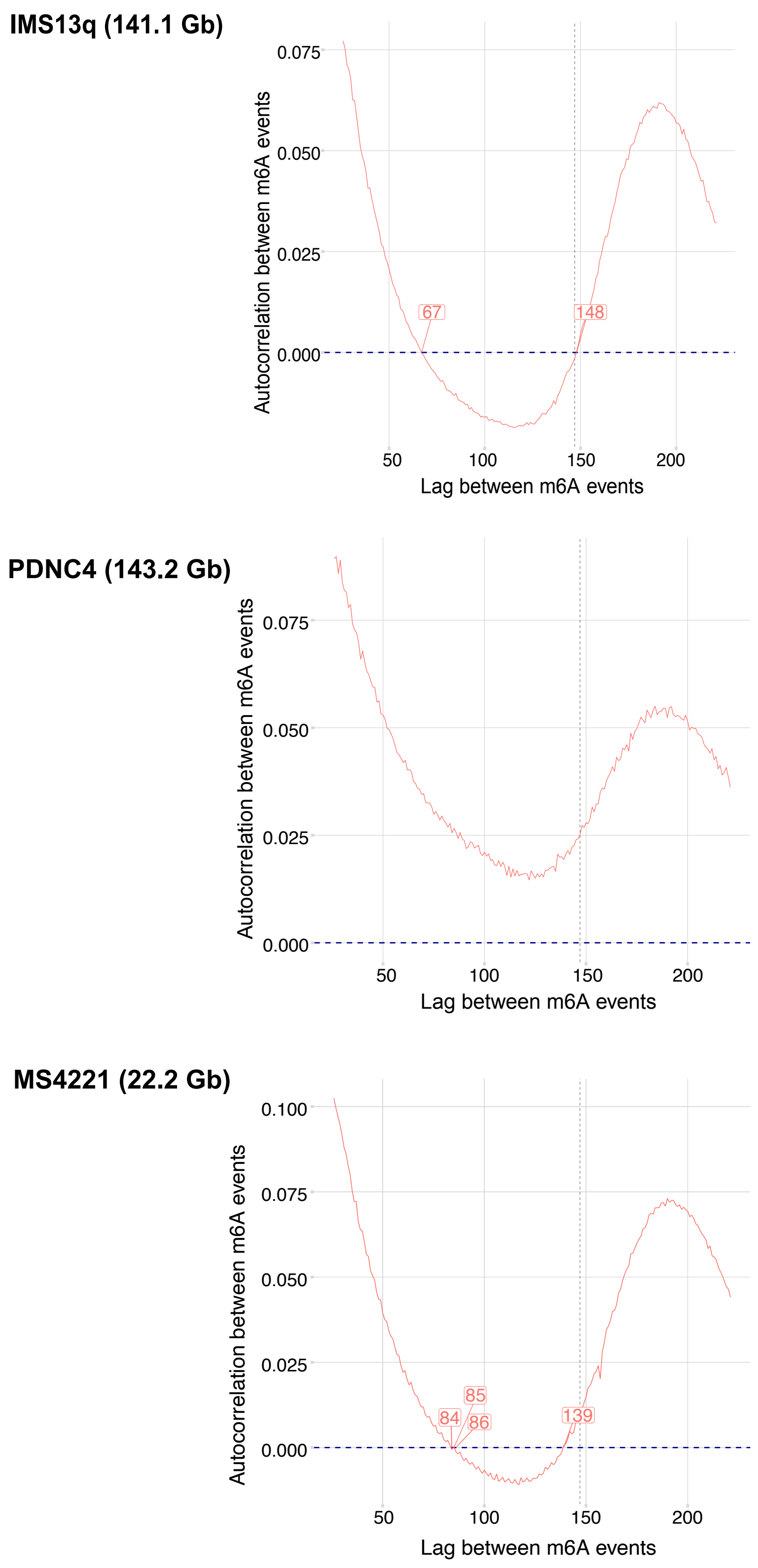

**Supplemental Figure 22. Autocorrelation and lag between m6A events in fiber-seq reads.** From top: IMS13q, PDNC4, and MS4221. Total Gb of data is shown for each line. Numbers denote the lag points where the red line crosses 0.000.

#### Supplemental Figure 23

**A.** Native centromeres

**B.** Neocentromeres

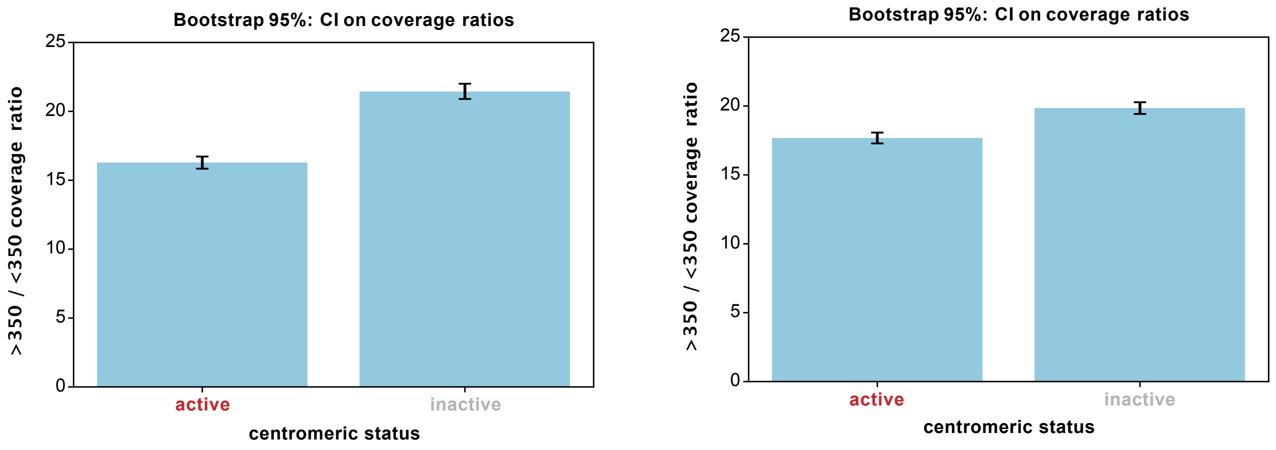

**Supplemental Figure 23. Nucleosome footprint coverage between NativeCens and NeoCens.** Barplots showing the ratio of occupancy of >350bp and <350bp footprints in active and inactive non-CDRs (CENP-A domain minus CDR(s)) of the **(A)** NativeCen and **(B)** NeoCen loci. Error bars indicate 95% confidence interval of ratio calculations from 1000x bootstrapped re-samplings.

#### Methods

##### Culturing of Human cell lines: PDNC4, MS4221, IMS13q

The two suspension lines (IMS13q and MS4221) were grown in RPMI-1640 media (Thermo Fisher Scientific) with 10% FBS, 1% L-glutamine, and 1% Pen-strep. The adherent line (PDNC4) was grown in DMEM media (Gibco) with 10% FBS, 1% L-glutamine, and 1% Pen-strep. All three cell lines were cultured in a humidity-controlled environment at 37°C with 5% $CO_2$.

##### Oxford Nanopore Technologies (ONT) sequencing of PDNC4, MS4221, IMS13q

##### PDNC4

High molecular weight (HMW) DNA was extracted from a frozen cell pellet with the Circulomics Nanobind HMW kit, sheared to 20kb with the Covaris g-TUBE (no size selection), and prepped with the Ligation sequencing kit from Oxford Nanopore Technologies (ONT) (SQK-LSK110) for a single flowcell on the PromethION. Ultra-high molecular weight (UHMW) DNA was extracted from a frozen cell pellet with the Circulomics Nanobind UHMW kit and prepped using the Ultra-Long DNA sequencing kit (SQK-ULK001). This library was run on two separate MinION flow cells. All runs were performed according to the manufacturer's protocol.

##### *MS4221 and IMS13q*

ONT data was generated from MS4221, including two whole genome runs (one HMW, one UHMW) and one adaptive sampling[1] run (HMW). HMW DNA was extracted from a frozen cell pellet with the Circulomics Nanobind HMW kit before being sheared with a Covaris G-Tube to 20kb and size selected using the Circulomics short-read eliminator XS kit to remove DNA <10kb. This sample was prepared using the Ligation sequencing kit from ONT (SQK-LSK110) and run on a single PromethION flowcell. DNA for the UHMW runs was extracted from frozen cell pellets with the Circulomics Nanobind UHMW kit, with no shearing or size selection prior to library preparation. The whole genome sample was prepared using the Ultra-Long DNA sequencing kit from ONT (SQK-ULK001) and run on the MinION across two flow cells. The other sample was prepped using the Ligation sequencing kit (SQK-LSK110) and run on the MinION using the adaptive sampling method[1], wherein the target was the NEOcen region/VNTR of chr8 +/- 1Mb from CHM13.

The remaining MS4221 data generated included both HMW and UHMW, all sequenced on the PromethION. UHMW was extracted as follows. Briefly, 5 x $10^7$ cells were lysed in a buffer containing 10 mM Tris-Cl (pH 8.0), 0.1 M EDTA (pH 8.0), 0.5% w/v SDS, and 20 ug/mL RNase A for 1 hour at 37°C.

Proteinase K (200 ug/mL) was added, and the solution was incubated at 50C for 2 hours. DNA was purified via two rounds of 25:24:1 phenol-chloroform-isoamyl alcohol extraction followed by ethanol precipitation. Precipitated DNA was solubilized in 10 mM Tris (pH 8) containing 0.02% Triton X-100 at 4°C for two days. Libraries were constructed using the Rapid Sequencing Kit (SQK-RAD004) from ONT. HMW DNA was extracted from cells using a modified Qiagen Gentra Puregene Cell Kit protocol[2]. HMW DNA libraries were prepared with the Ligation Sequencing Kit (SQK-LSK109).

Both UHMW and HMW ONT data was generated for IMS13q as follows. The UHMW DNA and libraries were generated as for MS4221 described above. The data generated included six sequencing runs: five whole genome runs (four HMW, one UHMW) and one adaptive sampling run (HMW). DNA for the five HMW runs was extracted from frozen cell pellets with either QIAGEN Genomic-tips or Circulomics Nanobind HMW kit. The four whole genome runs were sheared to 20kb with the Covaris g-TUBE and all but one were size selected with the Circulomics short-read eliminator XS kit to remove DNA <10kb. In contrast, HMW DNA for adaptive sampling was neither sheared nor size selected. All five HMW samples were prepared with the Ligation Sequencing Kit from ONT (SQK-LSK110) for runs on either the MinION or the PromethION. DNA for the UHMW run was extracted from a frozen cell pellet with the Circulomics Nanobind UHMW kit with no shearing or size selection prior to library preparation with the Ultra-Long DNA sequencing kit (SQK-ULK001) for sequencing on the MinION.

##### *ONT methods applied across all three lines*

DNA concentration was assessed with the Qubit dsDNA High Sensitivity kit and Agilent Genomic Tapestation analysis was performed to confirm HMW/UHMW integrity of all samples. Regardless of the extraction method, library kit, or sequencing machine used, all samples were run with ~25 fentamoles on R9.4 flow cells and were basecalled using the GUPPY super accuracy basecaller (either live or post-sequencing). Details of all extractions, library preparations, and sequencing runs are reported in SuppTable 1. To determine the fold-coverage and read N50 of each dataset, a custom python script, ont_stats.py (https://github.com/EichlerLab/compteam_tools/blob/main/ont_stats), was run with the following command: "ont_stats --fofn {fastq.fofn} -g 3.1". The total sequencing coverage, the coverage of >100 kbp-long-reads, and the read N50 were reported in SuppTable 2.

#### PacBio HiFi sequencing of PDNC4, MS4221, IMS13q

To generate PacBio HiFi data from the MS4221 genome, HMW DNA was extracted from cells using a modified Qiagen Gentra Puregene Cell Kit protocol as above. DNA was sheared using the Megaruptor 3 (Diagenode), and libraries were prepared using the SMRTbell Express Template Prep Kit v2 and SMRTbell Enzyme Clean Up kits (PacBio). Size selection was either performed with SageELF (Sage Science), with fractions sized 15 or 18 kbp (as determined by FEMTO Pulse (Agilent)) chosen for sequencing, or with PippinHT (Sage Science) with a 13 kbp high-pass cutoff. Libraries were sequenced on the Sequel II platform with four SMRT Cells 8M (PacBio) using Sequel II Sequencing Chemistry 2.0 or 2.2 with 2-hour pre-extension and 30-hour movies, aiming for a minimum estimated coverage of 30X in HiFi reads. Raw data was processed using the CCS algorithm (v4.2.0 or v6.2.0) with the following parameters: "--minLength 10 --maxLength 100000 --minPasses 3 --minPredictedAccuracy 0.99" (for v4.2.0) or "--all --all-kinetics --subread-fallback" (for v6.2.0) and reads with estimated quality scores >= Q20 were taken into downstream analysis.

To generate PacBio HiFi data from the IMS13q and PDNC4 genomes, HMW DNA was extracted from cells using the Monarch HMW Kit for Blood and Cells (New England Biolabs). DNA was sheared using the Megaruptor 3 (Diagenode), and libraries were prepared using the SMRTbell Prep Kit 3.0 with Barcoded Adapter plate v3.0 (PacBio). Size selection was performed with PippinHT (Sage Science) with a 17 kbp high-pass cutoff. Libraries were sequenced on the Sequel II or IIe platforms with three or four SMRT Cells 8M (PacBio) using Sequel II Sequencing Chemistry 3.2, with 2-hour pre-extension and 30-hour movies, aiming for a minimum estimated coverage of 35X in HiFi reads (assuming a genome size of 3.1 Gbp). Raw data was processed using the CCS algorithm (v6.3.0) with the following parameters: "--all --all-kinetics --subread-fallback", and demultiplexed with lima (v2.5.1) with the following parameters: "--hifi-preset SYMMETRIC-ADAPTERS --min-score 80 --min-qv 20". Reads with estimated quality scores >= Q20 were taken into downstream analysis.

#### Assembly, validation, and annotation of NativeCen and NeoCen regions in MS4221 (chr8), IMS13q (chr13), and PDNC4 (chr4)

##### *Assembly generation*

To generate whole-genome assemblies of the MS4221, IMS13q, and PDNC4 genomes, we assembled the PacBio HiFi and ONT reads using Verkko[3] (v1.0 for MS4221 and v1.1 for IMS13q and PDNC4) using the following command: "verkko -d {dir} --hifi $( cat {hifi_fastq.fofn} ) --nano $( cat {ont_fastqs.fofn} ) --sge --mbg {dir_to_mbg} --snakeopts "-j {number_of_threads}". We also assembled just the PacBio HiFi reads from each genome using hifiasm[4] (v0.16.1) using the following command: "hifiasm -o {dir_name) -t {number_of_threads} $(cat {hifi_fastq.fofn})". To determine the assembly statistics (reported in SuppTable 2), a custom python script, seq_stats.py (https://github.com/mrvollger/utilities/blob/master/seq_stats.py), was run with the following command: "seq_stats.py -r {assembly.fa}". The assemblies, per haplotype, are available on NCBI per cell line/BioSample as the following: PDNC4 (SAMN46577041), MS4221 (SAMN54265883), IMS13q (SAMN54262959).

To determine if the NativeCen and NeoCen regions were assembled, each assembly was aligned to the complete T2T-CHM13 with minimap2[5] using parameters "-t 4 -I 8G -a --eqx -x asm20 -s 5000". For PDNC4, Hifiasm assembled complete, phased chr4 NativeCens and NeoCens, while Verkko assembled chr13 phased NativeCens and NeoCens from IMS13q. However, it should be noted that genome assemblers available at the time of analysis assumed chromosomal diploidy. As such, only two NeoCen haplotypes were assembled for IMS13q, which carries three copies of 13q14.3[6,7] indicating that two of the IMS13q 13q14.3 haplotypes were merged into one assembly.

For each of the three assemblies, where both regions were completely resolved and phased (Fig. 1), some contigs were flipped in orientation compared to T2T-CHM13. These contigs were extracted with fastaexplode (https://github.com/nathanweeks/exonerate/blob/master/src/util/fastaexplode.c), reverse complemented with seqtk seq -r (https://github.com/lh3/seqtk), renamed, and then substituted in each of their respective cell-line derived assemblies.

##### *Assembly validation and repeat annotation*

To ensure there were no misassembles, the ONT (>30kb) and HiFi raw reads were mapped back to their respective assembly using Winnowmap v2.03[8] with parameters "-W meryl_k15mers.txt --eqx -ax map-ont -s 4000 -t 4 -I 8g" and plotted with NucFreq[9] across both the NativeCen and NeoCen haplotypes to assess coverage. Both read sets (ONT and Hifi) span the NeoCen regions relatively evenly without any read dropouts (SuppFig. 1). However, there are some sites with a drop in HiFi read coverage within the NativeCen, yet all have sufficient ONT read coverage across them indicating the drops in HiFi coverage are likely sequencing bias of the technology and a consequence of the read length disparity between HiFi (shorter; max 22kb) and ONT (longer; max 4Mb) reads mapped over a highly repetitive region (SuppFigs. 2-4). Additionally, due to the higher error rate in ONT reads compared to HiFi reads, the frequency of the second most common base (SuppFigs. 1-4, NucFreq plots, red) aligned is higher in ONT reads.

RepeatMasker was used for the annotation of HSat and αSats. We identified approximate boundaries of the HOR(s) based on these αSat annotations, which were further confirmed with StringDecomposer[10] and chromosome-specific αSat monomer consensus sequences. Any monomer with ≥ 90% identity was indicative of HOR-status and retained. These HSat/HOR boundaries were fed into TandemQUAST[11] for evaluation of the assembly of extra-long tandem repeats across the arrays using both read sets (ONT and Hifi). Three plots were produced from TandemQUAST: coverage, breakpoint ratio, and unique solid k-mer distribution (SuppFigs. 2-4).

##### *Assembly and validation of chr4 NativeCen in MS4221*

Both IMS13q and MS4221 Verkko assemblies were assessed for contiguous chr4 NativeCen haplotypes to include in the comparison with PDNC4 and publicly available chr4 assemblies (see below). The MS4221 Verkko assembly was the only one to have at least a single, contiguous chr4 NativeCen haplotype. This chr4 NativeCen haplotype was validated with both ONT and HiFi reads, and repeats were annotated using the same methods as described above in *Assembly validation and repeat annotation* (SuppFig. 15).

##### *HG002, T2T-CHM13, HG01114, HG00733 and HG02492 chr4 NativeCens*

Previously assembled NativeCens for chr4 from HG002[12] and T2T-CHM13[13] are available in browser: https://github.com/marbl/T2T-Browser. HPRC chr4 NativeCens HG00733 are available https://github.com/human-pangenomics/hprc_intermediate_assembly/blob/main/data_tables/assemblies_release2_v1.0.index.csv. HGSVC NativeCens for chr4 from HG01114 and HG02492 are available under the following accessions: HG01114, GCA_964198485 and PRJEB83624; HG02492, GCA_964199175 and PRJEB83624. The following annotations were included in our analyses: 1) T2T-CHM13v2.0 RepeatMasker[14], 2) HG002v1.1 ONT 5mC CpG Processed Methylation (Guppy 6.3.8)[12], 3) HG002v1.1 RepeatMasker, and 4) ONT 5mC CpG Processed Methylation (Guppy3.6.0, nanopolish) for T2T-CHM13v2.0: https://github.com/marbl/CHM13.

##### *IMS13q breakpoint resolution*

The IMS13q ONT (>1kb) reads were mapped to T2T-CHM13, which includes a normal, non-rearranged chr13, using Winnowmap v2.03 with parameters "-W meryl_k15mers.txt --eqx -ax map-ont -s 4000 -t 4 -I 8g" and viewed in the UCSC Genome Browser (SuppFig. 5). Since each cell has three copies of the qter region (one from the normal chr13, two from the inverted duplication chr13), we observed an expected increase in read coverage over this region, initiating at the breakpoint (SuppFig. 5, bottom left). With this single base breakpoint resolution, we determined the breakpoint lies in band 13q14.3 (T2T-CHM13 coordinates, chr13:51,976,458), which was further confirmed by assessment of all ONT reads (>100kb) spanning this site. These reads were extracted with seqtk subseq (https://github.com/lh3/seqtk) and assessed in Geneious v2019.1.3 revealing three different ONT read types spanning this region (SuppFig. 5). The first group of reads ("read 1" in SuppFig. 5) are those that span the breakpoint indicating they originated from the normal chr13 (no invdup). The second set of reads ("read 2") are those that contain the pter region, but do not span the breakpoint as they originated from the NativeCencarrying terminal-deletion chromosome. These reads extended from the 13pter region and were clipped at the breakpoint (13q14.3), with the remainder of the clipped reads mapping to 13q34 ~500kb from the 13q telomere, indicating a fusion occurred to stabilize the chromosome. We observed an expected increase in read coverage at 13q34 as well, making these cells tetraploid for the 13q34-qter region (SuppFig. 5, bottom right). While no reads were long enough to span this 500kb+ stretch from breakpoint (13q14.3) to 13q telomere, the longest ONT reads mapped ~100kb away from the telomeric repeats in CHM13; thus, we can assume there is a telomere at the end of the chromosome as identified with FISH[7].

The third set of reads ("read 3") are those that, like the second set, do not span the breakpoint but contain the qter region (q14.3-qter). These reads originated from the inverted duplication chr13. Further assessment of this third read set revealed that while the break was at 13q14.3, the increase in read coverage over T2T-CHM13 chr13 did not begin for another ~10kb (T2T-CHM13 coordinates, chr13:51,986,512) (SuppFig. 5, bottom left). Like the second read set, mapping to T2T-CHM13 resulted in clipping of the read at the breakpoint and a continuation of that same read at the 10kb mark. This 10kb region was found as a single copy in these reads flanked by the inverted duplicated q14.3-qter regions, suggesting that this region underwent non-allelic homologous recombination (NAHR) to repair this chromosome. YASS[15] dot plots (SuppFig. 5, bottom left) were also used to confirm this inverted duplication of chr13. Even without a full genome assembly, long-reads allowed for improved resolution of the breakpoint from 13q21.1 to 13q14.3 as well as inferring the mechanisms involved in rescue or stabilization of both abnormal chromosomes post DNA breakage.

##### *Derivation of ONT CpG DNA methylation*

FAST5s were super-accurately basecalled (≥Q10) with Bonito-Remora (using basecalling model: dna_r9.4.1_e8_sup@v3.3) for identification and tagging of the modified base, 5-methylcytosine (5mC), across reads. Reads were mapped to their respective cell line specific phased assemblies with a two-step pipeline, where step 1 was run per "{input_unaligned_bam}" and in step 2, multiple "{output_fastq}" were fed into Winnowmap v2.03 for a single "{remapped_bam}". The following code was used (requiring at least samtools v1.15.1(Danecek et al. 2021):

- Step 1) "samtools fastq -@ {threads} -T MM,ML {input_unaligned_bam} > {output_fastq}"
- STEP 2) "winnowmap -t {threads} -W {kmer_file} -ax map-ont -y {reference_file} {output_fastq} | samtools view -@ {threads} -Sb -F 256 -F 2048 - | samtools sort -@ {threads} -o {remapped_bam} –"

The -y parameter in Winnowmap ensures the retention of all tags, including the 5mC tags, that are contained within the fastq files. Reads mapped to each haplotype were viewed in IGV v2.15.4 as individual reads colored by their MM/ML tags (bam) or summarized with modbam2bed (https://github.com/epi2me-labs/modbam2bed) and visualized in IGV v2.15.4 as heatmaps tracks. Additionally, the modbam2bed file was averaged using BEDTools (v2.29.0) map -mean[16] into 1kb windows spanning the contig generated with BEDTools (v2.29.0) makewindows and subsequently viewed in IGV v2.15.4 as the smoothed appearance allowed for improved detection of broad methylation frequency changes.

##### *Identification of merged NeoCen haplotypes in IMS13q using individual methylated ONT reads*

There were three scenarios for how the haplotypes could have been merged during the IMS13q Verkko assembly process. Scenario one would be the result of the two duplicated copies (one active, one inactive) merged into one haplotype (SuppFig. 17D: b+c), resulting in one haplotype with no CDR (SuppFig. 17D: a) and the other with a “half-intensity” CDR because of averaging between active and inactive duplicated copies (SuppFig. 17D: b+c). Scenario two would be the result of the merging of a duplicated (active) and non-duplicated (inactive) copy (SuppFig. 17D: a+c), which would have the same methylation profiles as scenario one. Scenario three would be the result of the merging of the two inactive copies (SuppFig. 17D: a+b), resulting in one haplotype with no CDR (SuppFig. 17D: a+b) and the other with a CDR (SuppFig. 17D: c).

Rather than using a methylation average (i.e., a heatmap track), we assessed the methylation patterns from individual reads mapped to each haplotype from the Verkko IMS13q assembly. We did not observe any haplotype wherein 50% of reads were hypomethylated and 50% were hypermethylated (SuppFig. 17E). Thus, we determined that the two inactive copies were merged (scenario three) resulting in an unmerged active haplotype (SuppFig. 17C: c (Hap1) and a+b (Hap2).

##### *Assembly and regional based unique K-mer generation with Meryl*

Unique or single-copy 21-mers were generated with Meryl[17] in three ways.
The first was based on a given region between haplotypes within one assembly (Fig. 2, Fig. 4A-F, SuppFigs. 6, 9, 10, 19A-B), while the second was within each chr4 HOR (Fig. 3E), both of which are for assessment of sequence differences as a proxy for SNP detection. The third method was based on each cell-line specific phased assembly (PDNC4, MS4221, or IMS13q) for the purpose of filtering PRO-seq reads for increased mapping confidence (SuppFig. 21). It should be noted that since these assemblies are not at T2T-level, some unique 21-mers are likely missing and therefore, there may be more unique 21-mers reported than exist in these lines. The filtering method utilized the tool overlapSelect (GenomeBrowser/20180626) with the setting “-overlapBases=21bp”.

#### Chromatin Immunoprecipitation sequencing (ChIP-seq) data processing – Passage A (IMS13q, MS4221 and PDNC4)

This data is referred to as "Passage A" throughout the text as it was produced in a previous study published in 2013[18].

##### *Pre-processing and mapping*

Raw 100bp paired-end fastqs from[18] (GSE44724; anti-CENP-A and input DNA) were first quality trimmed (Phred score >=20) and Illumina TruSeq adapter sequences were removed using cutadapt[19]. Reads shorter than 50bp were discarded. The remaining reads were aligned to their respective cell line–specific phased assemblies using Burrows-Wheeler Aligner (BWA) with the parameters -k 50 -c 1000000. Unmapped, supplementary, and secondary alignments were subsequently filtered out using samtools view -F 2308.

##### *Post-processing*

The aligned bams were converted to bigwig format for visualization in IGV (v2.15.4) and were used for comparison against the input DNA control and mapping quality filtering using deeptools[20] bamCompare with the parameters --operation ratio --binSize 1 --minMappingQuality 1. To identify CENP-A enriched regions and/or peaks, MACS3 (Model-based Analysis of ChIP-Seq[21]) was used with input DNA as the control sample for increased specificity. Parameters were set to report broad peaks with a q-value of 0.01 or below and no more than two duplicate reads were retained to compensate for the possibility that a common repeat read was misclassified as a duplicate[22].

#### Chromatin Immunoprecipitation sequencing (ChIP-seq) for Passage B MS4221

This newly derived data is considered Passage B due to the time since the first set of ChIP-seq data[18] (Passage A) was generated.

##### *ChIP-seq procedure*

Native CENP-A ChIP-seq on MS4221 cells was performed as described previously[18,23] with some modifications. Briefly, 3-4 x $10^7$ cells were collected and resuspended in 2 mL of ice-cold buffer I (0.32 M sucrose, 15 mM Tris, pH 7.5, 15 mM NaCl, 5 mM MgCl2, 0.1 mM EGTA, and 2x Halt Protease Inhibitor Cocktail (Thermo Fisher 78429)). 2 mL of ice-cold buffer II (0.32 M sucrose, 15 mM Tris, pH 7.5, 15 mM NaCl, 5 mM MgCl2, 0.1 mM EGTA, 0.1% IGEPAL, and 2x Halt Protease Inhibitor Cocktail) was added, and samples were placed on ice for 10 min. The resulting 4 mL of nuclei were gently layered on top of 8 mL of ice-cold buffer III (1.2 M sucrose, 60 mM KCl, 15 mM, Tris pH 7.5, 15 mM NaCl, 5 mM MgCl2, 0.1 mM EGTA, and 2x Halt Protease Inhibitor Cocktail (Thermo Fisher 78429)) and centrifuged at 10,000 × g for 20 min at 4°C. Pelleted nuclei were resuspended in buffer A (0.34 M sucrose, 15 mM HEPES, pH 7.4, 15 mM NaCl, 60 mM KCl, 4 mM MgCl2, and 2x Halt Protease Inhibitor Cocktail) to 400 ng/mL. Nuclei were frozen on dry ice and stored at 80°C. MNase digestion reactions were carried out on 200-300 µg chromatin, using 0.2–0.3 U/µg MNase (Thermo Fisher #88216) in buffer A supplemented with 3 mM CaCl2 for 10 min at 37°C. The reaction was quenched with 10 mM EGTA on ice and centrifuged at 500 × g for 7 min at 4°C. The chromatin was resuspended in 10 mM EDTA and rotated at 4°C for 2 h. The mixture was adjusted to 500 mM NaCl, rotated for another 45 min at 4°C and then centrifuged at max speed (21,100 × g) for 5 min at 4°C, yielding digested chromatin in the supernatant. Chromatin was diluted to 100 ng/ml with buffer B (20 mM Tris, pH 8.0, 5 mM EDTA, 500 mM NaCl and 0.2% Tween 20) and precleared with 100 µL 50% protein G Sepharose bead (GE Healthcare) slurry for 20 min at 4°C, rotating. Precleared supernatant (10–20 µg bulk nucleosomes) was saved for further processing. To the remaining supernatant, 20 µg mouse monoclonal anti-CENP-A antibody (Enzo ADI-KAM-CC006-E) was added and rotated overnight at 4°C. Immunocomplexes were recovered by the addition of 200 mL 50% protein G Sepharose bead slurry followed by rotation at 4°C for 3 h. The beads were washed 3x with buffer B and once with buffer B without Tween. For the input fraction, an equal volume of input recovery buffer (0.6 M NaCl, 20 mM EDTA, 20 mM Tris, pH 7.5, and 1% SDS) and 1 mL of RNase A (10 mg/mL) was added, followed by incubation for one hour at 37°C. Proteinase K (100 mg/ml, Roche) was then added, and samples were incubated for another 3 h at 37°C. For the ChIP fraction, 300 µL of ChIP recovery buffer (20 mM Tris, pH 7.5, 20 mM EDTA, 0.5% SDS and 500 mg/mL Proteinase K) was added directly to the beads and incubated for 3–4 h at 56°C. The resulting Proteinase K–treated samples were subjected to a phenol-chloroform extraction followed by purification with a QIAGEN MinElute PCR purification column. Unamplified bulk nucleosomal and ChIP DNA were analyzed using an Agilent Bioanalyzer instrument and a 2100 High Sensitivity Kit.

##### *Illumina Library Preparation*

Sequencing libraries were generated using the TruSeq ChIP Library Preparation Kit - Set A (Illumina IP-202-1012) according to the manufacturer's instructions, with some modifications. Briefly, 5–10 ng bulk input nucleosomal (control) or ChIP DNA was end-repaired and A-tailed. Illumina TruSeq adaptors were ligated, libraries were size-selected to exclude polynucleosomes using an E-Gel SizeSelect II agarose gel, and the libraries were PCR-amplified using the PCR polymerase and primer cocktail provided in the kit. The resulting libraries were submitted for 150 bp, paired-end Illumina sequencing using a NextSeq 500/550 High Output Kit v2.5 (300 cycles).

##### *Pre-processing, mapping, and post-processing*

Raw 150bp paired-end fastqs (input and CENP-A) were pre-processed, mapped, and post-processed as for Passage A data.

#### CUT&RUN sequencing for Passage B PDNC4 and Passage B IMS13q

Note that while both ChIP-seq (above) and CUT&RUN were performed for Passage B PDNC4 and Passage B IMS13q, only ChIP-seq for Passage B MS4221 was successful. Numerous attempts to produce a CUT&RUN library for MS4221 failed, likely due to the slow growth and low cell count for the cell line.

##### *CUT&RUN procedure*

Live cells were processed with the Cell Signaling CUT&RUN Assay Kit #86652 with minor modifications. While approximately 250K cells were used for IMS13q, for PDNC4 we had to increase input to one million cells to achieve successful CENP-A-DNA enrichment. For both samples, the wash steps were skipped during "Live Cell Preparation" to reduce cell loss. For the positive and negative controls, we used the provided antibodies at the suggested concentrations (H3K4me3 (C42D8) Rabbit mAb #9751 and Rabbit (DA1E) mAb IgG XP® Isotype Control (CUT&RUN) #66362). For CENP-A enrichment, we used the Enzo CENP-A antibody (ADI-KAM-CC006-E) at 1:50 final concentration (added 2ul of 1:100 dilution). The chromatin of the input samples was fragmented by sonication using the Covaris S2 machine with Covaris microtube AFA Fiber Pre-slit snap cap tubes. We used the following settings to achieve a 300bp peak with a ~100-700bp smear of DNA: Intensity (5), Duty cycle (10%), Cycles/Burst (200), Treatment time (50 sec), Temp (4°C). For sample normalization, *S. cerevisiae* spike-in DNA (provided with kit) was added to each of the CENP-A-enriched samples. DNA concentration was assessed with the Qubit dsDNA High Sensitivity kit and the Agilent High Sensitivity D1000 Tapestation analysis was performed to confirm successful shearing of inputs and enrichment of CENP-A and control samples.

##### *Illumina Library Preparation*

CUT&RUN libraries were prepared with the NEBNext Ultra II DNA Library Prep Kit for Illumina (#E7645S) with minor modifications to the provided protocol. End prep consisted of 30 mins at 20°C followed by 30mins at 50°C. Adaptors were diluted to 1:15 (1.5mM working concentration) for

IMS13q and 1:10 (1mM working concentration) for PDNC4. Cleanup of adaptor-ligated DNA without size selection was performed with 1.1X AMPure XP beads, per the CUT&RUN protocol. NEBNext indexed primers were used, and PCR amplification was run in a thermal cycler according to the library prep protocol, with the following modifications: 1) number of cycles was set to 15 based on the CUT&RUN DNA yield (1-5ng), and 2) anneal and extension time was reduced from 75 sec to 13 sec to exclude amplification of large library DNA fragments (>1,000 bp). PCR cleanup was performed with 1.0X AMPure XP beads and eluted in 33ul of 0.1X TE. Final DNA libraries were assessed with the Agilent High Sensitivity D1000 Tapestation and then sequenced on an Illumina NovaSeq, producing paired-end 150bp reads.

###### *Pre-processing, mapping, and post-processing*

To assess positive and negative control CUT&RUN samples for each line, the raw 150bp paired-end fastqs were pre-processed, mapped, and post-processed the same way as the paired-end ChIP-seq reads (above). The CENP-A enriched fastqs were mapped slightly differently. Since *S. cerevisiae* spike-ins were included in the CENP-A enriched samples, the trimmed reads were mapped to a combined dual-genome for equal mapping competition. These dual genomes included: 1) their respective cell line specific phased assembly, and 2) the *S. cerevisiae* genome. Reads mapped to the *S.cerevisiae* genome were filtered out prior to post-processing. To identify CENP-A enriched regions or peaks, SEACR (Sparse Enrichment Analysis for CUT&RUN[24]) was used to report the top 1% of peaks based on the total signal within the peaks using a stringent threshold (the threshold at which the maximum percentage of target versus IgG signal blocks are retained, as per[24]). This data is referred to as "Passage B" throughout the text as it was generated with cells that had undergone many more passages since the original ChIP-seq data was produced[18].

##### Directed Methylation and Long-read sequencing (DiMeLo-seq)

###### *Experimentation, library prep, & sequencing*

To perform DiMeLo-seq for Passage B PDNC4 (SuppFigs. 10,17), we followed protocols according to DiMeLo-seq: Directed Methylation with Long-read sequencing V.2[25,26]. In brief, 10M cells were pelleted and washed in PBS, then resuspended in Wash Buffer (20mM HEPES-KOH pH 7.5, 150 mM NaCl, 0.5mM spermidine, 0.1% BSA, and protease inhibitor cocktail (CellSignaling #7012) with 0.02% digitonin (CellSignaling #16359). The samples were incubated on ice for 5 minutes to lyse and permeabilize cells, then split evenly between anti-CENP-A and no-antibody control samples. The samples were pelleted at 500xg for 3 minutes, then resuspended in Wash Buffer with 0.1% Tween-20 and either

1:50 CENP-A monoclonal antibody (3-19) (Enzo, ADI-KAM-CC006) or no antibody. The samples were rotated at 4°C for 2 hours, then spun at 500xg for 3 minutes. The pellets were washed twice with Wash Buffer containing 0.1% Tween-20; between each wash, samples were rotated at 4°C for 5 minutes. The samples were incubated on ice for 5 minutes to lyse and permeabilize cells, then split evenly between anti-CENP-A and no-antibody control samples. The samples were pelleted at 500xg for 3 minutes, then resuspended in Wash Buffer with 0.1% Tween-20 and either 1:50 CENP-A monoclonal antibody (3-19) (Enzo, ADI-KAM-CC006) or no antibody. The samples were rotated at 4°C for 2 hours, then spun at 500xg for 3 minutes. The pellets were washed twice with Wash Buffer containing 0.1% Tween-20; between each wash, samples were rotated at 4°C for 5 minutes.

To bind pA-Hia5, the pellets were gently resolved in 200 nM pA-Hia5 and rotated at 4°C for 2 hours, then the above-described wash steps were repeated for a total of two washes. To activate Hia5 MTase activity, the pellets were resolved in Activation Buffer (15mM Tris pH 8.0, 15 mM NaCl, 60 mM KCl, 1mM EDTA pH 8.0, 0.5 mM EGTA pH 8.0, 0.05 mM spermidine, 0.1% BSA, and 800 uM S-adenosylmethionine (SAM)). The samples were incubated for 2 hours at 37°C, mixing every 30 minutes and replenishing 800 uM SAM after 1 hour. Following incubation with SAM, the samples were spun at 500xg for 3 minutes and resuspended in 100 uL cold PBS before proceeding to DNA extraction with the Monarch HMW DNA Extraction Kit for Cells & Blood (T3050L).

To perform ONT sequencing, samples were sheared using the Megaruptor 3 (Diagenode) targeting 20-40kb and analyzed for size using the Femto Pulse system (Agilent). Samples were size-selected using the Short Read Eliminator (SRE) XS kit (Pacific Biosciences). Libraries were prepared using the LSK114 kit from ONT and sequencing on a R10.4.1 flow cell on the PromethION system.

##### *Data processing*

First, CENP-A and no-Hia5 controls were independently basecalled with Dorado (v0.8.2), using the "sup,6mA" algorithm and filtering for a minimum q score of 10. Reads were subsequently aligned to the reference assembly using Dorado (v0.8.2) and processed using samtools (v1.19.2). Alignment files were converted to methylated bed files using modbam2bed with the flag "--mod_threshold=0.8" to filter low confidence calls. Files were converted to bedgraphs using bedtools (v2.31.1), then the no-Hia5 control signal was subtracted from the CENP-A signal using awk commands. Bigwigs were generated using the bedGraphtoBigWig UCSC Genome Browser module. Full details of data processing are available in: https://github.com/gabriellehartley/cenpa-dimelo-processing.

#### Chromatin accessibility assessment using Fiber-seq via Pacbio HiFi reads

##### *Fiber-seq experimentation*

To perform Fiber-seq on IMS13q, MS4221, and PDNC4, ~3M cells were collected and processed according to the methods in[27]. DNA was extracted using the Monarch HMW DNA Extraction kit.

##### *Library prep & sequencing with PacBio*

To obtain HiFi reads, Fiber-seq DNA was sheared using the Diagenode Megaruptor® 3 system at speed 32, targeting 18kb fragment lengths. The DNA was size selected using the Pacific Biosciences Short Read Eliminator (SRE) XS kit (102-208-200). Libraries were created per manufacturer's instructions using the SMRTbell® prep kit 3.0 (102-141-700) and sequenced on either the Sequel IIE (for the 2023 libraries) or the Revio (for the 2025 libraries) (SuppTable 1).

##### *Data processing*

HiFi reads were processed with jasmine using the --keep-kinetics flag to maintain methylation information, then aligned to respective assemblies using pbmm2. 6mA calls were predicted using fibertools-rs, assessed with fiberseq-qc (SuppFig. 22) and then passed into the FIRE pipeline[28] to call inferred regulatory elements (FIREs). All scripts and/or tools are included in the Fiber-seq repository and were run without modification (https://github.com/fiberseq).

##### *Fiber-seq Footprinting analyses*

Fiber-seq footprints (Fig. 6D-F) were called using FiberHMM v1.4, using transition probabilities trained on a subset of the datasets. Footprints found at the edge of reads were filtered due to their ambiguous size. To compare differences in the distribution of footprint enrichment between active and inactive regions, counts of unique footprint sizes between 80bp and 450bp were normalized, with this process repeated after 10,000x bootstrapped resampling of footprints identified in active and inactive regions to identify 95th percentile ranges. To test the relative enrichment of 1x CENP-A and 2x CENP-A sized footprints in different regions, all footprints identified for active or inactive CDRs and CENP-A regions (excluding the CDR) in NativeCen and NeoCen were bootstrap resampled 10,000x and the relative enrichment of different size-ranges of footprints were plotted with error bars representing the 95th percentile of the resampled ratio of enrichment.

#### Precision Run-on sequencing (PRO-seq) for PDNC4, MS4221, and IMS13q

##### *Cell permeabilization*

For PDNC4, adherent cells were washed 2x in cold 1x PBS before adding 5mL of buffer P (10mM Tris-Cl pH 8.0, 10mM KCl, 250mM sucrose, 5mM MgCl2, 1mM EDTA, 0.05% Tween-20, 0.5mM DTT, 10% Glycerol). Cells were scraped, collected, and 10uL was removed for cell counting while the remainder was centrifuged at 1000xg for 5 min. For MS4221 and IMS13q, an aliquot of suspension cells was counted, and the remainder was centrifuged along with PDNC4. 1mL of buffer W (10mM Tris-Cl pH 8.0, 10mM KCl, 250mM sucrose, 5mM MgCl2, 1mM EDTA, 0.5mM DTT, 10% glycerol) was used to gently resuspend cell pellets, before adding an additional 9mL of buffer P, inverting and centrifugation at 1000xg for 5 min. 500uL of buffer F (50mM Tris-CL pH 8.0, 40% glycerol, 5mM MgCl2, 0.1 mM EDTA, 0.5mM DTT) plus 0.5uL of RNase-inhibitor (SuperAse from Thermo Fisher Scientific) was used to resuspend the cell pellets, followed by another 500uL to wash the tubes; both were pooled together (1mL total), transferred to a 1.5mL tube, and centrifuged at 1000xg for 5 min. Finally, permeabilized cells were resuspended in 57µL of buffer F with 1µL of RNase-inhibitor added before snap-freezing in liquid nitrogen and storage at -80°C.

##### *Illumina library preparation*

PRO-seq libraries were prepared as previously described[14,29] with minor modifications. Approximately 2 x $10^6$ permeabilized cells were mixed with permeabilized *Drosophila melanogaster* (Dm) S2 nuclei in all 4-biotin-NTP run-ons (5 x $10^4$ Dm nuclei in each sample); run-on RNA was extracted with Norgen columns and eluted in 50uL $H_20$. Base-hydrolysis included incubation in 25µL cold 1N NaOH for 10min on ice, followed by the addition of 125uL cold 1M TrisCl pH 6.8, a gentle vortex, and brief spin down before enrichment with streptavidin-beads. Following 3'-ligation and the second bead binding, both end-repair reactions and the 5'-ligation were all performed with nascent RNAs still bound to the beads. These on-bead reactions were performed in a total volume of 20µL with constant rotation before elution from the beads and the subsequent reverse transcription and PCR steps. Test amplifications were performed on 5% of the library and samples were amplified to the ideal number of cycles for final preparation. Following final amplification, libraries were PAGE purified to remove adapter- dimers and select molecules between 140-650bp in size. Libraries were then sequenced on an Illumina NextSeq 550, producing paired-end, 75bp reads.

##### *Pre-processing and mapping*

Raw fastq files were first quality trimmed (Phred score >=20) and adapter sequences removed using cutadapt[19]. Reads below 20nt were removed and remaining reads were reverse complemented using the fastx-toolkit[30]. Dm S2 spike-in reads were removed by aligning reads to the Dm6 genome with bowtie2[31] using "--very-sensitive" options. Remaining reads were then aligned to their respective cell line specific phased assembly with bowtie2 using the default (end-to-end alignment, "best match") option. Sorted bam files were converted to bed files with BEDTools (v2.29.0)[16], which were subjected to one or more of the following: 1) single copy 21-mer filtering (overlapSelect -overlapBases=21) or, 2) conversion into BigWig files for data visualization in IGV (GenomeBrowser/20180626).

#### Total RNA-seq for PDNC4, MS4221, and IMS13q

##### *RNA extraction and Illumina library preparation*

Total RNA was extracted from cells of all three NeoCen lines using the mirVana Total RNA Isolation kit (Thermo Fisher Scientific). RNA quality was assessed with the Agilent Bioanalyzer Total RNA Pico assay, Ribo-depleted, and subsequently processed with the Illumina TruSeq stranded total RNA library preparation kit. Libraries were sequenced on an Illumina HiSeq 2500, producing 100bp paired-end reads.

##### *Pre-processing and mapping*

Raw, 100bp paired-end fastqs were pre-processed, mapped, and post-processed the same way as the paired-end ChIP-seq reads.

#### Gene annotation

Gene annotations were carried over from the T2T-CHM13v2.0 NCBI RefSeq curated gene set (chm13v2.0_RefSeq_Liftoff_v4.gff3; https://github.com/marbl/CHM13) to each assembly with LiftOff[32] with haplotypes run separately to ensure haploid (CHM13) to haploid annotation. Ensembl was used for identification of the main functional isoform per protein-coding gene included in this study based on APPRIS predictions[33]. Genes for PDNC4 and IMS13q are listed in SuppTable 10.

#### αSat subclassifications and Structural Variant (SV) detection

Methods were followed as per[34] for subclassification of αSats across NativeCen contigs per cell line assembled herein. αSats were broken down into one of 20 suprachromosomal families (SF#)(reviewed in [35]). Scripts are provided here: (https://github.com/fedorrik/HumAS-HMMER_for_AnVIL) using version AS-SFs-hmmer3.0.290621. Since SF classification can be used to infer age, we grouped SFs into one of three broad HOR age groups[34]: HOR (new), dHOR (old), and mon (ancient). Resulting bed files (including color codes) were converted into bigBed format and visualized in IGV v2.15.4. A colored coded key is included in SuppTable 5. HOR array loci were defined using the broad HOR annotations to include those regions that were classified as either HOR or dHOR, but not mon, as denoted in SuppTable 5. This pertains to the following figures: SuppFigs. 7-8, 11-14.

To generate the αSat HOR monomers tracks (Fig. 3A, C, D; SuppFigs. 6, 9, 10), regions corresponding to native centromeres were first identified in whole-genome assemblies using the Centromere Mapping and Annotation Pipeline (CenMAP; v0.4.3.1; https://github.com/logsdon-lab/CenMAP[36]) and annotated using the Snakemake-HumAS-SD workflow (https://github.com/logsdon-lab/Snakemake-HumAS-SD). This workflow annotates sequences using chromosome-specific αSat monomer HMM profiles from the HumAS-HMMER library and performs string decomposition via StringDecomposer[10]. αSat \HOR and dHOR\ structures were identified using an enhanced method based on the StV approach (https://github.com/fedorrik/stv), which segments arrays based on inter-monomer distance, strand orientation, and monomer order. Active, or "live", αSat HOR arrays were then defined using CenStats (v0.0.10; https://github.com/logsdon-lab/CenStats). Contiguous monomers were grouped based on proximity and consistent strand orientation, with adjacent blocks separated by less than 8 kbp iteratively merged to account for minor interruptions. Resulting arrays were filtered to retain those with at least five αSat HOR monomers and composed of ≥90% active sequence. The coordinates, lengths, and orientations of filtered arrays were output in BED format. These "live" αSat HOR array coordinates are included in SuppTable. 5 (listed as ""live" HOR (AS-HOR based)") for all three neocentromere assemblies (PDNC4, MS4221, and IMS13q), as well as the additional chr4 assemblies included in Fig. 3.

Centromeric annotations were visualized using CenPlot (v0.1.4; https://github.com/logsdon-lab/CenPlot). Annotation tracks included RepeatMasker output, α-satellite HOR structures, active HOR array strand orientation, CpG methylation frequencies inferred with CDR-Finder (v1.0.4) and 1D local self-sequence identity tracks generated using ModDotPlot (v0.8.4[37]; see below in "Sequence Identity Heat Maps" for expanded methods). Tracks were formatted as BED files and displayed as ordered layers according to a configuration file.

HOR Structural Variants (SV) were determined using this script (https://github.com/fedorrik/stv_chm13), to detect HOR monomeric indels. These annotations (SuppTable 7) are HOR-specific[34]. HOR-SV composition was determined by calculating the coverage (bp) of each SV across the HOR and dividing it by overall size of the HOR. For chr13, we identified one dominant SV, S2C13/21H1L.11-1, shared among both IMS13q NativeCen haplotypes and T2T-CHM13 NativeCen, and other SVs with varying abundance (SuppFig. 7A, SuppTable 7). For chr8, there were three SVs that made up >97% of each HOR in T2T-CHM13 NativeCen and both MS4221 NativeCen haplotypes, regardless of centromeric activity (SuppFig. 11A, SuppTable 7). Lastly, for chr4, we found that the D4Z1 arrays of PDNC4 and T2T-CHM13 share a dominance of one SV (S2C4H1L.1-19) while the D4Z1B arrays which lack this SV completely (SuppFig. 12A, SuppTable 7). Tracks for each neocentromere assembly are included in SuppFigs. 6, 9, 10.

##### Regional boundary demarcation (CENP-A domains, flanks) across NeoCens and NativeCens

For PDNC4 and IMS13q, CENP-A domain boundaries were demarcated by CUT&RUN peaks from Passage B called with SEACR[24] (top 1%). For MS4221, CENP-A domain boundaries were demarcated by ChIP-seq peaks from Passage B called with MACS3[21] (using input DNA control). Across all three lines, flanks were demarcated by extending 100kb from the CENP-A domain on either side. However, with the realization that there could be indels in one haplotype, but not the other, these 100kb flank boundaries were subsequently adjusted to end on the same repeat between haplotypes for the most accurate comparisons. The sequences of these regions were isolated from each assembly and used in subsequent analyses. Coordinates and corresponding statistics are included in SuppTable 8.

##### CDR annotations

CDRs in each active NativeCen and NeoCen were annotated as described in . The CDRs in the active neocentromeres (PDNC4 and IMS13q) were mapped with minimap2[5] to their inactive centromeric counterpart and further confirmed using TE annotations as markers to determine corresponding boundaries for subsequent comparisons and analyses. In MS4221 (and NativeCens – as an approximation), where the centromeres fall within a tandem repeat array, we performed a manual annotation to identify a region of 1) equivalent size to the active CDR, and 2) representative of the methylation status and repeat content of the inactive centromere – made possible by the homogeneity of tandem repeat arrays. CDR loci for and stats are provided in SuppTable 4 for PDNC4, MS4221, and IMS13q. CDRs were annotated for the HPRC and HGSVC assemblies using CDR-Finder (v1.0.4[38]) as a part of the cenMAP pipeline (v0.4.3.1[36]; https://github.com/logsdon-lab/CenMAP).

##### Regional GC content and CENP-B box detection

CENP-A domains (see “Regional boundary demarcation”) per NeoCen and NativeCen were run through QUAST  to determine overall GC content, and hence AT content (SuppTable 8). CENP-B boxes are denoted by a 17-bp motif that is recognized by the CENP-B DNA-binding protein.  To detect the location of these CENP-B boxes across the NeoCen and NativeCen loci, FIMO (Find Individual Motif Occurrences) was utilized along with the CENP-B box motif in MEME file format. The resulting .tsv was converted into a .bed file for visualization in IGV v2.15.4. The number of CENP-B motifs were counted per CENP-A domain and reported in SuppFig. 20B.

**Sequence Identity Heat Maps**

To generate pairwise dot plots for sequence identity within and between centromeric haplotypes we used StainedGlass (v0.5)[40]. Input sequence was fragmented into 2kb windows (default) prior to alignment with minimap2 and subsequent visualization of the dot plot and corresponding sequence identity histogram. If more than one sequence was provided, plots were generated on individual color scales, as well as a shared color-scale, ideal for comparisons.

To generate pairwise dot plots for sequence variation between centromeric haplotypes (SuppFig 19A-B), we used an earlier version of StainedGlass (v0.5). In this case, the input sequences were fragmented into 5kb windows with a 2.5kb sliding window prior to alignment and subsequent visualization. Although these plots show variation between two sequences, the dots are still colored by percent sequence identity corresponding to the provided histogram.

To generate 1D pairwise sequence identity tracks of each centromeric region (Fig. 3A, C, D), we ran ModDotPlot (v0.8.4) as part of the CenMAP workflow. 1D pairwise sequence identity tracks were generated from BEDPE files using the `localselfident` option in the CenMAP YAML configuration. Sequence identity values were binned and recolored to normalize the color scale along each centromere.

**Single Nucleotide Polymorphism (SNP) detection in NeoCens**

To identify SNPs of the NeoCen region on Hap1 (and its homologous region on Hap2) of PDNC4 and IMS13q, a minimap2[5] alignment was generated between the two haplotypes and fed into SYRI (Synteny and Rearrangement Identifier) for counting. The number of SNPs were calculated per 1kb window across the NeoCen regions using BEDTools (v2.29.0) map -o count. A summary and individual SNP loci are listed in SuppTable 11.

**TandemAligner and corresponding dot plots**

Dot plots were generated between HORs of opposing haplotypes using TandemAligner and MUMmer for identification of structural differences in combination with αSat subclassifications (SuppFigs. 7, 11, 12), as well as between the NeoCen VNTRs of MS4221 (SuppFig. 16). Self-alignments were also generated using this method to define intra-array structure of the HORs for detection of high identity regions, indicating recent expansions (SuppFigs. 8, 13, 14). The color code for the dot plots are as follows: black = MEM (maximum exact match), red = MUM (maximum unique match; occurs exactly once in both sequences), blue = alignment match, light green = alignment indel or mismatch. The blue and/or light green line indicated how two sequences are aligned; if the lines are not on the diagonal (as in a self-alignment), this indicates the presence of an indel in one or the other sequence.

#### Repeat assessment

Regions denoted in "Regional boundary demarcation" were isolated from the RepeatMasker output and summarized with the script, buildSummary.pl using a corresponding genome file indicating the region's length. Base pair composition, rather than counts, was used to summarize by repeat class, family, or subfamily. Tandem or non-interspersed repeats include satellites, simple repeats, low complexity repeats, and RNAs. Interspersed repeats include all other repeat classes (TEs and CHM13-derived composite subunits[14]. Repeat class statistics included in SuppTable 13. Non-repetitive regions were identified by subtracting the repeats (as defined by RepeatMasker) from the overall region using BEDTools (v2.29.0) subtract[16].

#### Quantification of CpG methylation frequencies and transcriptional activity

BEDTools (v2.29.0) map -mean[16] was used to average the CpG methylation per region (i.e., CDR), repeat (as defined by RepeatMasker), or non-repeat (Fig. 4G, SuppFig. 18, SuppTables 4, 8). For assessment of gene expression, BEDTools (v2.29.0) coverage -counts was used to count the number of reads mapped with Bowtie2[31] default from single-end PRO-seq or paired-end RNA-seq (frags.pe.bed) data that overlap a given gene exon. We required that at least 60% of the read must overlap the element to be counted (-F 0.6). Counts for each gene of interest for PDNC4 and IMS13q were loaded into RStudio (v.2025.05.0). Cell line and sequencing type (PRO-seq or RNA-seq) were converted to factor for handling differential analysis. The DESeq2 (v.1.42.1) function was then applied to perform normalization and estimate dispersion values, using a fitType of 'mean' for dispersion estimation. A hierarchical clustering heatmap of the regularized log-transformed count data was generated using pheatmap(v.1.0.13)(Fig. 6A,B).

#### 1KGP phased variant calls against T2T-CHM13 and associated analyses

1000 Genomes Project (1KGP) variants across the 2504 unrelated individuals compared to CHM13v1.0 were phased with SHAPEIT-duohmm v2.r904 following the filtering and phasing pipeline performed by the New York Genome Center (NYGC) for 1KGP analysis. These variant calls were subsequently filtered to biallelic variants with no missing genotype calls, and MAF >= 0.01 using bcftools. Filtered variant calls were lifted to T2T-CHM13v2.0 for visualization in the UCSC genome browser alongside the recombination map. The sex-averaged recombination map and the recombination rate annotation (cM/Mb) was lifted from GRCh38 to T2T-CHM13v2.0 (Fig. 4H-I).

Heterozygous variants were quantified per sample in 10kb windows for both chr4 and chr13 (SuppFig. 19C-D). 10kb windows were first generated with BEDTools makewindows. Then, het variants were counted within these windows using vcftools --het with the filtered phased variants calls described above. To generate distributions of per-sample het calls, the mean number of heterozygous variants across all samples was calculated within each window. Given the haplotype structure apparent around the chr13 NeoCen locus, we were curious if there were fewer samples heterozygous for the two haplotypes than expected given the allele frequency distribution of those haplotypes. To quantify this, we selected two variants (chr13:96262531:A:G and chr13:96309388:G:C) that appeared to differentiate the two haplotypes and investigated deviations from Hardy-Weinberg equilibrium within each of the 26 1KGP populations for each of the two variants (SuppFig. 19E). For each population, the subsequent samples were extracted. The expected number of heterozygous samples was calculated as 2(p)(1-p)N where p is the allele frequency of the variant within that population and N is the number of samples in the population. A chi-squared test was run with chisq = sum((obs het - exp. het)^2 / exp. het) and 1 degree of freedom (SuppTable 12).

#### Visualization of assembly-specific tracks and plots

The UCSC Genome Browser was used for track hub development for each cell line-specific assembly and visualization of read density graphs (i.e., ChIP-seq, CUT&RUN, PRO-seq, RNA-seq; bigwig format) and individual read alignments (.bam format). IGV v2.15.4 was used for visualization of colored annotation tracks (.bed format) and all CpG methylation related data. GraphPad Prism v9.1.0 was used to generate plots in Fig. 4G and SuppFigs. 7A, 11A, 12A, 18, 20 and perform unpaired *t* tests to determine significant differences in methylation (Fig. 4G and SuppFig. 18B-C).

To evaluate the k-mer diversity across different sites in the NativeCens (CDR, D4Z1, or D4Z1B), two independent analyses were performed. Analysis 1 (Fig. 3E; left panel) included D4Z1-active alleles (n=7) and analysis 2 (Fig. 3E; right panel) included D4Z1B-active alleles (n=2). For both analyses, a linear mixed-effects model was fit using the lme4 package where k-mer diversity (dependent), site (independent), and cell line (random) were included, and a maximum likelihood estimation was used to fit the model. Hypothesis testing for the fixed effect of site was performed using a Type I ANOVA

(RStudio). Post-hoc comparisons were performed using emmeans(v.1.11.1) between sites. For analysis 1, p-values for pairwise comparisons were adjusted using the Tukey method. Due to the low sample size (n=2) of analysis 2 - resulting in low statistical power - a p-value adjustment was not used, and results should be interpreted with caution. GraphPad Prism v10.4.1 was used to generate the plot in Fig. 3E.
